# A genetic basis for cancer sex differences revealed in Xp11 translocation renal cell carcinoma

**DOI:** 10.1101/2023.08.04.552029

**Authors:** Mingkee Achom, Ananthan Sadagopan, Chunyang Bao, Fiona McBride, Qingru Xu, Prathyusha Konda, Richard W. Tourdot, Jiao Li, Maria Nakhoul, Daniel S. Gallant, Usman Ali Ahmed, Jillian O’Toole, Dory Freeman, Gwo-Shu Mary Lee, Jonathan L. Hecht, Eric C. Kauffman, David J Einstein, Toni K. Choueiri, Cheng-Zhong Zhang, Srinivas R. Viswanathan

**Affiliations:** Department of Medical Oncology, Dana-Farber Cancer Institute; Boston, MA, USA; Department of Data Science, Dana-Farber Cancer Institute; Boston, MA, USA; Department of Medicine, Harvard Medical School; Boston, MA, USA; Department of Pathology, Brigham and Women’s Hospital; Boston, MA, USA; Cancer Program, Broad Institute of MIT and Harvard; Cambridge, MA, USA; Department of Biomedical Informatics, Blavatnik Institute, Harvard Medical School; Boston, MA, USA; Department of Informatics & Analytics, Dana-Farber Cancer Institute; Boston, MA, USA; Department of Pathology, Beth Israel Deaconess Medical Center; Boston, MA, USA; Department of Urology, Roswell Park Comprehensive Cancer Center; Buffalo, Nesw York, USA; Division of Medical Oncology, Beth Israel Deaconess Medical Center; Boston, MA, USA; Department of Medicine, Brigham and Women’s Hospital; Boston, MA, USA

## Abstract

Xp11 translocation renal cell carcinoma (tRCC) is a female-predominant kidney cancer driven by translocations between the *TFE3* gene on chromosome Xp11.2 and partner genes located on either chrX or on autosomes. The rearrangement processes that underlie *TFE3* fusions, and whether they are linked to the female sex bias of this cancer, are largely unexplored. Moreover, whether oncogenic *TFE3* fusions arise from both the active and inactive X chromosomes in females remains unknown. Here we address these questions by haplotype-specific analyses of whole-genome sequences of 29 tRCC samples from 15 patients and by re-analysis of 145 published tRCC whole-exome sequences. We show that *TFE3* fusions universally arise as reciprocal translocations with minimal DNA loss or insertion at paired break ends. Strikingly, we observe a near exact 2:1 female:male ratio in *TFE3* fusions arising via X:autosomal translocation (but not via X inversion), which accounts for the female predominance of tRCC. This 2:1 ratio is at least partially attributable to oncogenic fusions involving the inactive X chromosome and is accompanied by partial re-activation of silenced chrX genes on the rearranged chromosome. Our results highlight how somatic alterations involving the X chromosome place unique constraints on tumor initiation and exemplify how genetic rearrangements of the sex chromosomes can underlie cancer sex differences.

## Introduction

Xp11 translocation renal cell carcinoma (tRCC) (*1*) is a rare, aggressive, and female-predominant subtype of kidney cancer defined by an oncogenic rearrangement of the *TFE3* gene on the short arm of chromosome X (chrXp). tRCC comprises approximately 1-5% of RCCs in adults and 20% to 75% of pediatric RCCs (*2*). There are currently no effective therapies for tRCC (*3–6*), owing in part to an incomplete understanding of its biology and how it differs from more common types of kidney cancer.

tRCC is clinically and biologically distinct from other kidney cancers. Demographically, it shows a ∼2:1 female predominance in incidence, which is in stark contrast to the male predominance observed in most other subtypes of kidney cancer (*1*, *7*). Biologically, the defining genetic lesion in tRCC is an activating fusion involving *TFE3*, a transcription factor in the *MiT/TFE* family. *TFE3* fusions typically occur in-frame with a partner gene that may be encoded on either chrX or on an autosome (*8*). Although fusions involving other *MiT/TFE* family members (*TFEB* and *MITF*) can also drive kidney cancer, these are much rarer than *TFE3* fusions and likely comprise clinically distinct entities (*9*). Additionally, recent genomic studies have found few recurrent alterations in tRCC apart from the *TFE3* fusion, suggesting that it may function as the sole driver in many cases (*7*, *10–15*).

Besides tRCC being an unmet medical need, it also represents a unique model in which to address fundamental questions about tumor initiation, owing to the location of the *TFE3* gene on chromosome X. For example, genomic studies of tRCC can inform us about the mechanisms underlying the development of oncogenic fusions. Although it has been conventionally accepted that oncogenic fusions usually arise by simple, reciprocal translocations, whole-genome sequencing has shown that *bona fide* reciprocal translocations are rare in human cancers, particularly in solid tumors. Instead, many oncogenic fusions in carcinomas arise from complex events such as chromothripsis or chromoplexy (*16–18*). For example, a recent WGS study revealed that the *EWSR1-ETS* fusions driving Ewing sarcoma arise via chromoplexy rather than simple reciprocal translocations in >40% of cases (*16*). The *TMPRSS2-ERG* fusion in prostate cancer also frequently arises via chromoplexy (*17*). The *EML4-ALK* fusion in lung cancer can arise through diverse mechanisms including simple inversion, chromothripsis, and other complex events (*19*, *20*). A recent large lung cancer WGS study revealed that 74% of known fusion oncogenes arise via complex genomic events such as chromothripsis or chromoplexy (*18*). Translocations that occur as part of complex events can also underlie oncogene amplification (*21*). While *BCR-ABL1*, the fusion that drives chronic myeloid leukemia (CML) is often cited as a classical reciprocal fusion, it was recognized early on that *BCR-ABL1* fusions are often accompanied by complex alterations to one or both of the chromosomes contributing fusion partners (chr9 and chr22) and/or to additional chromosomes (*22–26*). In the specific case of tRCC, prior genomic studies have employed panel or whole exome sequencing (WES). These platforms are not of suitable resolution to fully characterize the structural variants that lie at the origins of this fusion-driven cancer. The genomic mechanisms that underpin *TFE3* fusions in tRCC therefore remain uncharacterized.

A second mechanistic question specific to tRCC is whether oncogenic *TFE3* fusions can be generated from the inactive X chromosome (chrXi) in females, thus providing a genetic explanation for the female sex bias in tRCC incidence (*1*). The X chromosome is unique amongst chromosomes in that baseline composition varies between the sexes, with females having two copies and males having one. In females, the two chrX homologs have markedly distinct chromatin states and transcriptional profiles despite nearly identical sequences. The active X (chrXa) is transcribed at a similar level as chrX in males, whereas the chrXi undergoes epigenetic silencing during early development, thereby achieving dosage compensation of chrX genes between males and females (*27*). Thus, somatic alterations involving chrX may have differential consequences if they involve chrXa vs. chrXi. In the case of tRCC, this also implies that fusions involving *TFE3* on chrXi could result in a partial reversal of XCI, a process that is typically thought to be initiated during early female development and stably maintained thereafter. As *TFE3* is located on chrX, the genomic analysis of this cancer also requires sex-specific considerations that have not been uniformly applied in prior cancer genomic studies (*28*, *29*).

By leveraging tRCC as a prototypical sex-biased cancer driven by an oncogene on chrX, we sought to define the origin and consequences of genomic events to the X chromosome during cancer initiation and evolution.

## Results

### Landscape of genomic alterations in tRCC from whole-genome sequencing

We performed WGS on 29 tRCC tumor specimens (10 primary, 19 metastases) and matched germline controls, collected from 15 distinct individuals (5 male and 10 female). The cohort included samples derived from both treatment-naïve patients (11 samples) and patients who received systemic therapy for tRCC prior to sample collection (18 samples). In one case (TRCC12), tumor tissue was sampled from the same metastatic site prior to, and following progression on, systemic therapy with a tyrosine kinase inhibitor (lenvatinib) and mammalian target of rapamycin (mTOR) inhibitor (everolimus). In a second case (TRCC18), primary tumor tissue was collected prior to the initiation of systemic therapy with a tyrosine kinase inhibitor (cabozantinib) and immunotherapy (nivolumab), and 13 metastatic sites along with 2 adjacent normal tissue samples were collected upon rapid autopsy. Bulk RNA-Sequencing (RNA-Seq) was performed on 10 samples in the total cohort. Both bulk and single-nucleus RNA-Seq (sNuc-Seq) were performed on the primary tumor from subject TRCC18 (**Fig. 1A, fig. S1, table S1**).

**Fig. 1:**
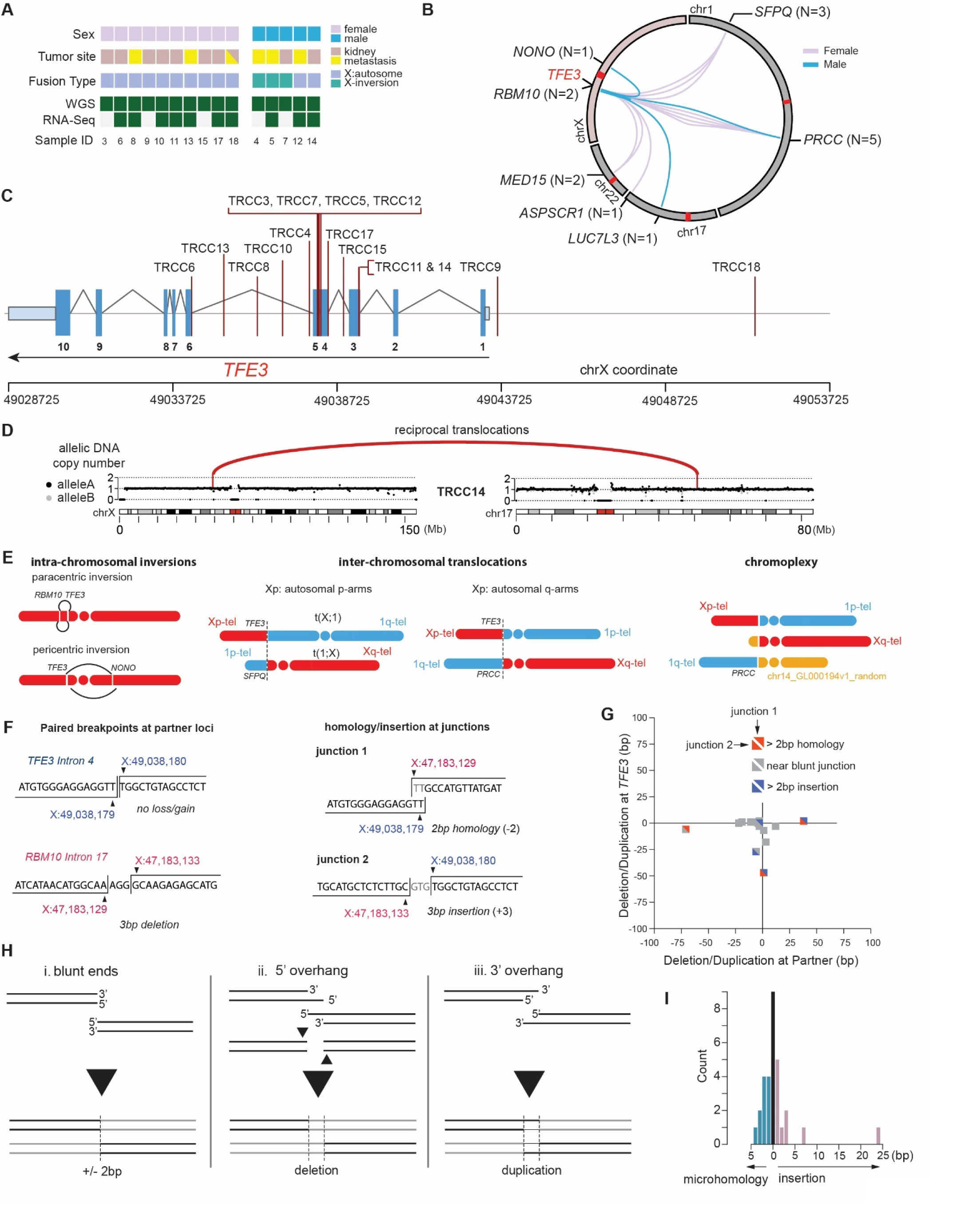
Etiology of TFE3 fusions revealed by tRCC whole-genome sequencing. **(A)** Overview of tRCC samples profiled by WGS and RNA-seq in this study. **(B)** Summary of *TFE3* fusions in the tRCC WGS cohort. Each line represents a *TFE3* fusion in an individual patient and is colored based on the sex of the patient. The total number of instances of each fusion is listed in parentheses. **(C)** Rearrangement breakpoints at the *TFE3* locus in the tRCC WGS cohort. **(D)** DNA copy number across chrX and chr17 in case TRCC14 with a *TFE3*-*LUC7L3* fusion. The fusion partners (*TFE3* on chrX and *LUC7L3* on chr17) are marked by vertical lines; the completely balanced DNA copy number indicates balanced reciprocal translocations, which are also supported by rearrangements (red arcs). **(E)** Summary of different types of rearrangements giving rise to *TFE3* fusions in the WGS cohort. From left to right: (1) intra-chrX inversion, including paracentric (i.e., inversion within the Xp arm) and pericentric (inversion spanning the centromere) inversion; (2) X:autosome translocations with autosomal partners on either the p-arm (e.g. *SFPQ*) or q-arm (e.g. *PRCC*) of an autosome; (3) chromoplexy, i.e., balanced translocations between three or more pairs of break ends (see **table S4**). **(F)** Sequence features of reciprocal translocations at *TFE3* and partner loci illustrated by an example of *TFE3-RBM10* fusion detected in TRCC7. *Left*: minimum DNA loss or gain at each partner locus. The example shows no DNA loss or gain at the *TFE3* breakpoint and a 3bp deletion at the *RBM10* breakpoint. *Right*: minimum homology or insertion at rearrangement junctions. The example shows a 2bp microhomology at the *RBM10*-*TFE3* junction and a 3 bp insertion at the *TFE3*-*RBM10* junction. **(G)** Summary of DNA loss/gain at fusion partner loci and junction homology of all cases in the tRCC WGS cohort. The x-and y-axes indicate the number of duplicated (+) or deleted (-) nucleotides at the *TFE3* translocation partner (x) and the *TFE3* breakpoint (y). The upper right and lower left triangles of each square show the classification of microhomology or insertion at reciprocal junctions (*Partner-TFE3* or *TFE3*-*Partner*). **(H)** Mechanisms that result in no change (**i**), deletion (**ii**), or duplication (**iii**) between opposite breakpoints generated by canonical non-homologous end-joining of two break ends generated by a single double-strand break. **(I)** Histogram showing the length of microhomology (left) or insertion (right) at *TFE3* fusion breakpoints. Both Partner-*TFE3* and *TFE3*-Partner breakpoints are pooled for this histogram. An apparent outlier with 24bp insertion at the *TFE3-MED15* junction detected in TRCC10 also shows a significant deletion (27bp) at the *TFE3* locus (see panel G), both of which are more consistent with microhomology-mediated end-joining.

We applied standard variant-detection methods across the cohort to call somatic mutations, copy number variants, and structural variants, with additional custom analyses to perform the detailed haplotype-specific reconstruction of genomic evolution in sample TRCC18 (see **Methods;** *Note*: only the primary tumor from TRCC18 and both samples from TRCC12 are included in cohort-level statistics that follow). The average frequency of single nucleotide substitution variants was 1.0 SNV/Mb (range 0.28-2.7 across frozen samples; FFPE samples were excluded from SNV analysis due to the high frequency of library artifacts in this data type(*30*, *31*)). The observed SNV/indel frequency in tRCC is somewhat lower than in clear cell renal cell carcinomas and most other solid tumors, and instead more similar to Ewing sarcoma, which carries a median substitution rate of <1 mutation/Mb (*16*, *32–34*) (**fig. S2; table S2**). One case (RCC16) was a clear outlier in terms of copy number profile and point mutation burden; this case carried chr3p loss, inactivating *VHL* mutation, and a *TFE3* fusion was not detectable, indicating that it was likely a misclassified ccRCC; it was excluded from further analysis in this study (**fig. S3**). We performed harmonized detection of copy number alterations and structural variants and observed a broad range of alteration frequency across the cohort, with some samples displaying pervasive segmental copy number alterations on a number of chromosomes (e.g. TRCC12, TRCC15, TRCC18) while other samples had virtually no copy number change (e.g. TRCC3, TRCC11, TRCC17; all three were treatment-naïve primary tumors) (**fig. S4**). While tumors from subject TRCC12 displayed many segmental copy number alterations, there were few differences in the copy number profile between pre-and post-treatment samples (**fig. S5**).

Given the small size of our WGS cohort relative to recent tRCC WES and panel-sequencing studies of tRCC (*7*, *13*, *14*), we did not perform statistical analysis of recurrently altered genes but assessed only the frequency of pathogenic variants in the Tier1 group of driver genes in the COSMIC Cancer Gene Census, or in genes previously reported to be recurrently altered in tRCC (*7*, *13*, *14*, *35*). Amongst these genes, only *CDKN2C* and *KMT2C* were mutated in two samples in our cohort with at least one of those mutations predicted to be of high pathogenic impact. While neither of these genes, to our knowledge, has been reported to be mutated in tRCC, *KMT2D* mutations have been noted in tRCC (*14*) and *CDKN2C* mutations are recurrent in glioblastoma (*36*). The latter finding is also intriguing given our prior report of *CDKN2A* alterations being the most recurrent alteration in tRCC apart from the *MiT/TFE* fusion (*7*). Indeed, in the current cohort, deletion spanning *CDKN2A* was observed in 6/15 patients, with clearly homozygous deletions in 4 of these cases (26%). One sample, TRCC15, showed biallelic inactivation of *ATM*, a mutation that has also previously been reported in a minority of tRCCs (*10*) (**fig. S2; table S3**).

### *TFE3* fusions universally arise by reciprocal translocations

We determined *TFE3* fusion breakpoints in WGS data and validated them in RNA-Seq data when available (**Methods; table S4**). There were seven distinct partners across 15 patients (2 on chrX and 5 on autosomes), all of which have been previously reported (*7*, *13*, *14*) (**Fig. 1B**). DNA breakpoints in *TFE3* were most commonly found between exons 4 and 6, which would be predicted to result in an in-frame fusion between *TFE3* and the partner gene. However, in two cases (TRCC9 and TRCC18), genomic breakpoints lay upstream of the gene body, which could lead to preservation of the full-length *TFE3* gene within the fusion product (**Fig. 1C**). Interestingly, in 3 cases (TRCC11, TRCC12, TRCC14), DNA breakpoints were consistent with the joining of an intron of the *TFE3* partner gene to an exon of *TFE3*, a rearrangement that would be predicted to result in an out of frame fusion product (**Fig. 1C and table S4**). In one of these cases (TRCC12), RNA-Seq data was available and demonstrated a fusion between exon 1 of *PRCC* and exons 5-10 of *TFE3*; DNA breakpoints were in intron 1 of *PRCC* and exon 4 of *TFE3*. This suggests exon skipping to restore the reading frame of the fusion, indicating the presence of a post-transcriptional splicing alteration acquired, and presumably required, by the tumor to maintain expression of the driver rearrangement (**fig. S6**).

*TFE3* fusions arose by four distinct mechanisms of rearrangement, all reciprocal: (1) paracentric inversion on Xp (2 cases); (2) pericentric inversion on chrX (1 case); (3) reciprocal X:autosome translocations (11 cases); and (4) chromoplexy (1 case) (**Fig. 1D-E**). Notably, the chromoplexy event in TRCC15 involved paired breakpoints on chr1, chrX, and a locus (chr14_GL00019v1_random) corresponding to the repetitive DNA sequence that may be located on the short arm of any acrocentric chromosome (chr13, chr14, chr15, chr21, or chr22). This case also contained multiple other reciprocal translocations that may be related to *ATM* inactivation observed in this sample (*37*). In contrast to oncogenic fusions in other cancers (*18*, *22–26*), we almost never detected complex rearrangements or copy-number alterations on either chrX or the partner chromosome. The only exception was TRCC18, where we detected complex somatic copy number alterations (SCNAs) on the partner chromosome (1p) in a subset of metastatic samples; the absence of these alterations in other metastatic lesions or the primary tumor suggests that the *TFE3* fusion was still originally generated by a simple reciprocal translocation in that case.

In summary, the DNA copy-number and rearrangement data suggest that *TFE3* fusions universally arise by reciprocal translocations.

### *TFE3* fusion junctions are consistent with canonical non-homologous end joining

We next sought to more carefully interrogate breakpoints in *TFE3* fusions at nucleotide resolution. The reciprocality of *TFE3* fusions enabled us to directly analyze DNA sequences at paired breakpoints and at rearrangement junctions, and to compare their features to rearrangements engineered in experimental systems (**Fig 1F**). Remarkably, we observed minimal DNA deletion or insertion in all cases at both the *TFE3* locus and the partner loci, including at all three loci in the case of chromoplexy (TRCC15). Among 31 pairs of break ends (3 in TRCC15 and 2 in each of the other 14 cases), 22 had deletions or insertions of no more than 10 base pairs; the largest deletion was 71bp (*ASPSCR1* in TRCC8) and the largest insertion was 38bp (*PRCC* in TRCC12). The absence of significant deletion or insertion between all paired break ends (**Fig. 1G**) is consistent with the break ends arising as blunt or near blunt double-strand breaks (DSBs) (**Fig. 1H**). We then assessed microhomology between rearrangement partners or insertions at the rearrangement junctions. 26/31 (84%) of junctions displayed ≤ 2bp of microhomologies or insertions, consistent with a mechanism of canonical non-homologous end joining (c-NHEJ) rather than alternative end joining (a-EJ) or single-strand annealing (SSA) (**Fig. 1I**). Therefore, *TFE3* rearrangements observed in human tumors recapitulate signatures of reciprocal translocations generated previously in experimental models and suggest that the same mechanism underlies fusions generated via chromoplexy (*38–40*). Importantly, the preservation of both translocations in tRCCs enables a direct assessment of DNA losses or insertions between a pair of break ends generated by a single double-strand break, which is generally not accessible in experimental models where only the oncogenic fusion is typically preserved.

To further demonstrate that *TFE3* fusions could arise from c-NHEJ of double-strand breaks, we used CRISPR/Cas9 to induce DSBs in intron 3 of *TFE3* and intron 1 of *PRCC* (a recurrent *TFE3* fusion partner located on chr1) in 293 human embryonic kidney cells. Following introduction of sgRNAs targeting both *TFE3* and *PRCC*, we were able to detect a *PRCC-TFE3* fusion in both genomic DNA and cDNA by polymerase chain reaction (PCR). Sequencing of the amplicons across the DNA breakpoints of both the *PRCC-TFE3* and *TFE3-PRCC* fusions revealed a tight distribution of small insertions and deletions (median indel size: *PRCC-TFE3*: 3 bp, *TFE3-PRCC*: 3.5 bp), remarkably consistent with what we observed in our tRCC WGS tumor data, and in line with an underlying c-NHEJ mechanism of *TFE3* fusion generation in human cells (**fig. S7**).

Based on our human tumor genomic data and experimental data, we conclude that *TFE3* fusions arise via c-NHEJ of paired DSBs in a majority of cases. Moreover, these fusions can be faithfully modeled in human cells using CRISPR/Cas9 engineering, consistent with a prior study that engineered translocations in human cells using designer nucleases (*44*).

### Preservation of reciprocal fusions in most cases of tRCC

We next interrogated large public datasets to extend our observation that *TFE3* fusions universally arise via reciprocal rearrangements. To assemble the largest cohort of tRCCs to date, we curated 203 *TFE3* tRCC cases from 11 different studies (including this study); 160 of these cases had whole-exome or whole-genome sequencing data; this included 35 cases that had originally been misclassified as other types of kidney cancer (**Fig. 2A)**. Across 203 cases with sex annotation, we observed a female sex bias in tRCC incidence (61.1% females [N=124]; 38.9% males [N=79]), in line with prior smaller studies (**Fig. 2B**). In addition to annotating fusion type and sex across these cases, we analyzed DNA copy number and allele-specific RNA expression using a harmonized workflow, where raw data were available (**Methods**).

**Fig. 2:**
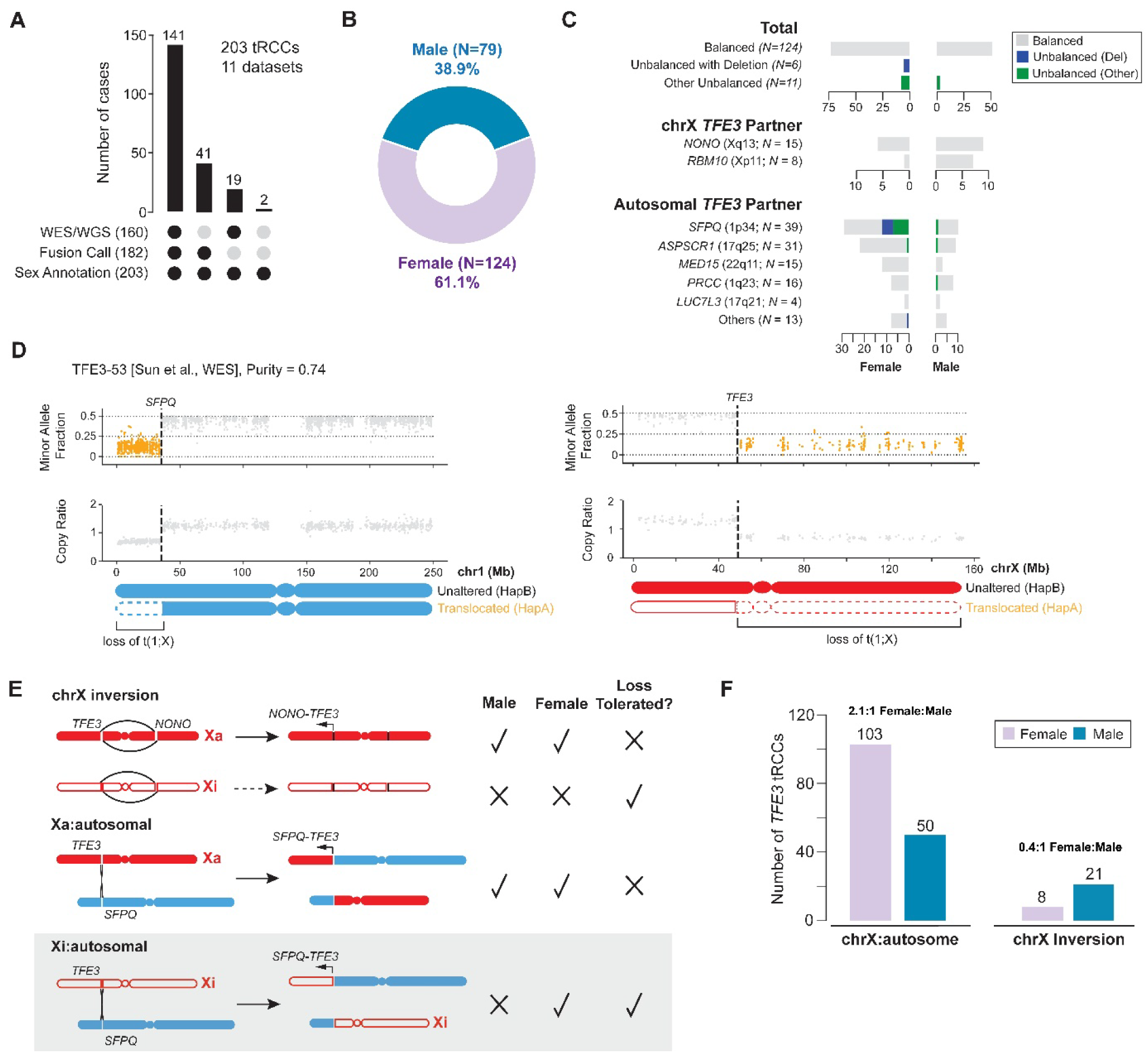
Sex differences in frequency of X:autosomal translocations underlies female predominance of tRCC. **(A)** Summary of the aggregated tRCC cohort generated from 11 independent studies (including the current study). **(B)** Sex distribution of tRCC cases in the aggregate tRCC cohort (*N* = 203). **(C)** Classification of *TFE3* fusions in the aggregate cohort based on the DNA copy-number status, sex, and rearrangement partner. Top (“Total”): Copy-number imbalance is rare in both female and male tRCCs and deletion is exclusive to female tRCCs. Middle (chrX intra-chromosomal inversions): No copy-number imbalance is detected in either female or male tRCCs. Bottom (X:autosomal translocations): There is female dominance in all autosomal *TFE3* partners except *PRCC* and *LUC7L3*, and deletion is exclusive to female tRCCs. Copy number imbalance consistent with loss of the *TFE3* reciprocal fusion is shown in blue while all other types of copy number imbalance involving either the *TFE3* fusion or the reciprocal *TFE3* fusion are in green. **(D)** An example of deletion of the reciprocal *TFE3-SFPQ* fusion t(1;X) in tRCC sample profiled via WES. Shown are the minor allele fraction (top) and copy ratio (tumor/normal depth, bottom) of both chr1 (left) and chrX (right). Data points in regions of allelic imbalance are highlighted in orange. **(E)** Constraints on each class of rearrangement leading to *TFE3* fusions and the copy-number status of translocated chromosomes imposed by the genetic differences in chrX between males and females. Intra-chromosomal inversions can only lead to oncogenic fusions on the active (filled red) but not the inactive chrX (open red) due to the silenced epigenetic state chrXi; by contrast, chrXi:autosomal translocations may generate oncogenic *TFE3* fusions if accompanied by partial activation of *TFE3* (bottom). Importantly, deletion of the reciprocal translocation would only be tolerated in female tRCCs when the translocation is generated on the inactive X. **(F)** The frequency of intra-chromosomal inversions and X:autosome translocations in either male or female samples (N=182) indicates that the female tRCC dominance is largely caused by X:autosomal translocations with a near exact 2:1 female-to-male ratio. The slight enrichment of intra-chromosomal inversions in male patients (*P*=1.2e-2, binomial test against 1:1 female-to-male ratio) is largely attributed to the higher prevalence of *RBM10-TFE3* fusions in male tRCCs; *RBM10* shows a male bias in mutation frequency(*51*).

Even though one cannot typically detect the *TFE3* rearrangement junctions themselves in WES data, the reciprocality of *TFE3* rearrangements can be inferred based on DNA copy number on chrX and on the *TFE3* fusion partner chromosome. The presence of segmental deletions or duplications on either chrX or the partner chromosome would be suggestive of complex rearrangements being involved in the generation of *TFE3* fusions. By contrast, fusions arising via simple reciprocal translocations or inversions on chrX would be expected to have no large segmental or arm-level SCNAs. In the case of chrX:autosome translocations, copy-number imbalance at either the *TFE3* locus or at the partner locus could indicate either: (1) a translocation that was unbalanced at the onset (e.g., resulting from large terminal deletion or retention generated by chromosome bridge resolution); or (2) a balanced translocation that was followed by subsequent copy number change, such as loss of the translocated chromosome not containing the oncogenic fusion (*45*, *46*) (**Fig. 2C**).

Strikingly, across our large aggregate cohort of tRCCs, we found that nearly all *TFE3* rearrangements or translocations are devoid of copy-number alterations or imbalance on chrX and the partner chromosome. Deletions spanning the breakpoint on the reciprocal *TFE3* fusion chromosome occurred in only 6/141 (4.3%) cases, and all of these were female samples (Unbalanced (Del) category in **Fig. 2C**). Other unbalanced events (Unbalanced (Other) category in **Fig. 2C**) included gains of the *TFE3* fusion chromosome or of the reciprocal *TFE3* fusion chromosome, as well as losses on autosomal arms from the breakpoint to the telomere (without any copy number change on chrX, suggesting their origin in a chromoplexy event, such as tRCC-19, TFE3-50, and TFE3-48 in **fig. S8**) and were observed in an additional 7.8% of cases (11/141; 8 females, 3 males).

Intriguingly, a majority of the samples with deletion of the reciprocal *TFE3* fusion chromosome (83.3%, 5/6) had *SFPQ-TFE3* fusions (**Fig. 2C-D; fig. S8**). We speculate that such events could result in the inactivation of tumor suppressor genes located on the deleted portion of chr1p. Notably, *ARID1A*, a SWI/SNF component and tumor suppressor gene whose haploinsufficiency has also been linked to tumorigenesis, is located in this region (*47*, *48*); SWI/SNF components are also known to be mutated in kidney cancers, including tRCC (*7*, *10*, *49*); this region was also deleted in case RCC16 (ccRCC) (**fig. S3**).

The preservation of both translocated chromosomes in a majority of cases raises the question of whether the reciprocal *TFE3* fusion is required for oncogenesis. However, the observation that a small fraction of tRCCs display deletion of the reciprocal chromosome disfavors this hypothesis, as does the finding of two cases (TRCC9 and TRCC18) in which the *TFE3* fusion breakpoint occurs upstream of the *TFE3* gene body (and thus would not be predicted to result in a reciprocal fusion protein product).

A second possibility is that the preservation of both translocated chromosomes is not due to positive selection of both fusions but due to negative selection against deletions (**Fig. 2E**). For males (who have only one chrX), deletion of either translocated chromosome would be intolerable, as it would result in the deletion of essential genes on chrX (*50*). In females (who have one active chrX and one inactive chrX), translocations to chrXa would have similar consequences as in males. However, translocations involving chrXi would tolerate loss of the reciprocal chromosome, since they remove only the q-arm of chrXi (which is transcriptionally inactive) and an autosomal segment (which is compensated for by its homologous chromosome). Therefore, detection of either clonal or subclonal deletion of the reciprocal translocation strongly indicates that chrXi was accessed for the *TFE3* translocations. Importantly, the ability to generate *TFE3* fusions from both chrXa and chrXi in females could explain the clinically observed 2:1 female sex bias in tRCC.

Consistent with this model, we observed a near-exact 2:1 ratio (103 females, 50 males) of chrX:autosome fusions (*P*=1.3e-5 by binomial test) in our aggregate cohort. By contrast, *TFE3* fusions arising via chrX inversions showed a 2.63:1 male bias (21 males, 8 females; *P*=0.012). When excluding fusions involving *RBM10*, which have previously been reported to show a male cancer bias in mutation frequency (*51*), the male bias in chrX inversions was insignificant (11 males, 7 females; *P*=0.24 **Fig. 2F**). The absence of sex bias in fusions arising via chrX inversion is likely because fusions generated by chrXi inversion are likely to remain transcriptionally silenced. By contrast, the observation of the 2:1 female-to-male ratio of chrX:autosomal translocations and the exclusive presence of deletions in only female tRCCs strongly support a model of chrXi being involved in oncogenic *TFE3* fusions, ascribing a genetic mechanism to the female predominance of this cancer.

### Paired genomic and transcriptomic analyses establish chrXi involvement in *TFE3* fusions

We next turned our attention to case TRCC18, in which the primary tumor was collected at diagnosis and multi-site sampling was performed at the time of rapid autopsy (**Fig. 3A**). In this case, we detected deletions on the 1p-terminus telomeric to the translocation partner (*SFPQ*) in six out of 13 metastatic lesions. We performed joint DNA copy-number, rearrangement, and single-nucleotide variant detection and constructed the phylogeny of all metastatic lesions based on single-nucleotide variants (**Fig. 3B, fig. S9-S10**); this reconstructed phylogeny is also consistent with copy-number alterations (**Methods**).

**Fig. 3:**
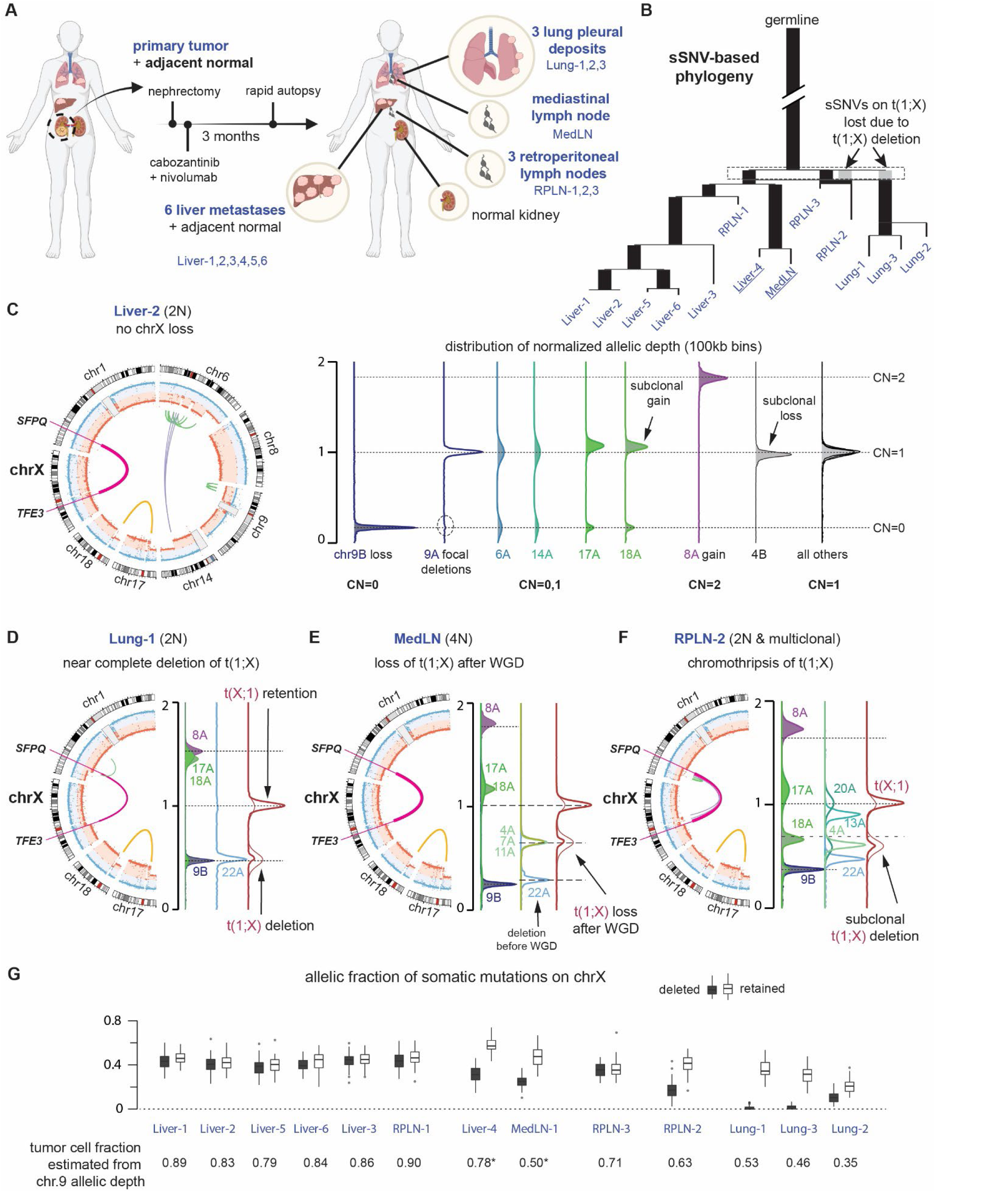
Genome evolution of tRCC reveals independent deletions of the reciprocal *TFE3* fusion. **(A)** Summary of systemic treatment history and anatomic sites of 14 tRCC samples from subject TRCC18. **(B)** Phylogenetic tree of the dominant tumor clones in 13 metastatic tRCC samples determined from single-nucleotide mutations (see Methods and **fig. S9-S10**). Ancestral branches (with more than one progeny clones) are shown as thick bars; terminal branches are shown as thin lines. Branch lengths are proportional to the number of somatic mutations. The gray bars preceding RPLN-2 and the Lung biopsies denote mutations on the translocated chromosome t(1;X) that were lost from these samples. Two samples inferred to have near tetraploid genomes (Liver-4 and MedLN) highlighted in underline (see panel E below). **(C)** Summary of ancestral copy-number alterations in all tumor biopsies using data from Liver-2. *Left*: CIRCOS plot showing haplotype-specific DNA copy number (blue and red), and rearrangements (green for intra-chromosomal rearrangements, purple for inter-chromosomal rearrangements) of chromosomes with copy number alterations. The reciprocal *TFE3-SFPQ* translocations are highlighted as a thick magenta line. *Right*: Distributions of haplotype-specific read depth data for chromosomes with altered DNA copy number (chrs.6,8,9,14,17,and 18). We reserve the A homolog for the haplotype with altered DNA copy number and the B homolog for the intact one; for chr9, the A homolog had focal deletions and the B homolog underwent complete loss. The integer copy-number states of each chromosome are determined by the allelic depths of the lost 9B homolog (CN=0), the gained 8A homolog (CN=2), and regions with normal DNA copy number (CN=1). There is a minor subclonal loss of the 4B homolog that is only present in the Liver-2 sample; subclonal gains of the translocated 17A and 18A are also observed in other samples and inferred to have arisen independently. All other alterations are shared by all metastatic lesions and the primary tumor. **(D-F)** CIRCOS and allelic-depth distribution plots of selected chromosomes showing downstream evolution after the ancestral changes as shown in **C**. (**D**) The lung metastases (using Lung-1 as the representative) show a near clonal deletion of most of the reciprocal translocation t(1;X) (**see fig. S10**), copy-number gains of 17A and 18A, and a near clonal deletion of 22A. (**E**) The MedLN and Liver-4 metastases (using MedLN as the representative) were inferred to be 4N based on the presence of 4A, 7A, and 11A at the median copy-number state between 22A (CN=0) and the basal copy-number state. Based on this inference, we further conclude that the loss of entire t(1;X) occurred after whole-genome duplication. (**F**) We infer the RPLN-2 to be polyclonal with many subclonal copy-number alterations (4A, 13A, 20A, 22A). Due to these alterations and the scarcity of point mutations, we cannot determine the number of subclones or their ploidy states. However, the deletion of t(1;X) shows the same clonality as the deletion of 4A and the p-arm of 20A, and these deletions cannot be accounted for by losses after whole-genome duplication as the copy-number states are lower than the median between complete deletion (CN=0 inferred from 9B) and the basal copy number state (CN=1 in 2N genomes or CN=2 in 4N genomes). **(G)** *Top*: Box plots of allelic fractions of somatic mutations on Xq in all metastatic biopsies. Each mutation is phased to either the retained (open boxes) or the deleted (filled boxes) homolog based on their allelic fractions in all samples. Each box plot shows the average variant allele fraction of mutations phased to each homolog in each sample. The Lung-1 and Lung-3 biopsies show complete deletion of mutations on the deleted homolog, whereas RPLN-2, Liver-4, MedLN show lower allelic fractions of these mutations consistent with incomplete losses. The Lung-2 sample is a polyclonal mixture. The mean allelic fraction of both groups of mutations in each sample is consistent with the clonal fraction of tumor cells estimated from the allelic depth of 9B (inferred to be deleted in the founding clone). The ratio between the average allelic fraction of mutations on the retained versus those on the deleted homolog in MedLN and Liver-4 is also consistent with a 2:1 copy-number ratio between the two homologs.

The *SFPQ*:*TFE3* fusion was first generated as a reciprocal translocation with completely balanced DNA copy number of both translocated chromosomes (chrX and chr1) (**Fig. 3C**) and remains preserved in five liver metastases (Liver-1,2,3,5,6) and in two metastases involving the retroperitoneal lymph nodes (RPLN-1 and RPLN-3). Copy-number imbalance between the oncogenic translocation t(X;1) and the reciprocal translocation t(1;X) is seen in the remaining samples. In the Liver-4/MedLN clade (**Fig. 3B, 3D**), we inferred that this imbalance is due to a single-copy loss after whole-genome duplication; this would have been tolerated regardless of whether the loss was of chrXa or chrXi, given the presence of two copies of each homolog after duplication. By contrast, we determined that there was complete deletion of t(1;X), containing the entire Xq arm in the clade of lung metastases, which are all of 2N ploidy (**Fig. 3B, 3E**). We further inferred the presence of a dominant 2N subclone in the RPLN-2 metastasis with complete deletion of t(1;X) (**Fig. 3F**). The deletion of t(1;X) in lung metastases and the loss of t(1;X) after WGD in Liver-4/MedLN are further validated by the complete absence of somatic mutations phased to the deleted X chromosome based on allelic imbalance in the lung metastasis, and by the approximately 2:1 allelic ratio of somatic mutations between the retained:deleted chrX homologs in the Liver-4/MedLN clade (**Fig. 3G**).

The identification of clonal deletion of t(1;X) in all three lung metastases and in a dominant clone in RPLN-2 strongly implies that the translocation was generated on the inactive X chromosome. To definitively demonstrate this, we assessed haplotype-specific expression of genes on Xq in the primary tumor, where both chrXa and chrXi were preserved; the estimated purity of this sample is ∼80%. We determined haplotype phase based on allelic imbalance on chrXq in lung metastases (**Fig. 4A-C**). Haplotype-specific expression to the right of the breakpoint in the primary tumor showed mono-allelic expression of the retained haplotype (chrXa) with the only exception being the *XIST* gene, which encodes the *XIST* RNA that orchestrates X-chromosome inactivation and is known to be mono-allelically expressed from chrXi. Together, these data unambiguously demonstrate that chrXi was involved in the oncogenic *TFE3* fusion in TRCC18 and that the non-functional translocation t(1;X) underwent frequent secondary loss or deletion after the initial translocations during tumor metastasis.

**Fig. 4:**
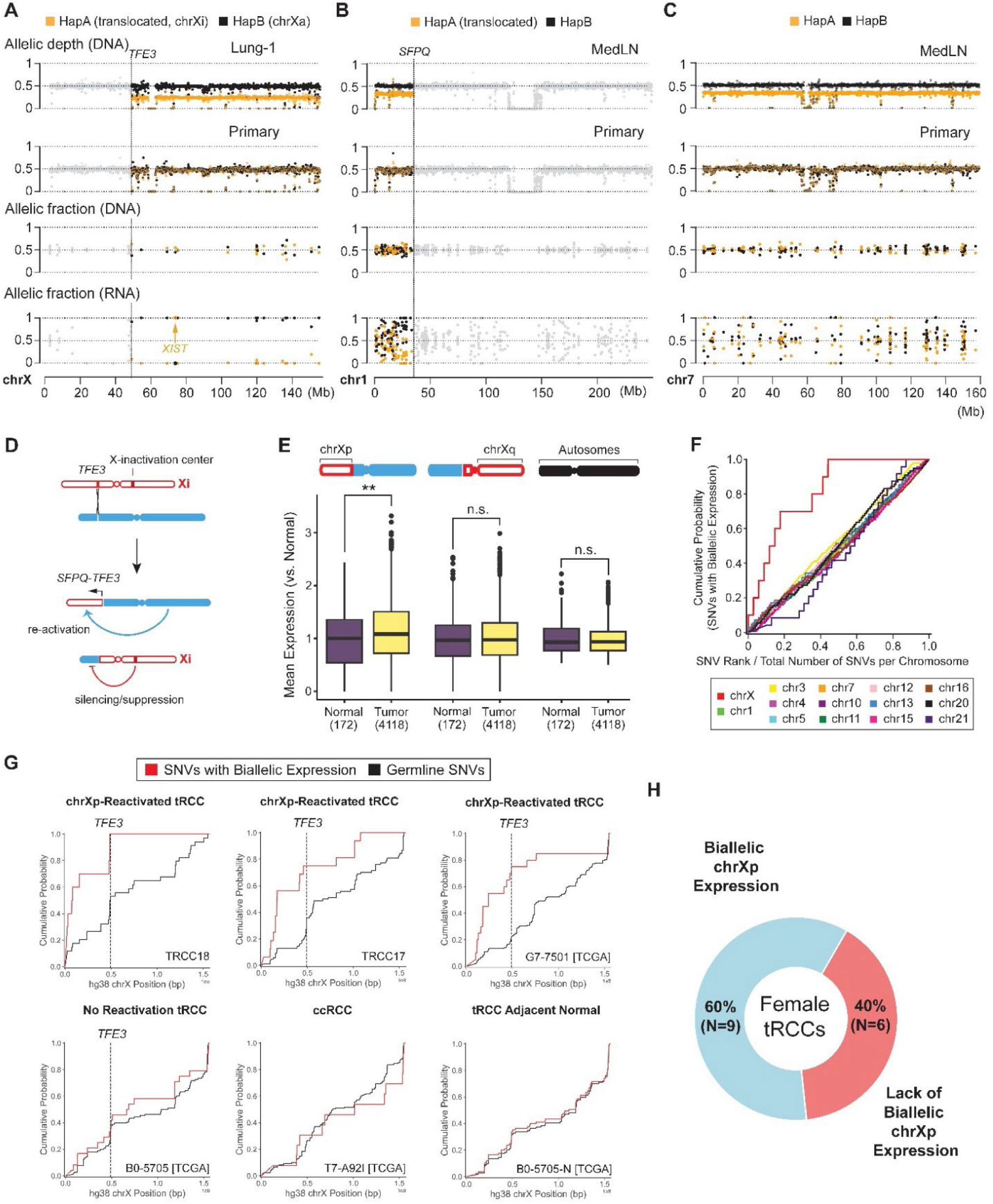
Xi:autosome translocations are associated with Xi reactivation. **(A-C)** Haplotype-specific transcription of chrX (A), chr1 (B) and chr7 (C) in the primary tumor from TRCC18. These chromosomes have segmental (chrX and chr1) or whole-chromosome allelic imbalance (chr7) in different metastases (first row), enabling the determination of haplotype phases of germline variants in colored regions (highlighted with orange and black dots corresponding to the allelic depths of parental homologs; allelic depths in regions in allelic balance are shown as gray dots). Shown in the 1^st^ and 2^nd^ rows are the allelic DNA depths at heterozygous variants in tumors with allelic imbalance (chrX: Lung-1, chr1 & chr7: MedLN) and in the primary tumor. All three chromosomes are disomic and in complete allelic balance in the primary tumor (second row). Haplotype-specific transcription (fourth row; shown is allelic transcription from expressed heterozygous variant sites) was assessed in the primary tumor, using phased variant genotype (third row; shown are allelic DNA fractions of reference and alternate genotypes at heterozygous sites). There is near complete monoallelic transcription on chrXq from the chrX homolog retained in lung metastases. The exception is a variant in *XIST* which is phased to the chrX homolog deleted in Lung-1, establishing that the inactive X (chrXi) was deleted in the lung metastases (and is also lost in Liver-4/MedLN and RPLN-2). Haplotype-specific transcription on chr1p also revealed allelic imbalance telomeric to the *SFPQ* locus, reflecting silencing of this region under the influence of XIC from the translocated chrXi. Chr7, disomic in the primary, has completely balanced allelic transcription. *P*-values calculated by Mann-Whitney U test between distributions of RNA minor allele fraction for heterozygous sites to the left and right of the *TFE3* breakpoint (chrX, *P* = 1.05e-3) or *SFPQ* breakpoint (chr1, *P* = 1.14e-2). **(D)** Predicted transcriptional consequences of Xi:autosome translocations. **(E)** Single nucleus RNA sequencing of TRCC18 primary tumor sample reveals evidence for increased transcriptional output from the translocated Xp segment. Transcriptional output is assessed for regions telomeric to *TFE3* breakpoint (left), chrXq (middle), or diploid autosomes (right) as indicated in the chromosome schematic. The distribution of normalized transcriptional output in each of the above regions is then plotted for malignant (N=4118) and normal cells (N=172) (see **Methods**). *P*-values calculated by Mann-Whitney U test (chrXp genes telomeric to TFE3: *P* = 6.3e-3 [**], chrXq genes: *P* = 0.48 [n.s.], diploid autosomes: *P* = 0.79 [n.s.]). **(F)** Cumulative fractions of biallelically expressed heterozygous variant sites (≥ 10 DNA reads and ≥ 10 RNA reads, minor allele fraction ≥ 0.2, excluding genes escaping XCI) on each chromosome (y-axis) compared to the cumulative fractions of heterozygous sites on each chromosome (x-axis). Only diploid autosomes and chrX are included. The fractions of heterozygous expression (y-axis) or heterozygosity (x-axis) are cumulated beginning at the p-terminus of each chromosome. Deviations from the diagonal (most obvious for chrX) indicate a positional divergence from biallelic expression. **(G)** Cumulative distributions of biallelic transcription (red) compared to cumulative distributions of DNA heterozygosity (black) on chrX (excluding genes escaping XCI) for selected female samples from the aggregate cohort in the following categories: female tRCC samples with autosome-*TFE3* fusions and chrXp reactivation (*P* < 0.05 by Mann-Whitney U test between distributions of minor allele fraction for heterozygous sites to the left and right of the *TFE3* breakpoint); female tRCC samples with no evidence of reactivation (*P* > 0.05); ccRCC; tRCC cancer-adjacent normal sample. See also **fig. S11**. **(H)** Donut plot of female tRCC samples with or without biallelic chrXp expression across the combined TCGA and DF/HCC cohorts. *P*-values calculated for each sample by Mann-Whitney U test between distributions of minor allele fraction for heterozygous sites to the left and right of the *TFE3* breakpoint.

### Xi:autosomal translocations lead to chrXi reactivation

X-chromosome silencing is orchestrated by the non-coding RNA *XIST* during early embryogenesis and stably maintained thereafter (*27*). It has been reported that *XIST* can induce epigenetic silencing of entire chromosomes when inserted ectopically and that spreading of chrX silencing may extend to some autosome:chrXi translocations (*52–54*). In the latter case, the translocation would bring autosomal regions under the influence of the X-chromosome inactivation center (XIC) located on chrXq. However, it is unclear whether translocations that sever chrXp from the XIC and join it with euchromatic autosomal arms may alter the silencing of chrXp, given the stable and heritable nature of X chromosome inactivation. For *TFE3* fusions involving chrXi to be oncogenic, they would have to be accompanied by at least partial reactivation of chrXi, minimally at the *TFE3* locus, but possibly extending further from the rearrangement breakpoint to the telomere (**Fig. 4D**).

In the TRCC18 tumors, chrXp is in allelic balance, which prevents determination of the haplotype phase. We therefore sought to assess the degree of chrXi reactivation by analyzing the frequency of bi-allelic expression, which would indicate transcription from chrXi. Strikingly, we observed a significant increase of biallelic gene expression on the chrXp arm telomeric to the *TFE3* breakpoint in comparison to the chrXq arm (**Fig. 4A**, *P*=1.05e-3 by Mann-Whitney U test between distribution of minor allele fraction for heterozygous sites to the left versus right of the *TFE3* breakpoint; see also **fig. S11**). The region of chromosome 1 telomeric to the *SFPQ* breakpoint also displayed features of partial transcriptional silencing, consistent with spreading from the adjacent chrXi (*P*=1.14e-2 by Mann-Whitney U test between distribution of minor allele fraction for heterozygous sites to the left versus right of the *SFPQ* breakpoint, **Fig. 4B**). By contrast, bi-allelic expression on an autosome with allelic balance in the primary tumor (chr7) is evenly distributed across the entire chromosome (**Fig. 4C).** We further examined this pattern across all chromosomes and found that only chrX displayed a positional bias in biallelic SNV expression, with the transition point located precisely at the *TFE3* breakpoint (**Fig. 4F-G**).

We then performed single nucleus RNA-Seq (sNuc-Seq) of the TRCC18 primary tumor (4,290 cells) and assessed the total level of gene expression on Xp in tumor cells in comparison to normal cells. With reactivation of genes on the p-arm of chrXi, we expected to see a higher level of total chrXp expression (left of the *TFE3* breakpoint) in tumor cells as compared with normal cells, in which chrXi is silenced. Consistent with this prediction, we observed increased transcriptional output from chrXp in tumor cells relative to non-malignant cells. By contrast, there was no difference in transcriptional output between tumor and normal cells from chrXq or from disomic autosomes (**Fig. 4E)**. These data further support partial reactivation of genes located on the telomeric side of the *TFE3* breakpoint in t(X;1) selectively in tumor cells.

Finally, we interrogated whether X chromosome reactivation induced by chrXi:autosome *TFE3* translocations is also detectable in other tRCCs in our aggregate cohort described above. We restricted our analysis to RNA-Seq data generated from frozen samples (6 female tRCCs from our cohort including TRCC18, 9 from external cohorts). FFPE samples were excluded due to the lower quality of the RNA-Seq data limiting our ability to infer allelic expression (*55*, *56*). In 9/15 (60.0%) female tRCCs with autosome-*TFE3* fusions, we observed evidence of chrXp reactivation as indicated by an enrichment of bi-allelic expression telomeric to the *TFE3* breakpoint (*P*<0.05 by Mann-Whitney U test between distribution of minor allele fraction for heterozygous sites to the left versus right of the *TFE3* breakpoint; **Methods**). We did not observe a significant difference in the minor allele fraction distributions of X-linked genes on either side of the *TFE3* breakpoint in adjacent normal samples, which display consistent biallelic expression across chrX due to mosaicism of XCI; nor did we observe a significant difference in a cohort of randomly selected cohort of female ccRCC samples from the TCGA, which were copy-neutral on chrX (9 samples, size-matched with TCGA female tRCC cohort; **Fig. 4G-H and fig. S11**). Therefore, reactivation of Xp genes may be a general outcome of oncogenic *TFE3* fusions generated by chrXi:autosome translocations.

As further supporting evidence for chrXi *TFE3* rearrangements, we sought to determine whether any existing patient-derived cell line models display evidence for chrXi rearrangements. We performed Western blotting for TFE3, using an antibody against the C-terminus of the protein in a panel of *TFE3* fusion cell lines. We expected cell lines with chrXi fusions to display two bands (wild-type TFE3 from chrXa, TFE3 fusion from reactivated chrXi), while those with chrXa fusions would only display one band (TFE3 fusion from chrXa, with wild-type TFE3 subject to XCI on chrXi). Indeed, two *TFE3* fusion cell lines (sTFE and ASPS-KY) displayed a double band while only one band was seen in the remaining lines. Use of a shRNA targeting the *TFE3* 3’UTR in sTFE and ASPS-KY cells reduced the intensity of both bands, confirming that they both represent TFE3 species (**fig. S12**). We conclude that the inactive X chromosome can be recurrently accessed for somatic rearrangements in tRCC, resulting in varying degrees of reversal of XCI.

## Discussion

We performed whole-genome analysis on a cohort of 29 tRCC tumor samples and reanalyzed genomic data and annotations from 203 tRCC specimens from 11 different studies (*7*, *13*, *14*, *57*). This represents, to our knowledge, the most comprehensive whole genome analysis to date of this rare cancer. Our investigation provides insights into several outstanding questions about the etiology of *TFE3* fusions, and more broadly, lends insight to the implications of somatic chrX rearrangements.

We found that *TFE3* fusions almost uniformly arise via reciprocal rearrangements or translocations, including chromoplexy, with minimal loss or gain of genetic material at break points on each chromosome involved in the rearrangements (**Fig. 1D**). This is in contrast to oncogenic fusions in other tumors where a varying fraction of the fusions arise from complex rearrangements (*16–18*). This observation has two implications for the etiology of *TFE3* fusions. First, the finding of minimal microhomologies or insertion at fusion breakpoints (**Fig. 1G**) recapitulates the observations of engineered oncogenic fusions in human cells (*44*) and suggests an origin in c-NHEJ after near-blunt double-strand breaks (*44*, *58*). The precise mechanisms that generate near blunt DSBs *in vivo* remain to be elucidated. Moreover, the presence of minimal DNA loss or gain at *TFE3* and partner loci contrasts with other cancers in which fusions arise via reciprocal translocations or chromoplexy with 10-100kb deletions between paired break ends; this raises a possibility that such fusions may arise from different mechanisms of DNA breakage (*16–18*). Second, preservation of the reciprocal *TFE3* fusion in a majority of cases reflects unique constraints imposed by the monoallelic transcription of chrXa in both males and females: without whole-genome duplication or reactivation of chrXi, no large segmental deletion of chrXa is tolerable due to the presence of essential genes on chrX (*50*). By contrast, segmental or whole-chromosome loss of chrXi is likely tolerable and may occur frequently in female cancers. Given the drastically different consequences of chrXa or chrXi copy-number alterations, haplotype-specific copy-number analysis is essential to uncovering the functional implications of alterations involving chrX.

The observation of rare copy-number imbalance across the *TFE3* breakpoint in female tRCCs both indicates that the reciprocal *TFE3* fusion is dispensable for oncogenesis and suggests that the inactive chrX can be involved in the translocation, which allows cells to tolerate chrXq deletion. To the extent that the *TFE3* reciprocal fusion is preserved in a majority of tRCCs, our data indicate that this is dictated by tumor evolutionary constraints from chrX, but not due to functional selection for retaining the reciprocal fusion.

We provide definitive evidence that chrXi not only can produce oncogenic *TFE3* fusions but that the translocation is also accompanied by at least partial reactivation of the p-arm of chrXi that forms the translocated chromosome with an autosome (and partial silencing of autosomal regions brought *in cis* with the XIC by the translocation), supporting recent findings reporting somatic perturbation of XCI in cancer cells (*59*, *60*). The fact that chrXi can be accessed for functional *TFE3* fusions through chrXi:autosome translocations presents a genetic explanation for the observed female sex bias in tRCC, with a striking 2:1 female-to-male ratio of chrX:autosome translocations. Many cancers show differences in incidence or outcomes between males and females (*61*, *62*) that cannot be fully explained by differences in lifestyle, environment, or hormonal factors alone. Our study highlights how baseline genetics of the sex chromosomes can be linked to sex differences in cancer incidence and outcome. This study also raises intriguing mechanistic questions about whether specific factors promote somatic rearrangements to arise on chrXi (versus chrXa or autosomes), given that chrXi is distinct in epigenetic state, chromatin ultrastructure, physical location and DNA replication timing (*63–67*).

Finally, several prior studies in experimental model systems have shown that disruption of the X-inactivation center (XIC) usually has modest effects on maintaining XCI, though the phenotype varies based on context (*68–71*). tRCC represents a physiologic context in which a large segment of chrX is physically disconnected from the XIC and therefore represents a model in which to study the somatic plasticity of XCI in cancer. Our data suggest that *TFE3* rearrangements can result in at least partial reactivation of silenced chrX genes and partial silencing of autosomal genes in some cases. More broadly, somatic rearrangements involving chrXi may underlie sex-specific transcriptional differences in other cancers.

## Acknowledgements

We are deeply grateful to the patients and their families for their generosity in contributing to this research project. We acknowledge the Broad Institute Genomics Platform and Center for Cancer Genomics (DFCI) for assistance with sample preparation and sequencing.

## Author contributions

M.A., A.S., C.B. made equal contributions to different aspects of this project and are ordered based on the duration of their commitment to this project. M.A. led the whole-genome sequencing analysis; C.B. performed haplotype-specific DNA copy-number calculation and the phylogenetic analysis of multi-site biopsies; A.S. identified and curated tRCC cases from external cohorts, performed the DNA copy-number and allelic expression analysis on these cases, performed X-reactivation analyses, and performed experimental work related to engineering of the *TFE3* fusion via CRISPR/Cas9. F.M. analyzed *TFE3* fusion breakpoints. F.M., Q.X., and M.N. assisted in copy number analysis of the whole-genome sequencing cohort. P.K. performed the single-cell RNA-Seq analysis. R.W.T. performed initial DNA copy-number analysis on WGS samples. J.L. assisted in engineering of the *TFE3* fusion via CRISPR/Cas9. D.S.G. generated immunoblots. J.O’T, D.F, D.S.G., U.A.A., G.M.L., J.L.H, E.C.K., D.J.E., T.K.C. contributed clinical samples and/or data. S.R.V. conceived the project. S.R.V and C.-Z.Z designed and supervised the analysis. S.R.V., C.-Z.Z, M.A., A.S., C.B. wrote the manuscript with help from all authors.

## Data and materials availability

Sequencing data reported in this study have been deposited in the database of Genotypes and Phenotypes (dbGaP) under study accession phs003008.v1.p1 with controlled access in accordance with an IRB-reviewed protocol. Algorithms and code implemented in this study are described in **Methods**; scripts are uploaded at https://github.com/SViswanathanLab/WGS-TRCC. Any additional information required is available upon reasonable request.

