## Supplemental Materials (Methods, Supplementary Figures S1-S12 and Captions) for "A genetic basis for cancer sex differences revealed in Xp11 translocation renal cell carcinoma"

### Materials and Methods

#### Selection and curation of tRCC and matching germline DNA samples

A total of 31 tRCC tumors from 16 individuals (including two tumors from one misclassified ccRCC female case), each with matched germline control (either peripheral blood or adjacent normal tissue) were collected for whole-genome sequencing and RNA sequencing (excluding formalin-fixed paraffin embedded samples). One subject (TRCC18) had nephrectomy specimen plus tissue specimens from 13 metastatic sites collected at rapid autopsy. A second subject (TRCC12) had both pre- and post-progression biopsies taken from the same metastatic site. Age range of the cohort was 25-68 years old; N=10 females; N=6 males,

All individuals who donated samples provided written informed consent consistent with genomic analysis of freshly collected or banked tumor samples with blood/normal tissue control. Rapid autopsy samples for subject TRCC18 were collected under an IRB-approved rapid autopsy protocol that allows for genomic profiling.

#### Sequence data generation

##### Whole genome sequencing (WGS)

Tumor and matching germline tissue samples were subjected to whole genome sequencing in one of three categories: (1) standard PCR-free WGS performed on frozen tissue (26 samples); (2) standard WGS with PCR performed on FFPE tissue (4 samples); (3) linked-read WGS performed on frozen tissue (4 samples). DNA and RNA extraction, library preparation, and whole genome whole transcriptome sequencing was performed by the Genomics Platform at the Broad Institute using the standard procedures as described below. For single nucleus RNA sequencing, extraction, library preparation, and sequencing was performed at the Center for Cancer Genomics (DFCI).

*Standard PCR-free WGS*

For each tumor or germline reference sample, an aliquot of genomic DNA (100ng in 50µL) was used for library construction. Genomic DNA was fragmented to a target size of 385bp by shearing using a Covaris focused-ultrasonicator, with additional size selection performed using a SPRI cleanup. Library preparation was performed using a commercially available kit provided by KAPA Biosystems (KAPA Hyper Prep with Library Amplification Primer Mix, product KK8504), and with palindromic forked adapters using unique 8-base index sequences embedded within the adapter (purchased from Roche). DNA libraries were sequenced on the HiseqX platform (2 x 151bp mode) at the Broad Institute of MIT and Harvard.

*Standard WGS with PCR*

For FFPE samples, the DNA libraries were further amplified by 10 cycles of PCR. Following sample preparation, libraries were quantified using quantitative PCR (kit purchased from KAPA Biosystems) with probes specific to the ends of the adapters; the qPCR quantification was performed on Agilent's Bravo liquid handling platform. Based on qPCR quantification, libraries were normalized to 2.2nM and pooled into 24-plexes. Multiplexed DNA libraries were sequenced on the NovaSeq 6000 platform (2 x 151bp) at the Broad Institute of MIT and Harvard.

For standard WGS with or without PCR, unmapped reads were aligned and subjected to post-alignment processing following the best practice workflow recommended by the GATK without base score recalibration.

*10x Genomics WGS*

Linked-read DNA libraries were constructed using the standard protocol from 10X Genomics, starting with 1.2 ng of DNA for each sample as previously described (*72*). Library fragment sizes were determined using the DNA 1000 Kit and 2100 BioAnalyzer (Agilent Technologies) and quantified using qPCR (KAPA Library Quantification Kit, Kapa Biosystems). Libraries were sequenced the HiSeqX platform (2 x 151bp mode). Sequencing reads (FASTQ files) were processed by the LongRanger software (10X Genomics) for barcode-aware alignment and variant calling.

**RNA Sequencing**

*Bulk RNA-seq*

Bulk RNA sequencing libraries were constructed for 10 frozen tumor samples. DNA/RNA extraction was performed using AllPrep (Qiagen). Total RNA was quantified using the Quant-iT™ RiboGreen® RNA Assay Kit and normalized to 5ng/ul. Following plating, 2ul of ERCC controls (using a 1:1000 dilution) were spiked into each sample. An aliquot of 200ng for each sample was used for library preparation with the TruSeq™ Stranded mRNA Sample Preparation Kit. The resultant 400bp cDNA then went through dual-indexed library preparation ('A' base addition, adapter ligation using P7 adapters, and PCR enrichment using P5 adapters). Libraries were quantified using Quant-iT PicoGreen (1:200 dilution) and normalized to 5ng/ul. Pooled libraries were quantified using the KAPA Library Quantification Kit for Illumina platforms.

Pooled libraries were normalized to 2nM and denatured using 0.1 M NaOH prior to sequencing. Sequencing was performed on the Illumina NovaSeq 6000 platform (2 x 101bp, single 8bp index) at the Broad Institute of MIT and Harvard. Sequencing data were processed as described below.

*Single-nucleus RNA-seq*

Single-nucleus RNA-Seq was performed on the nephrectomy sample from TRCC18. Nuclei isolation was performed as previously described (*73*). Briefly, OCT-embedded tissue fragments were quickly thawed in PBS on wet ice to remove excess OCT, tissue was next mechanically dissociated by pipetting and homogenized in TST solution, filtered through a 30 µm MACS SmartStrainer (Miltenyi Biotec, Germany), and pelleted by centrifugation for four minutes at 500g at 4C. The nuclei pellet was resuspended in 200 µL of ST-SB buffer, and nuclei were counted by eye using INCYTO C-Chip Neubauer Improved Disposable Hemacytometers (VWR International Ltd., Radnor, PA, USA). Low-retention microcentrifuge tubes (Fisher Scientific, Hampton, NH, USA) were used throughout the procedure to minimize nuclei loss.

Approximately 8,000 nuclei were loaded into a single channel of the Chromium Next GEM Chip G for library construction on the 10x Chromium Controller (10x Genomics, Pleasanton, CA, USA) following manufacturer's instructions (Chromium Next GEM Single Cell 3' Reagent Kits v3.1 User Guide, Rev D). Libraries were normalized and pooled for sequencing on a NovaSeq SP-100 flow cell at the Center for Cancer Genomics at Dana-Farber Cancer Institute.

**DNA Sequence Data Processing and Variant Calling**

All sequencing data were aligned to GRCh38. Linked-reads data were aligned by the LongRanger program from 10X Genomics. Standard WGS data were aligned using bwa.

**Germline and somatic variant calling by HaplotypeCaller**

Variant calling was performed jointly on tumor and matched normal samples collected from each patient using HaplotypeCaller from Genome Analysis Toolkit (GATK) as described previously (*46*). The same parameters were used for variant calling for all libraries; differences in downstream processing for each data type is described below.

To select germline heterozygous variants for allelic DNA and RNA analysis, we imposed three criteria: (1) variant is included in the 1000 genomes reference data;
ftp.ebi.ac.uk/1000genomes/ftp.1000genomes.ebi.ac.uk/vol1/ftp/data_collections/1000G_2504_high_coverage/working/20201028_3202_phased/
(2) heterozygous genotype emitted by HaplotypeCaller; (3) five or more reads of both the reference and alternate genotypes in the germline reference sample. We recognized the presence

of contaminating DNA (from unrelated individuals) in the germline reference of TRCC15 and tumor sample of TRCC17 based on the presence of many common variants with low allelic fractions. To eliminate these variants from DNA copy-number analysis, we additionally imposed the following criteria: (4) the allelic fractions of both reference and alternate genotypes need to be within the range of 0.35 to 0.65; (5) the combined allelic depth needs to be equal to or greater than 20. These additional criteria were unnecessary for the other samples.

For linked-read data, we directly performed haplotype phasing using mLinker, which can purify false heterozygous variants based on haplotype linkage(*74*).

For standard WGS data, we further selected somatic variants in the callset with the following criteria: (1) variant sites not among common polymorphisms in the 1000 Genomes Phase 3 reference haplotype panel; (2) homozygous genotype in the germline reference; (3) five or more reads of the alternate genotype across all tumor samples. The resulting somatic variant callsets were used to intersect with the somatic callsets generated by Mutect2 to improve variant calling accuracy.

**Somatic mutation detection with Mutect2**
Somatic mutation detection was performed on all WGS samples using MuTect2 v4.3.0 from Genome Analysis Toolkit (GATK). All samples were run in paired tumor-normal mode.

To filter false variants due to recurrent alignment errors, we used a 'reference' panel of variants provided by GATK:

gs://gatk-test-data/mutect2/M2PoN_4.0_WGS_for_public.vcf

To filter false variants due to recurrent alignment errors, we imposed the following read filters:

--minimum-mapping-quality 30 (excluded reads having low mapping quality)
--read-filter OverclippedReadFilter
--filter-too-short 25
--downsampling-stride 20 (truncate regions of extreme depth due to mapping error)
--max-reads-per-alignment-start 6
--max-suspicious-reads-per-alignment-start 6

To filter rare artifacts and germline variants that were missed in the matching germline reference, we used a germline resource consisting of >10,000 genomes from gnomAD provided by GATK:

gs://gatk-best-practices/somatic-hg38/af-only-gnomad.hg38.vcf.gz

For three patients with multiple tumors (TRCC12, RCC16, and TRCC18) or multiple normal references (TRCC18), Mutect2 was run in the joint-calling mode with similar filtering parameters as described above.

**Filtering of somatic mutations**
For FFPE samples (TRCC3, TRCC4, TRCC7 and TRCC9), we did not analyze sSNVs (including INDELs) given the depth of sequencing and the high rate of sequencing errors introduced during library construction resulting from DNA degradation (*30*, *31*).

For linked-read data (TRCC5, TRCC6, TRCC8 and TRCC9), we intersected variant callsets generated by the LongRanger pipeline with callsets generated by Mutect2 using `bcftools` (`bcftools isec`). This intersection was performed to exclude low confidence variants generated by LongRanger and false positive variants generated by Mutect2 (which does not incorporate molecular barcode information present in linked-read data).

For callsets generated from standard WGS data, we performed the following filtering steps. (1) FilterMutectCalls to filter false variants due to contaminating DNA and artifacts that may introduce orientation bias (such as 8-oxoguanine artifacts) by specifying tumor contamination, segmentation and read orientation models. (2) FilterAlignmentArtifacts to remove alignment artifacts from poorly mappable regions of the genome. All variants located in centromeres were also removed.

We imposed a hard filter of variant AD ≥ 3 for all detected mutations. For TRCC18, we intersected variants generated by Mutect2 with those generated by HaplotypeCaller and selected bi-allelic single nucleotide variants for phylogenic inference. We further removed variants with missing genotype calls ('./.') in one or more samples in the Mutect2 callset and variants with either missing genotype calls or called as homozygous reference by HaplotypeCaller.

**Variant annotation with VEP**
All somatic variants that passed filters were annotated using Ensembl Variant Effector Predictor (version 106) with default parameter settings. Given the cohort size, we limited to somatic variants within Tier 1 genes from Cancer Gene Census (CGC; GRCh38 COSMIC v96). For downstream analyses, these Tier 1 filtered genes were assessed for impact and pathogenicity prediction from VEP106. Mutations were visualized with python package CoMut (*75*). For the CoMut plot, genes are shown if they were mutated with HIGH/MODERATE impact in at least two samples in the cohort.

**Somatic DNA Copy-Number and Rearrangement Analysis**
DNA copy number was calculated using a haplotype-specific copy-number workflow (*46*). For standard WGS, haplotype phasing was performed by `eagle2` using common variants and haplotype phase from the 1000 genomes consortium as reference (this included both single-nucleotide variants and insertion/deletion variants); for linked-reads data, haplotype phasing was performed using mLinker (*74*) on germline variants detected in the germline reference linked-reads data (this only included SNVs). Local haplotype fractions were calculated based on allelic depths at heterozygous variants and their haplotype phase in either 10kb intervals (samples with high purity and more SCNAs: TRCC12, TRCC13, TRCC14, TRCC15, TRCC16, TRCC18) or 30kb intervals (samples with few SCNAs or low purity: TRCC3, TRCC4, TRCC7, TRCC9, TRCC11, TRCC17; linked-reads samples: TRCC5, TRCC6, TRCC8, and TRCC10). The larger interval size was chosen to either maximize the detection sensitivity of low-clonality SCNAs in FFPE samples (TRCC3/4/7/9), in samples with no large-scale SCNAs (TRCC3/11/17), and in linked reads samples with more variable total sequence coverage at 10kb intervals (TRCC5/6/8/10). For linked-read data, allelic depth was calculated from the number of unique molecular barcodes at each variant site using mLinker; for standard WGS data, allelic depth was generated by HaplotypeCaller.

The combined sequence coverage (from both haplotypes) was calculated from the normalized sequencing coverage in 10kb intervals (or aggregated into 30kb intervals); for linked reads data, the combined sequence coverage was calculated from the number of unique molecular barcodes using mLinker.

The allelic depth was calculated from the combined sequence coverage and the haplotype fractions. We then performed haplotype phasing based on allelic imbalance as described in (*46*). We calculated and reviewed 100kb-interval DNA copy number generated from the allelic-depth phased chromosomal haplotype, and manually corrected rare long-range switching errors (usually 0-3 per sample). The final haplotype and haplotype-specific DNA copy number (in 100kb intervals) were used for downstream DNA copy-number analysis.

**Rearrangement detection**
We identified rearrangements based on discordant (including split mapped) reads as described previously (*46*). Candidate rearrangements were emitted when there were at least two distinct discordant reads within the maximum insert size of the DNA library (~1000bp) at each candidate

breakpoint region. All *TFE3* fusion breakpoints were manually reviewed (breakpoint coordinates listed in **table S4**).

For three samples with FFPE DNA (TRCC3/4/9), we only considered *TFE3* fusion breakpoints as they were assessed to have either low tumor cell fractions (TRCC3/4) based on the number of reads supporting the TFE3 fusions or displayed few large segmental SCNAs (TRCC9). We manually reviewed copy-number changepoints and identified 11 rearrangements in TRCC7.

For linked-reads data, we imposed the following filtering criteria on candidate rearrangements: (1) The breakpoints need to be at least 100kb away from each other or reside on different chromosomes (due to the prevalence of short-range chimeras in linked-reads data); (2) The number of unique molecular barcodes of all supporting reads needs to pass a sample-specific threshold (10 for TRCC6 and TRCC10, 6 for TRCC8); (3) The number of unique molecular barcodes of supporting reads in the normal sample needs to be lower than 5% of the number of unique molecular barcodes of supporting reads in the tumor sample. Only rearrangements with one or both breakpoints co-localized with copy-number changepoints are included in the CIRCOS plots. There were no long-range breakpoints detected in TRCC5 when reviewed against DNA copy-number data.

For other standard WGS data, we required at least 5 supporting discordant reads (10 reads for TRCC12 or RCC16 with two tumor biopsies) for each candidate rearrangement. We further imposed the following criteria: (1) Supporting reads only come from each individual sample (this is not imposed on TFE3 fusions as the breakpoints may be in close proximity); (2) There are at least two non-split discordant reads and at least one split read; (3) The total number of supporting reads from all normal reference samples is less than three. Last, we intersected the rearrangement breakpoints with copy-number changepoints and excluded short insertions, deletions, or tandem duplications (<100kb). We manually reviewed all breakpoints against the DNA copy-number plots.

#### Inference of Purity/Ploidy

We performed tumor purity/ploidy inference using TITAN (*76*) but verified the purity/ploidy solution based on the haplotype-specific allelic depths generated for each chromosome (46 for female, 45 for male). For samples with few or no SCNAs, we further used the AF distribution of somatic mutations to corroborate the purity estimates (for non-FFPE samples). The final results are presented in **table S1**.

#### Phylogenetic inference in TRCC18

Due to the low overall burden of both sSNVs (including INDELs) and SCNAs, we performed phylogenetic inference on all 13 tumor samples (excluding the primary) from the TRCC18 patient by a joint analysis of sSNVs and SCNAs.

*Somatic SNV-based Phylogenetic Reconstruction in TRCC18*

To infer the phylogenetic relationship based on sSNVs in TRCC18, we first determined the appropriate variant allelic fraction (VAF) cutoff for clonal sSNVs in each tumor sample. We performed *k*-means ($k$ = 2) clustering on the VAF data from each tumor sample to determine the cutoff separating high and low VAF variants. We verified these sample-specific cutoffs against the tumor cell fractions estimated from the allelic depth of the deleted chr.9 haplotype that was inferred to be a truncal event shared by all tumors (and must be clonal in each sample). We further determined a set of truncal sSNVs that passed the VAF threshold in 9 or more tumor samples based on the histogram of sample presence of all sSNVs; based on these truncal variants, we independently verified the sample-specific VAF cutoffs (nearly all truncal variants passed the cutoff in each sample) and estimated the rate of false negative detection based on the number of truncal variants that do not meet the VAF cutoff.

We next calculated the number of sSNVs shared by two or more tumor samples and constructed the phylogenetic tree by neighbor-joining. We started by grouping sample pairs with the most shared sSNVs (e.g., MedLN and Liver-4), merged the pairs into one node in the phylogenetic tree, and iteratively constructed a binary tree from all samples.

There are several noteworthy caveats. First, there are 28 mutations shared by all samples except the lung tumors (Lung-1/2/3) and RPLN-2, 9 mutations present in all but Liver-4 plus the same lung and RPLN samples, and 14 mutations shared by all samples except Lung-1/3 and RPLN-2. Nearly all these variants (51 total) reside on the translocated chromosome t(1;X) that was deleted in the lung samples, in the dominant clone of RPLN_2, and underwent single-copy loss in MedLN and Liver-4 after whole-genome duplication. Therefore, the absence of these mutations in the Lung/RPLN/Liver-4 is most plausibly due to deletion of this chromosome, instead of reflecting the accumulation of new mutations in the remaining samples.

Second, the Lung-2 sample shared 39 mutations with the other two Lung biopsies, but also contained 14 mutations that were absent in Lung-1/3 but present in the Liver and MedLN samples, mostly on t(1;X). A close examination of the somatic copy-number data revealed that the Lung-2 sample was both polyclonal and had low tumor cell fractions. We therefore concluded that the conflicting sSNV evidence may have been due to the mixing of tumor DNA (possibly during rapid autopsy) and grouped Lung-2 with the other two lung biopsies because of their genetic similarity as well as anatomical proximity.

Finally, the RPLN-2 sample has very few shared mutations with other samples (excluding the truncal mutations). This is not due to low tumor cell fraction in the RPLN-2 sample (the tumor cell fraction was estimated to be ~60%) as we detected a large number of private mutations in this sample (second only to Liver-3). Moreover, we detected a private chromothripsis event on the t(1;X) that is specific to this sample. We grouped RPLN-2 with RPLN-3 based on 8 mutations shared between them but note that RPLN-2 likely diverged early from the other tumors and also acquired more genetic alterations during downstream evolution.

*Further Validation of somatic SNV-based phylogeny*

After determining the phylogeny, we then assigned each mutation to a branch in the phylogenetic tree based on its allelic fractions in al tumor samples. For each mutation classified as not clonal or not present in a given sample (based on the VAF cutoff), we assigned a false-negative detection likelihood based on the fraction of truncal mutations that were missed in this sample; for each mutation classified as clonal in a given sample, we assigned a constant false-positive rate (0.01). We then calculated the combined likelihood of each branch assignment based on the presence or absence of this mutation in all the samples and assigned this mutation to the branch with the maximum likelihood. We grouped mutations by clonality patterns and reviewed the genome-wide distribution of mutations to identify mutations whose absence arose from chromosomal deletions or losses. We manually adjusted their branch assignment as needed.

*Consistency between SSNV based phylogeny and SCNA-based phylogeny*

We inferred most segmental SCNA breakpoints to be ancestral, which were not useful for phylogenetic inference. Both MedLN and Liver-4 samples were inferred to be tetraploid based on the allelic depth distribution; this inference is consistent with their adjacency in the phylogenetic tree. In addition, Liver-4 and MedLN also share breakpoints on 17p and 22q that are distinct from the other samples. We observed independent SCNAs in different phylogenetic branches, including (1) deletion of chr22A; (2) subclonal gains of segments from 17 and 18 that were joined together by an ancestral rearrangement; (3) loss of chr4A; (4) deletion of the translocation t(1;X). We verified that the deletion of 22A was either of the entire chromosome (which may occur independently in different branches), or showed distinct deletion boundaries (e.g., between Lung and Liver-4). The same pattern was verified for deletions of t(1;X). In summary, there are no recurrent segmental SCNA breakpoints in any two tumor samples that are not immediately adjacent in the phylogenetic tree.

### RNA-Sequencing Analysis

#### Bulk RNA-Seq Analysis

Paired-end FASTQ files were aligned to the hg38 genome build using STAR (v2.7.10a), followed by fusion analysis using STAR-Fusion (v1.7.0) with the GRCh38 gencode v37 CTAT library (vMar012021). The identified gene fusions were subsequently assessed using Integrative Genomics Viewer (IGV).

#### snRNA-Seq analysis

Demultiplexing of sequencing reads, barcode processing, read alignment, and UMI counting were performed using the 10x Cell Ranger analysis pipeline. The reads were aligned to the human genome reference, GRCh38 from GENCODE and extracted RNA counts were analyzed using the Seurat V4 algorithm (*77*). Doublet detection and removal on gene-barcode matrices were performed to exclude data from droplets containing more than one cell using Scrublet (*78*). Cells with fewer than 200 genes detected or more than 5% of counts attributed to mitochondrially-encoded transcripts were removed. Genes detected in fewer than three cells were also excluded. Tumor cells were identified based on known tRCC gene markers (TRIM63 and GPNMB) and inferred transcriptional copy number variations (CNVs) estimated via the inferCNV package that were consistent with WGS-based copy number profile (*79*). Normal cell types were determined through manual annotation via known marker genes. For arm-level expression analysis, the mean expression level across all genes belonging to the specific chromosome region was calculated for each malignant and normal (non-tumor compartment) cell. For autosome analysis, disomic autosomes were considered (chromosomes 1, 3, 4, 5, 7, 10, 11, 12, 13, 15, 16, 20, and 21). The mean expression of respective chromosome arm per cell was then normalized to the average expression of that arm across all non-malignant cells. Statistical significance was identified using a Mann-Whitney U test between tumor vs normal of individual chromosome arms. For autosome analysis, disomic autosomes were considered (chromosomes 1, 3, 4, 5, 7, 10, 11, 12, 13, 15, 16, 20, and 21).

#### Allele-Specific Expression

RNA data were uniformly re-aligned using TopHat2 (*80*) and DNA was re-aligned using bwa mem (*81*) to GRCh38 fasta reference, using bedtools bamtofastq (*82*) to convert deposited bam files to fastq (paired-end for the cohort from this study using inner distance between mate pairs of 100) and without preserving mate information for TCGA due to differing sequencing protocols for different samples. We called germline variants using GATK HaplotypeCaller followed by GenotypeGVCFs (*83*), filtering the vcf to heterozygous SNVs. We then used ASEReadCounter to count reference and alternative RNA reads across the heterozygous sites (*83*). We restricted to sites with total RNA and DNA count of at least 10. We further restricted to sites with germline DNA variant allele fraction within one standard deviation of the mean for the sample (to reduce the impact of mapping errors and germline copy number variation). For analysis of chrX, we filtered pseudoautosomal regions (PAR1 hg38 coordinates: 1-2781479, PAR2 hg38 coordinates: 155701383-156030895), escapee exons, and variable escapee exons as defined by kidney-specific methylation (*84*) and coordinates determined from Ensembl BioMart (*85*). We determined whether chrXp reactivation was present by comparing the distribution of minor allele fraction for expressed SNVs left versus right of *TFE3* using a two-tailed Mann-Whitney U test. For TRCC13, due to very low sequencing depth (total reads: 4,495,792 vs. median reads for other female samples in our cohort: 134,598,212), only two non-escapee coding SNVs had sufficient RNA read count on chrX meeting the above criteria; we therefore excluded this sample from allele-specific expression analysis.

**Haplotype-Specific Expression in TRCC18**

Common germline variants left of the *TFE3* breakpoint were statistically phased and switching errors were corrected using haplotype-specific expression pipeline previously described (*46*). Germline heterozygous variants right of the breakpoint were phased using allelic imbalance. We took samples with clonal deletions right of the breakpoint (no WGD): 'Lung-1', 'Lung-3' ('Lung-2' was not used due to being polyclonal and having low tumor cell fraction, as discussed above) and compared them to normals: 'BTRCC18Normal', 'KidneyNormal', 'LiverNormal'. For these two sets of samples, we summed DNA reference allele counts and summed DNA alternative allele counts at each SNV. The cumulative reference allele fraction (summed reference allele counts / (summed reference allele counts + summed alternative allele counts)) was calculated at each SNV for the two sets of samples. If it was greater in the cohort with deletions (Lung-1 and Lung-3) right of the *TFE3*, the variants were inferred to be on the retained allele (HapB); otherwise the variants were inferred to be on allele deleted in lung metastases (HapA). So as not to determine allelic fraction in outlier copy number regions, we restricted our analysis to 100kb intervals where HapA and HapB copy number on chrX was within one standard deviation of the mean for the tumor sample. For chr1 and chr7, 'MedLN' and 'Liver-4' were used for phasing, as these samples have copy number change on those chromosomes. Lung metastases were not used for chr1 phasing as these sites had a distinct structural variant resulting in partial retention of part of the chr1p reciprocal fusion chromosome. The allele-specific expression pipeline was then applied to obtain haplotype expression fractions, using the aforementioned assigned haplotypes.

### Analysis of External Cohorts

#### Public Datasets

Data from external cohorts were obtained and, if necessary, tRCCs detected as previously described for TCGA, Durinck et al., Malouf et al., MSK Panel, MSK WES, Oncopanel, and Sato et al. (*7*).Raw sequencing data and metadata for Sun et al. (*13*) and Qu et al. (*14*) were obtained from the respective studies. For *TFE3* tRCC samples in Sun et al. with RNA-seq data but lacking fusion calls, we ran STAR fusion (v1.7.0) on the RNA-seq to infer *TFE3* fusion partners. As Sun et al. lacked patient sex annotations, we assessed chrX heterozygosity in the matched normal samples to determine sex, as previously described (*59*).

#### Copy Number Analysis

tRCC samples from published cohorts were uniformly processed with the TitanCNA pipeline (*76*) using the baitsets specified in the source publications and converted to hg38 (UCSC LiftOver) if needed, with the following parameters TitanCNA_maxNumClonalClusters: 3; TitanCNA_maxPloidy: 4; ichorCNA_normal: c(0.25, 0.5, 0.75); ichorCNA_ploidy:c(2,3,4); ichorCNA_includeHOMD:TRUE; ichorCNA_minMapScore: 0.75; ichorCNA_maxFracGenomeSubclone: 0.5; ichorCNA_maxFracCNASubclone:0.7. The optimal cluster solution calculated by TitanCNA was taken as the most likely solution in the absence of manual review. Copy number imbalance (CNI) was determined by manual review analysis of allelic copy number on chrX and the autosomal fusion partner in all cases. Deletion or gain from the breakpoint to the telomere on at least one arm was considered CNI. All samples with CNI are shown in **fig. S8** (purity/ploidy-corrected allelic TITAN copy number, overlayed on the allelic .seg file) and annotated in **table S7**.

### Experimental Work

#### Cell Lines

293T and 786-O were obtained from American Type Culture Collection (ATCC, CRL-3216 & CRL-1932). S-TFE was obtained from the RIKEN BioResource Research Center (BRC) Cell Bank (RCB4699). UOK146 was obtained from Dr. W. Marston Linehan's laboratory (National Cancer Institute). ASPS-1 was obtained from Dr. Robert H Shoemaker's laboratory (National Cancer Institute). ASPS-KY was obtained from Dr. Shunsuke Yanoma's laboratory (Niigata University).

All cells were cultured in DMEM supplemented with 10% FBS, 100 IU/mL penicillin, 100 µg/mL streptomycin, and 100 mg $mL^{-1}$ Normocin at 37°C in 5% $CO_2$.

Generation of *TFE3* Fusion via CRISPR/Cas9

sgRNAs targeting intron 1 of *PRCC* and intron 3 of *TFE3* were cloned into lentiCRISPR v2 (Plasmid #52961 on Addgene, **table S6**) (*89*). The combination of sgRNAs were transfected into 293T cells using 4:1 polyethylenimine (PEI):DNA. After 72 hours, total RNA was isolated with the RNeasy Plus Mini Kit (Qiagen, #74136). cDNA was synthesized from 1 µg of total RNA using SuperScript IV VILO Master Mix (Thermo Fisher Scientific, #11756050). Genomic DNA was isolated using the DNeasy Blood & Tissue kit (Qiagen, #69504). cDNA or gDNA was amplified using appropriate primers (see **table S6**) for 40 cycles and run on a 1% agarose gel containing SYBR Safe DNA gel stain (1:10000). Gels were imaged using UV light.

Amplicon Sequencing and Analysis

Amplicons were excised from gels and DNA was isolated using QIAquick Gel Extraction Kit (Qiagen, #28706X4). Amplicons were submitted for linear/amplicon long-read sequencing (PlasmidSaurus). Reads were first aligned to a fasta reference containing *TFE3* and *PRCC* for inference of the breakpoint location, using bwa mem. Using this information, a .fasta reference consisting of the *PRCC-TFE3* translocation (hg38 breakpoint: chr1:156777745-chrX:49038604) and *TFE3-PRCC* translocation (hg38 breakpoint: chrX:49038605-chr1:156777746) was created (spanning a 2000 bp window around the *PRCC-TFE3* and *TFE3-PRCC* breakpoints). Reads were re-aligned to this reference using bwa mem. The aligned breakpoint-spanning reads were visualized in IGV to determine the number of nucleotides of insertion or deletion across the consensus breakpoint.

Immunoblotting

The cells were lysed on ice using RIPA lysis buffer (Thermo Fisher Scientific, #89901) supplemented with protease inhibitors (Roche, #11836170001). The amount of protein in lysates was determined using the Pierce BCA Protein Assay Kit (Thermo Fisher Scientific, #23225). Equal quantities of protein were then loaded onto NuPAGE 4–12% Bis-Tris Protein Gels (Thermo Fisher Scientific, #NP0335) and separated by SDS-PAGE. The proteins were transferred from the gels to nitrocellulose membranes using an iBlot2 (Thermo Fisher Scientific) and incubated overnight at 4°C with rabbit anti-TFE3 (CST, #14779S, 1:1000). Afterward, the membranes were washed in TBS-T and then incubated at room temperature for 1 hour with IRDye 680LT goat anti-Rabbit IgG: LI-COR (#926–68021, 1:20,000). Immunoblots were imaged using the Odyssey CLx Infrared Imaging System (LI-COR Biosciences).

### Supplementary Figures and Captions:

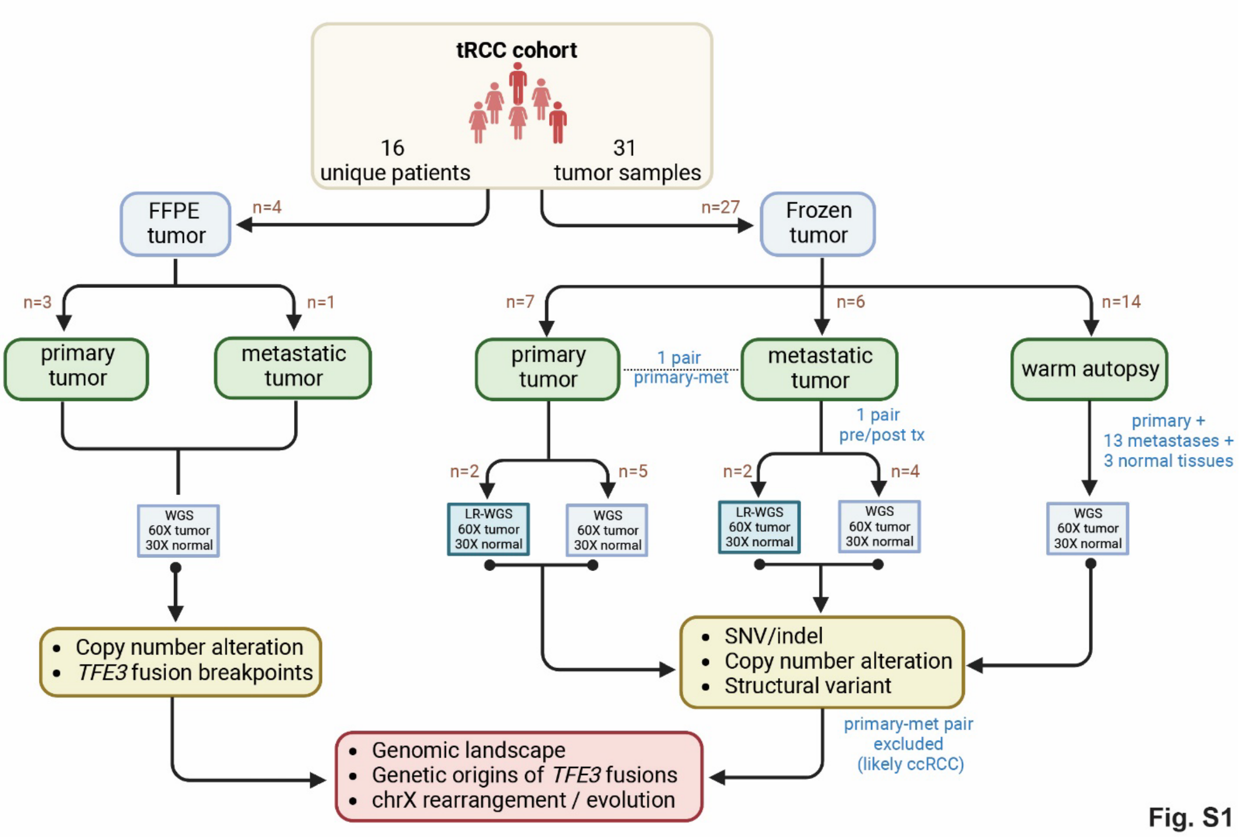


**Fig. S1. Samples profiled by DNA and/or RNA Sequencing in this study.** Flowchart summarizing the workflow of WGS analysis in this study. Two samples (TRCC1 and TRCC2) that failed DNA extraction are not included; two tumors later reclassified as ccRCC (RCC16) are included. For subject TRCC18 (total of 17 samples), primary tumor and adjacent normal were collected at the time of initial nephrectomy and the remaining 15 samples were collected at warm autopsy (see **Fig. 3**).

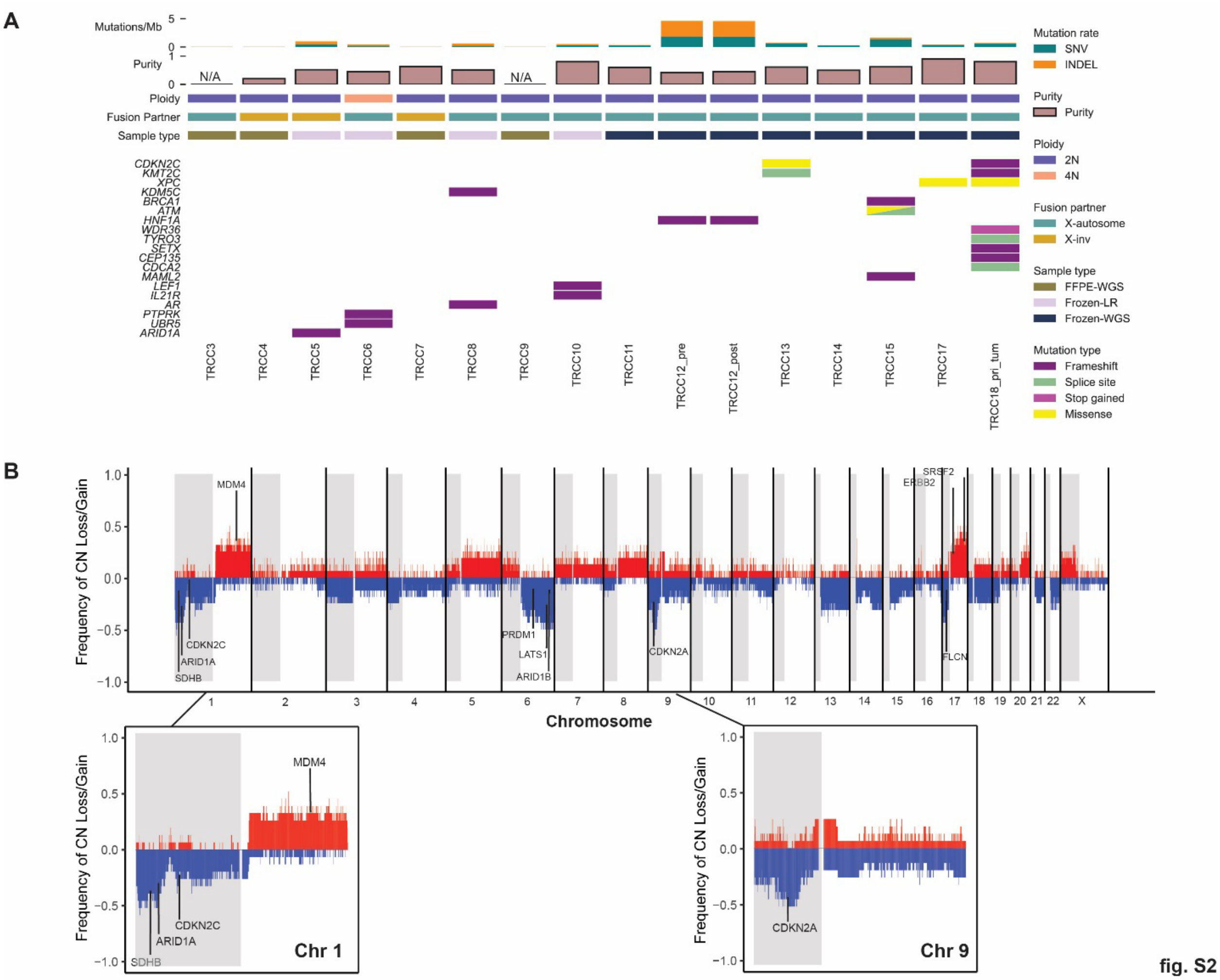


**Fig. S2. Recurrent mutations and copy number alterations in tRCC WGS cohort. (A)** Co-mutation plots of recurrent single-nucleotide substitution and insertion/deletion mutations of Tier 1 genes in the cancer gene census (CGC) in the tRCC WGS cohort. Only cancer genes with mutations predicted to have high functional impact in at least one sample, or high/moderate impacts in two or more tRCCs samples are included. Note: Mutation analysis was not performed for FFPE samples (TRCC3/4/7/9) due to technical challenges in accurately calling mutations in this data type (*30*, *31*) and due to the relatively low clonal fractions of tumor cells in some FFPE samples (e.g. TRCC3/4). Purity could not be reliably assessed in two of these samples (TRCC3/TRCC9) due to absence of SCNAs. **(B)** Genome-wide frequency of copy number changes per 100 kB genomic bin across the tRCC WGS cohort. Gains are depicted above the horizontal and losses below the horizontal. Known cancer drivers in peak/valley regions are annotated; two insets highlight recurrent focal deletions on chr1p (spanning *CDKN2C*, *ARID1A*) and 9p (spanning *CDKN2A/2B*).

**A**

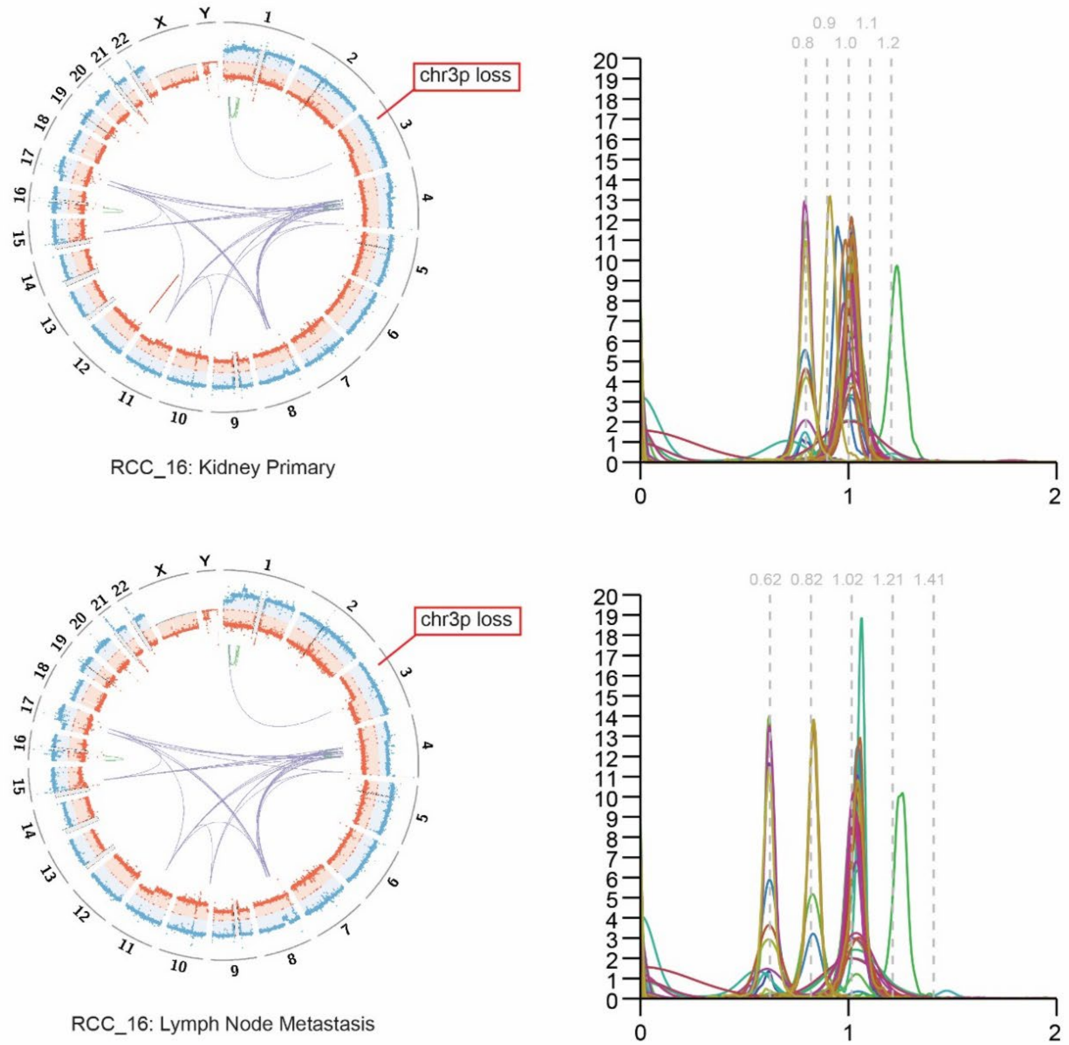
chr3p loss
RCC_16: Kidney Primary
0.8 0.9 1.0 1.1 1.2
chr3p loss
RCC_16: Lymph Node Metastasis
0.62 0.82 1.02 1.21 1.41

**B**

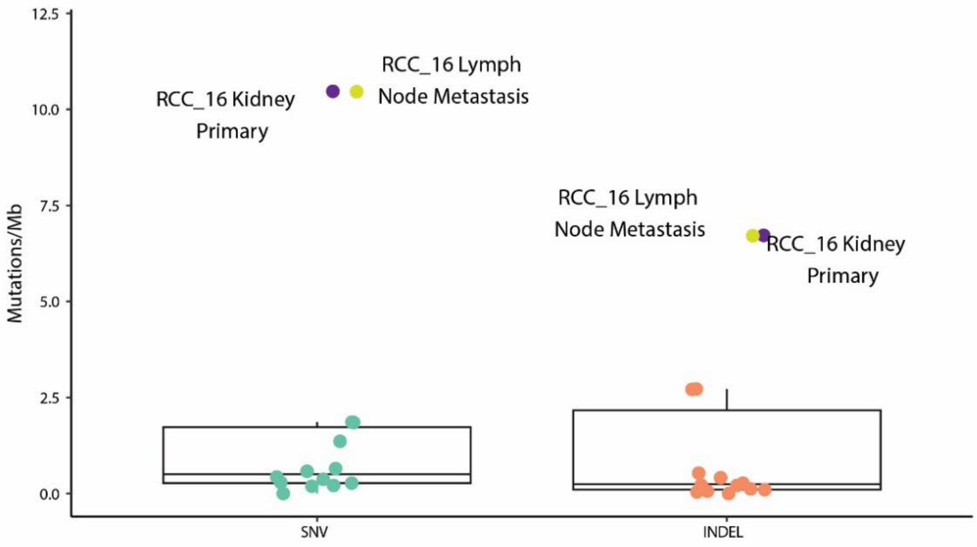
Mutations/Mb
RCC_16 Kidney Primary
RCC_16 Lymph Node Metastasis
SNV
INDEL

fig. S3-1

C

| Chr location | Gene | Consequence | Impact | CDS Pos | Codons |
|---|---|---|---|---|---|
| chr1:36466673-36466674 | CSF3R | Frameshift | HIGH | 2194-2195 | gac/gGac |
| chr1:110341568-110341569 | RBM15 | Frameshift | HIGH | 2163-2164 | aaCTct/aact |
| chr2:15940588 | MYCN | Start lost | HIGH | 2 | aTg/aCg |
| chr3:10149852 | **VHL** | Stop gained | HIGH | 529 | Aga/Tga |
| chr3:47123930 | **SETD2** | Stop gained | HIGH | 322 | Aga/Tga |
| chr3:52402771-52402806 | BAP1 | Splice donor | HIGH | 1902 | - |
| chr3:52563324 | **PBRM1** | Frameshift | HIGH | 4045 | Ctg/tg |
| chr3:70976939 | FOXP1 | Splice donor | HIGH | - | - |
| chr3:186052063-186052064 | ETV5 | Frameshift | HIGH | 1277-1278 | cTC/c |
| chr7:75541926 | HIP1 | Frameshift | HIGH | 2945 | gAt/gt |
| chr8:17966176 | PCM1 | Frameshift | HIGH | 3033 | gaT/ga |
| chr9:15490116 | PSIP1 | Frameshift | HIGH | 158 | tTa/ta |
| chr11:108320709 | ATM | Splice acceptor | HIGH | - | - |
| chr13:32380135-32380136 | BRCA2 | Frameshift | HIGH | 9246-9247 | -/A |
| chr14:55616528 | KTN1 | Frameshift | HIGH | 535 | Aaa/aa |
| chr15:44711588-44711591 | B2M | Frameshift | HIGH | 42-45 | tcTCTT/tc |
| chr20:32369110 | ASXL1 | Frameshift | HIGH | 224 | cTt/ct |
| chr1:114399498 | TRIM33 | Missense | MODERATE | 3079 | Gat/Aat |
| chr1:179110302 | ABL2 | Missense | MODERATE | 1760 | tCt/tTt |
| chr2:42328909 | EML4 | Missense | MODERATE | 2365 | Ggg/Tgg |
| chr2:120091594-120091596 | EPB41L5 | Inframe deletion | MODERATE | 1083-1085 | gcAAGa/gca |
| chr2:197396105 | SF3B1 | Missense | MODERATE | 3490 | Gac/Tac |
| chr3:169084947 | MECOM | Missense | MODERATE | 3313 | Gcg/Acg |
| chr5:177092510 | FGFR4 | Missense | MODERATE | 917 | aAg/aGg |
| chr6:44260240 | NFKBIE | Missense | MODERATE | 1240 | Acc/Gcc |
| chr6:167922934 | AFDN | Missense | MODERATE | 2966 | cAc/cGc |
| chr7:2919390 | CARD11 | Missense | MODERATE | 2492 | aCc/aTc |
| chr7:138579364 | TRIM24 | Missense | MODERATE | 2417 | gAt/gGt |
| chr8:70174797 | NCOA2 | Missense | MODERATE | 322 | Ggt/Agt |
| chr9:95150077 | FANCC | Missense | MODERATE | 532 | Gag/Aag |
| chr10:8055704 | GATA3 | Missense | MODERATE | 49 | Gcc/Acc |
| chr12:56095784 | ERBB3 | Missense | MODERATE | 2033 | aGg/aTg |
| chr12:65066733 | WIF1 | Missense | MODERATE | 638 | cTt/cCt |
| chr12:132668387 | **POLE** | Missense | MODERATE | 2142 | gaA/gaC |
| chr15:90893375 | FES | Missense | MODERATE | 2006 | gCt/gTt |
| chr16:64991851 | CDH11 | Missense | MODERATE | 728 | gGa/gAa |
| chr17:7673803 | **TP53** | Missense | MODERATE | 817 | Cgt/Tgt |
| chr17:31182589 | NF1 | Missense | MODERATE | 812 | aTc/aCc |
| chr17:43091696 | BRCA1 | Missense | MODERATE | 3835 | Gca/Aca |
| chr19:17841512 | JAK3 | Missense | MODERATE | 1019 | tCg/tTg |
| chr19:18747056 | CRTC1 | Missense | MODERATE | 385 | Gac/Aac |
| chr19:52220245 | PPP2R1A | Missense | MODERATE | 1359 | gaT/gaA |

**fig. S3-2**

**Fig. S3. Genomic features of RCC16 are more consistent with clear cell RCC than tRCC.** **(A)** Genome-wide DNA copy number and somatic rearrangements of the primary tumor (top) and a resected metastatic lesion (bottom) from subject RCC16. A *TFE3* fusion could not be identified in either sample form this patient, suggesting that it may have been misclassified. The following additional copy-number/rearrangement features of this genome are consistent with features of a clear-cell renal cell carcinoma (ccRCC): 1) The tumor genome is inferred to be 4N based on the distribution of allelic depths (right) showing four different peaks at equal spacing indicating allelic copy-number states of 0,1,2, and 3; moreover, the presence of many segmental SCNAs at odd copy-number states suggests downstream SCNA evolution after ancestral whole-genome duplication. The sequence of early WGD followed by downstream evolution contrasts with tRCC genomes, as demonstrated in this study, which are usually 2N or show few alterations either before or after WGD. 2) The presence of 3p loss is a hallmark feature of ccRCC but not of tRCCs. Moreover, the 3p loss is coupled to a terminal deletion on 1p accompanied with chromothripsis, a frequent rearrangement pattern seen in ccRCCs (*33*). 3) The clustering of breakpoints on Chrs.4,8,10,11,15, and 17 near the boundaries of large segmental SCNAs is consistent with rearrangement outcomes generated by the breakage of multi-chromosome bridges, including both regional chromothripsis and tandem-short-template rearrangements (*33*, *45*). Although a similar pattern was observed in TRCC18 between chr6 and chr14, such events are rare in tRCCs and we never observed multi-chromosomal (3 or more) rearrangements as seen in this case. **(B)** The frequencies of sSNVs and INDELs in RCC16 are both significantly higher than observed in the remainder of the cohort of tRCCs. **(C)** RCC16 harbors inactivating mutations in prototypical ccRCC tumor suppressors including *VHL*, *SETD2*, *PBRM1*, and *BAP1*, none of which was not detected in the remaining samples in the tRCC cohort. Also listed are the top pathogenic mutations identified in RCC16; notably, a *POLE* mutation with moderate impact was also identified in this case. Full list of variants is in **table S3**.

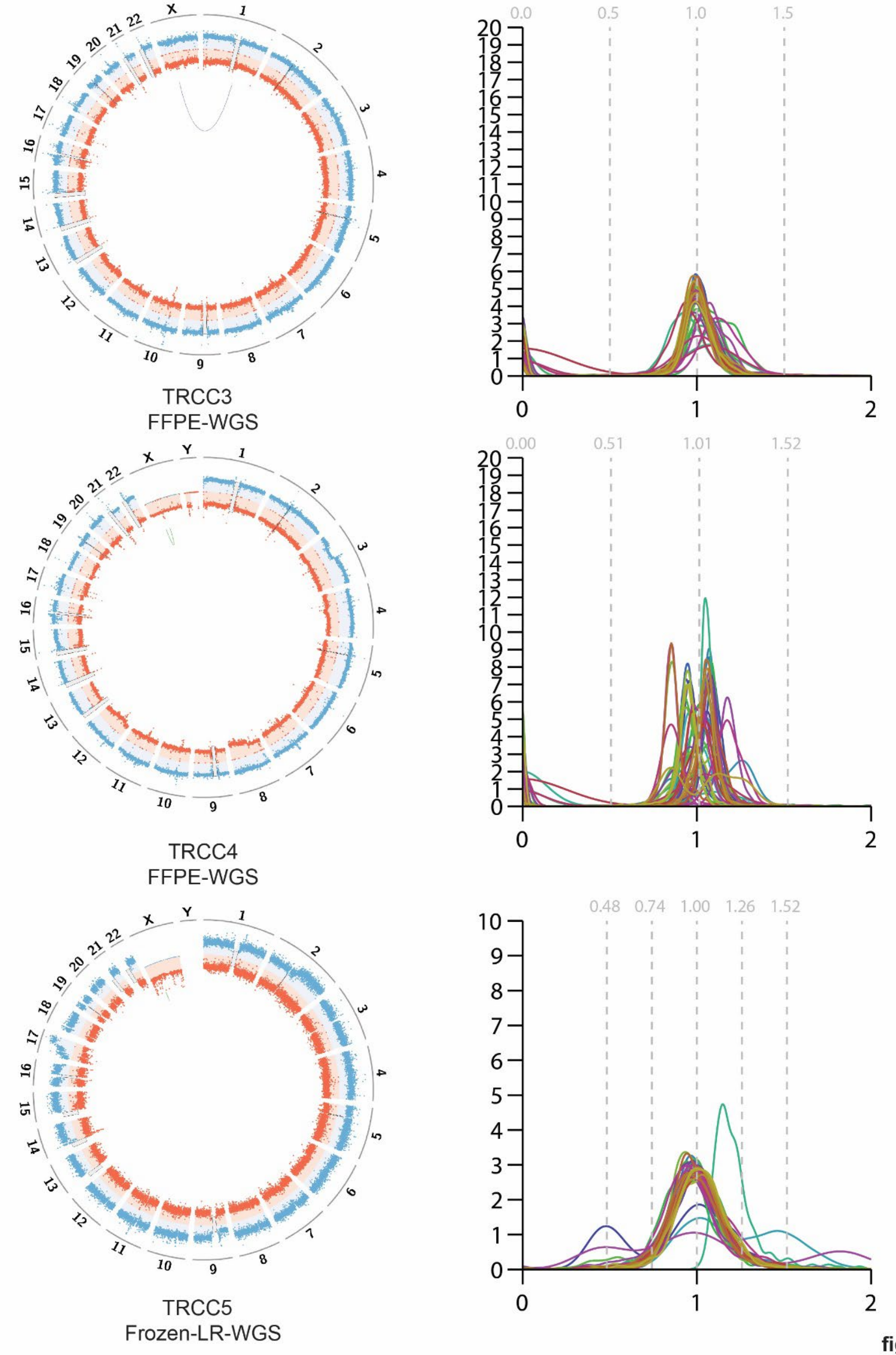
TRCC3
FFPE-WGS
0.0 0.5 1.0 1.5
TRCC4
FFPE-WGS
0.00 0.51 1.01 1.52
TRCC5
Frozen-LR-WGS
0.48 0.74 1.00 1.26 1.52

fig. S4 - 1

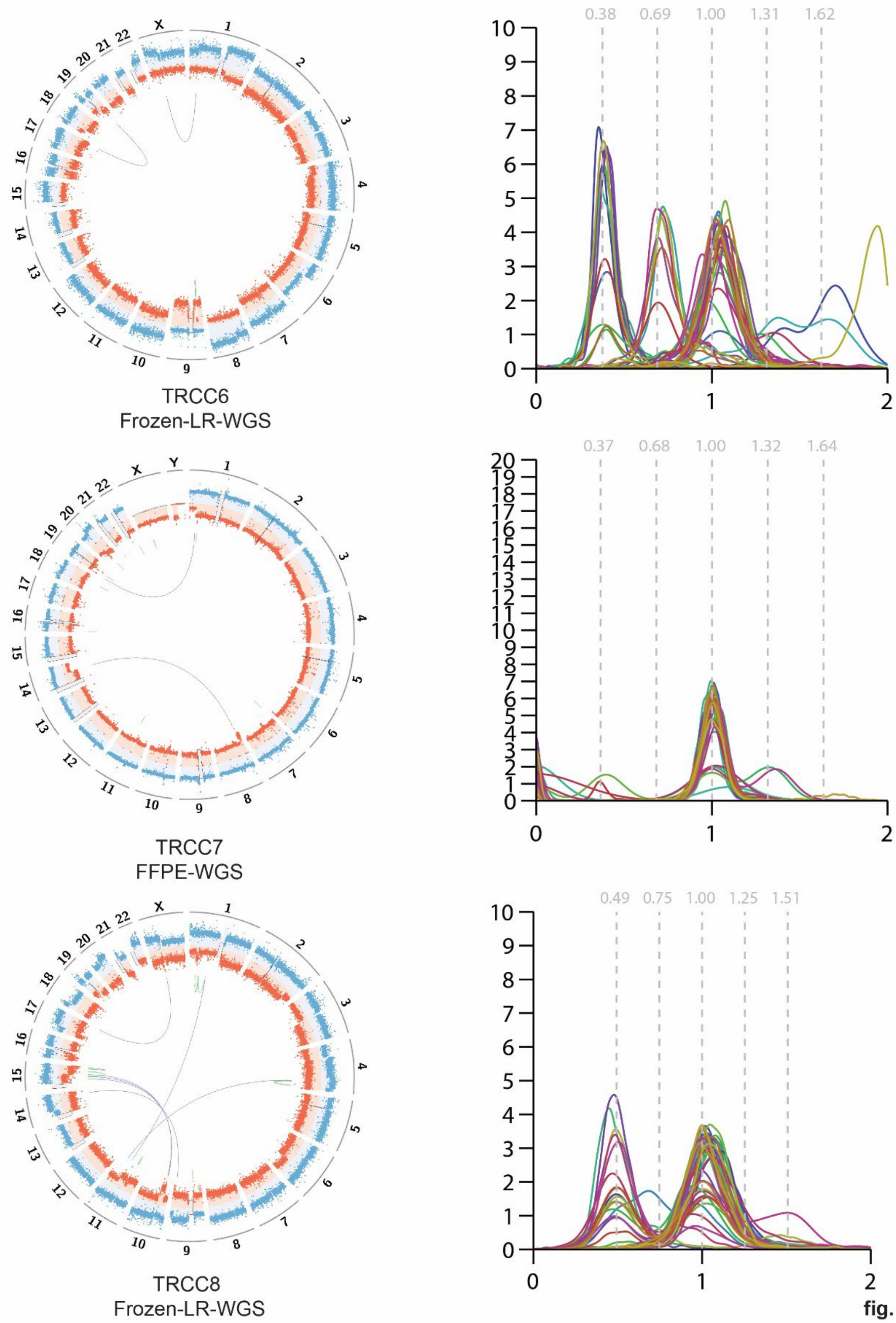
0.38
0.69
1.00
1.31
1.62
TRCC6
Frozen-LR-WGS
0.37
0.68
1.00
1.32
1.64
TRCC7
FFPE-WGS
0.49
0.75
1.00
1.25
1.51
TRCC8
Frozen-LR-WGS

**fig. S4 - 2**

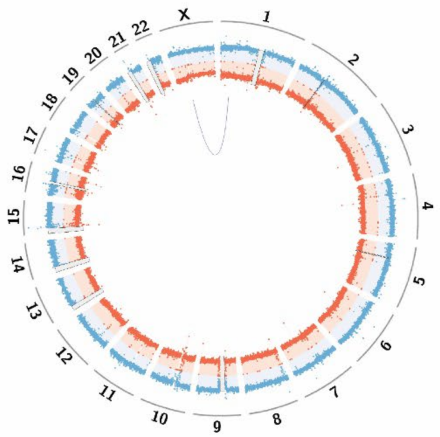


TRCC9
FFPE-WGS

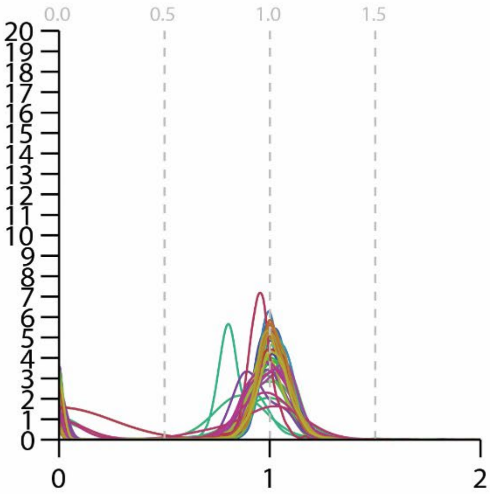


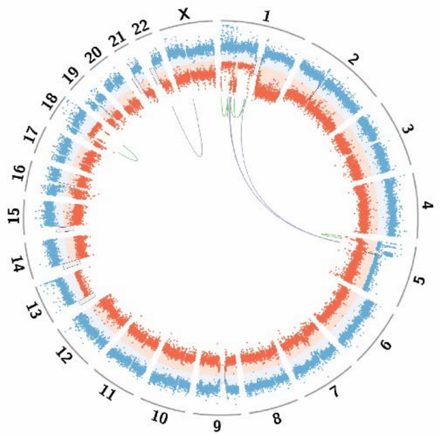


TRCC10
Frozen-LR-WGS

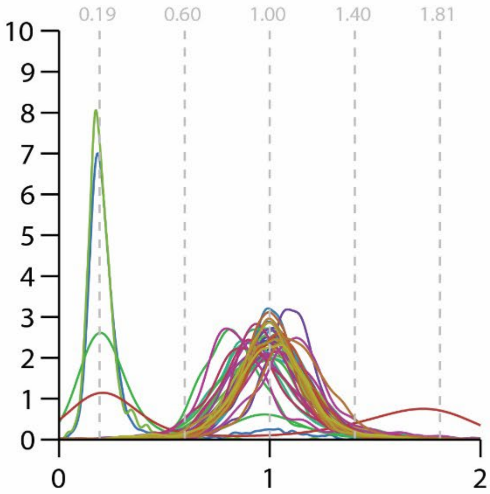


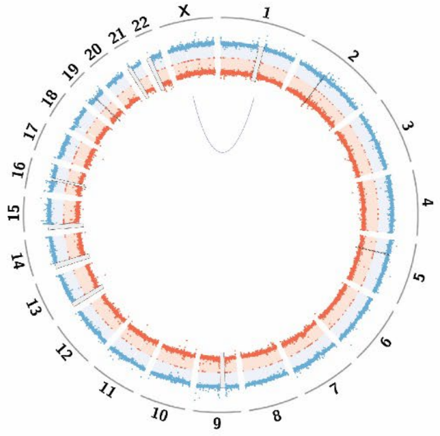


TRCC11
Frozen-WGS

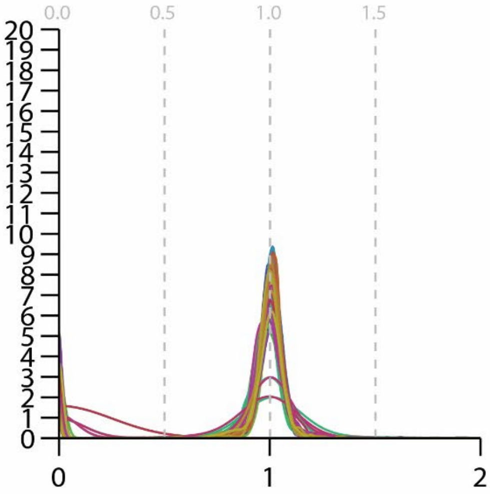


**fig. S4 - 3**

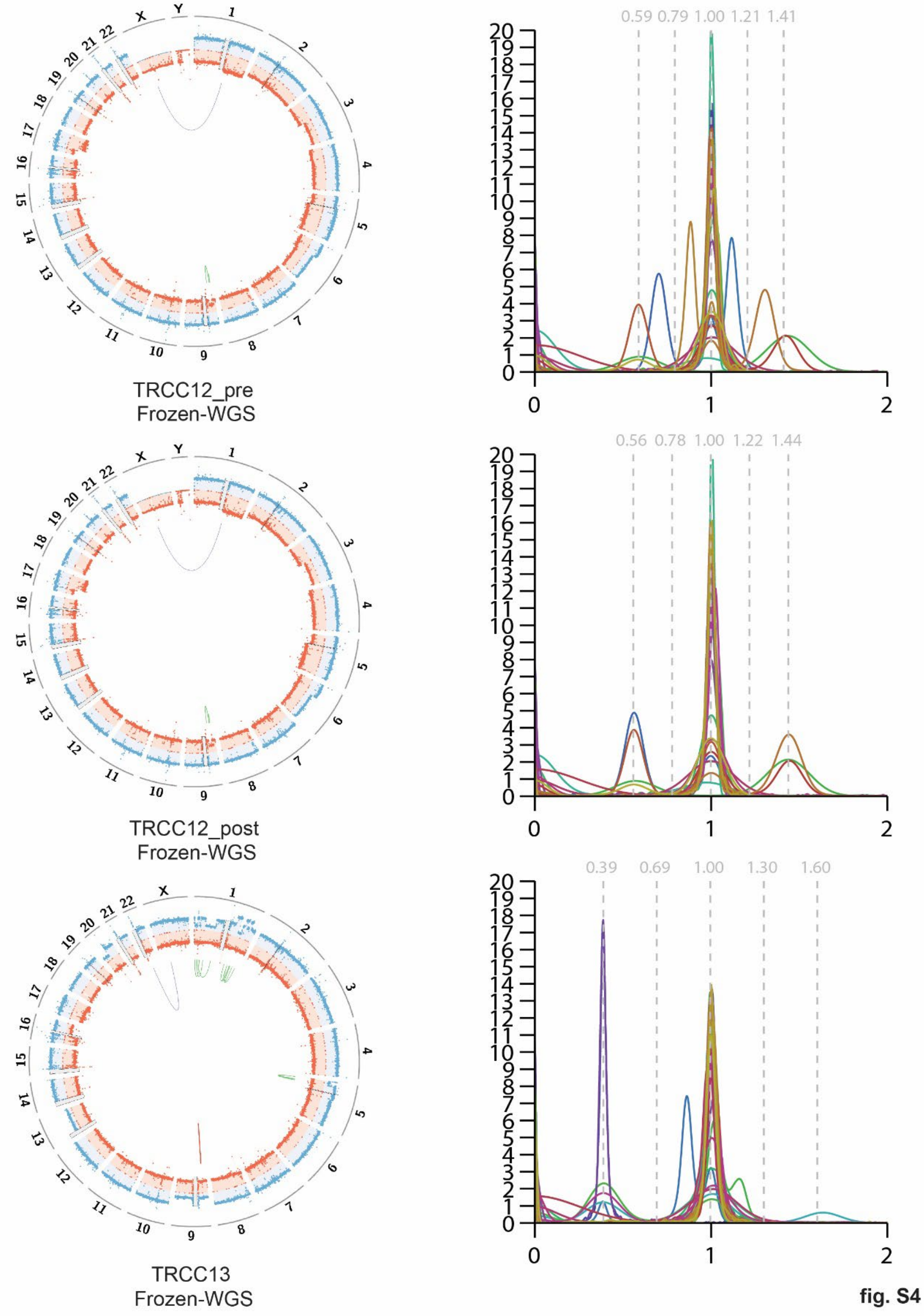

TRCC12_pre
Frozen-WGS
0.59 0.79 1.00 1.21 1.41
TRCC12_post
Frozen-WGS
0.56 0.78 1.00 1.22 1.44
TRCC13
Frozen-WGS
0.39 0.69 1.00 1.30 1.60


fig. S4 - 4

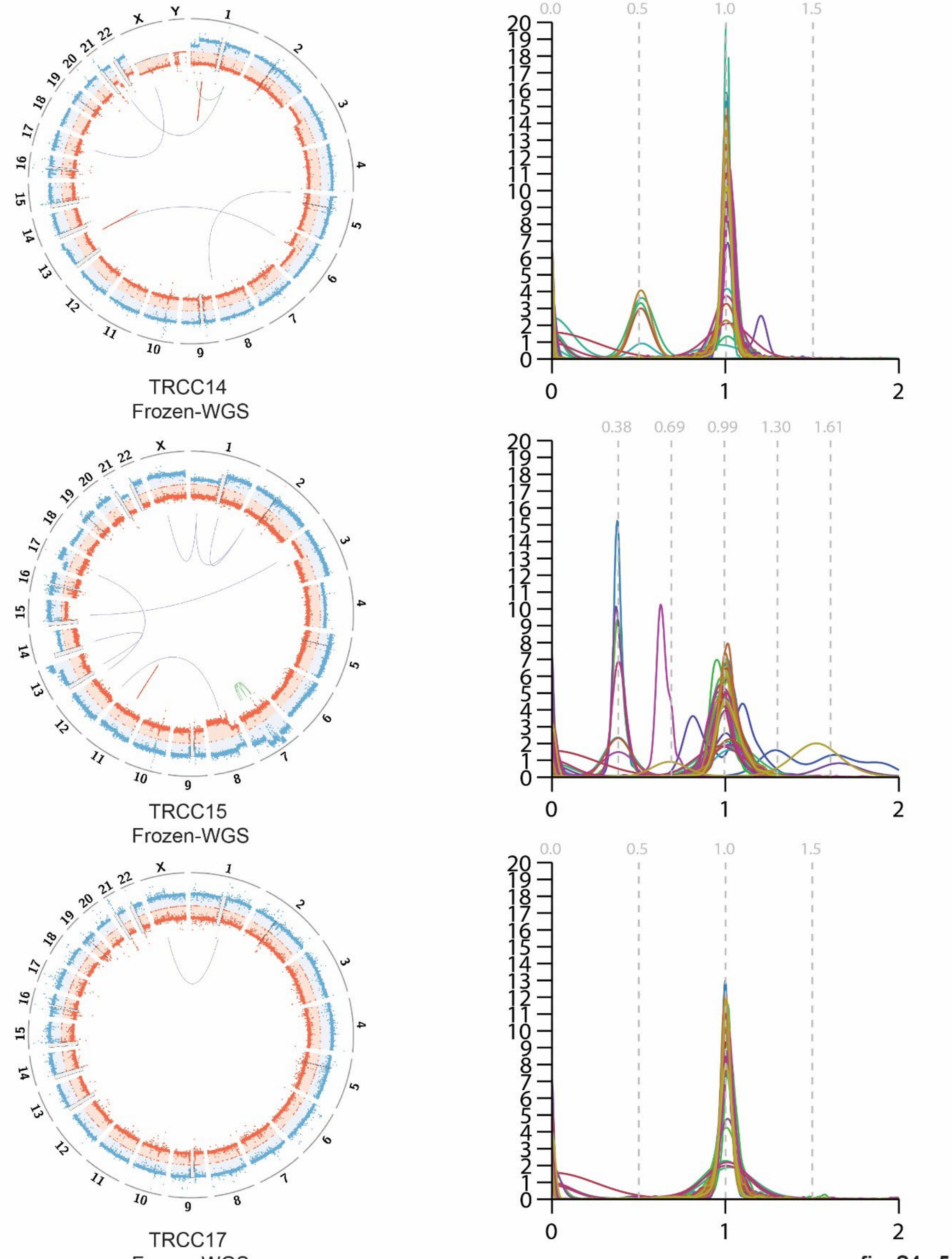
TRCC14
Frozen-WGS
TRCC15
Frozen-WGS
TRCC17
Frozen-WGS
1 2 3 4 5 6 7 8 9 10 11 12 13 14 15 16 17 18 19 20 21 22 X Y
0.0 0.5 1.0 1.5
0.38 0.69 0.99 1.30 1.61
0 1 2 3 4 5 6 7 8 9 10 11 12 13 14 15 16 17 18 19 20
0 1 2

fig. S4 - 5

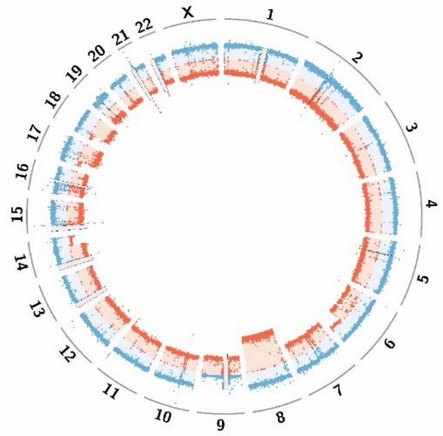


TRCC18 primary tumor
Frozen-WGS

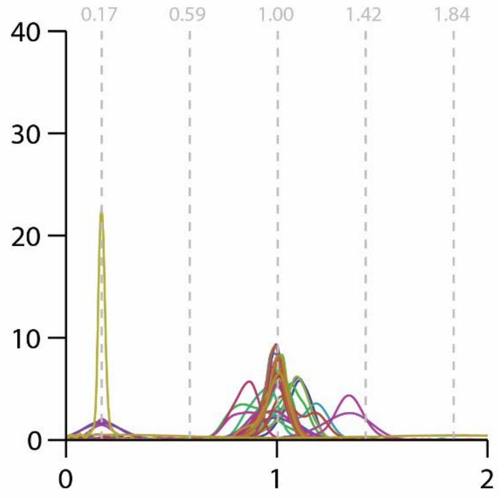


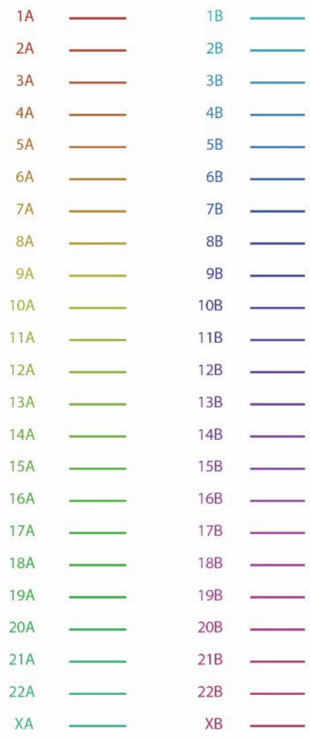


**fig. S4 - 6**

**Fig. S4. Somatic copy number alterations and rearrangements in tRCCs revealed by WGS.** For each case, left column shows CIRCOS plot representing genome-wide DNA copy number profile (red and blue represent coverage of each parental haplotype) and rearrangements. Right column shows the distributions of haplotype-specific sequence coverage of each chromosome (line color scheme by chromosome is on the last page). Note, only the primary is shown for TRCC18 as the metastatic sites are treated separately in **Fig. 3** and **figs. S9-S10**.

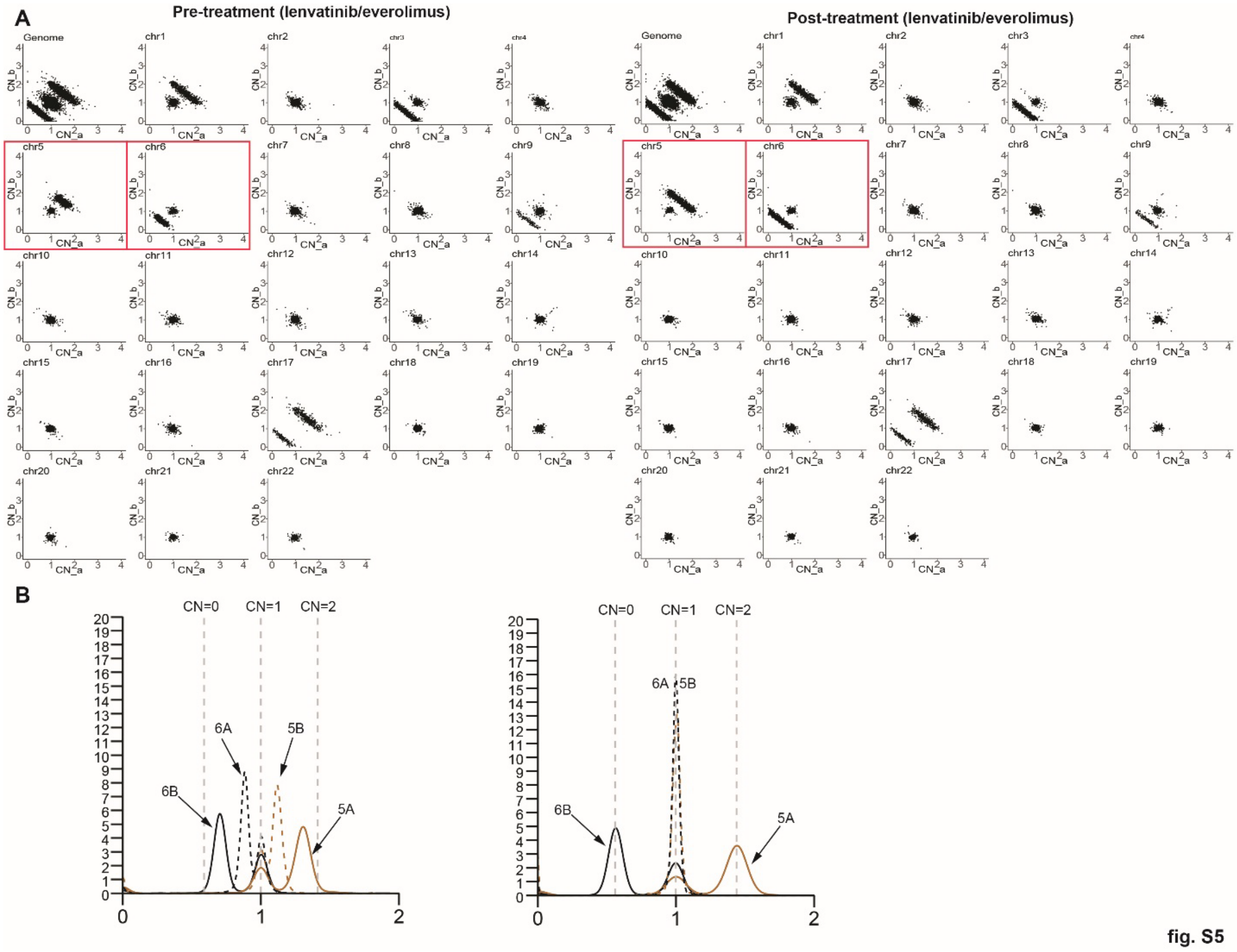


**Fig. S5. Copy-number evolution in pre- and post-treatment metastatic lesions from TRCC12. (A)** Scatterplots of allelic copy number states (calculated via TITAN) within 100 kb bins of all (first panel) and individual chromosomes for the pre- (left) and post-treatment (right) tRCC biopsies from subject TRCC12 before and after progression on lenvatinib/everolimus. Significant copy-number evolution is seen only on chromosomes 5 and 6 (red boxes). **(B)** Haplotype-specific coverage for chromosomes 5 and 6 in the pre- (left) and post-treatment (right) samples. The integer copy-number states were determined from truncal deletions of chr3q and chr17p shared by both samples (data for all chromosomes presented in **fig. S4)**. Allelic DNA copy number of chr5 and chr6 indicates clonal loss of 6q of the B haplotype (6Bq) and gain of 5q of the A haplotype (5Aq) in the post-treatment clone (right), but subclonal losses of 6q and gains of 5q of both haplotypes in the pre-treated clone (left). One possible explanation is that the pre-treatment tumor has two subclonal populations: one containing t(6Bp;5Aq) and 6A but missing 6Bq, the other containing t(6Bq;5Bq) and 6B but missing 6A; and the post-treatment tumor only consists of the first subclone in the primary. Interestingly, both subclones have disomic 6p, monosomic 6q, and trisomic 5q (from both homologs) that match the average DNA copy number of these arms across all tRCCs (**Fig. S2B**).

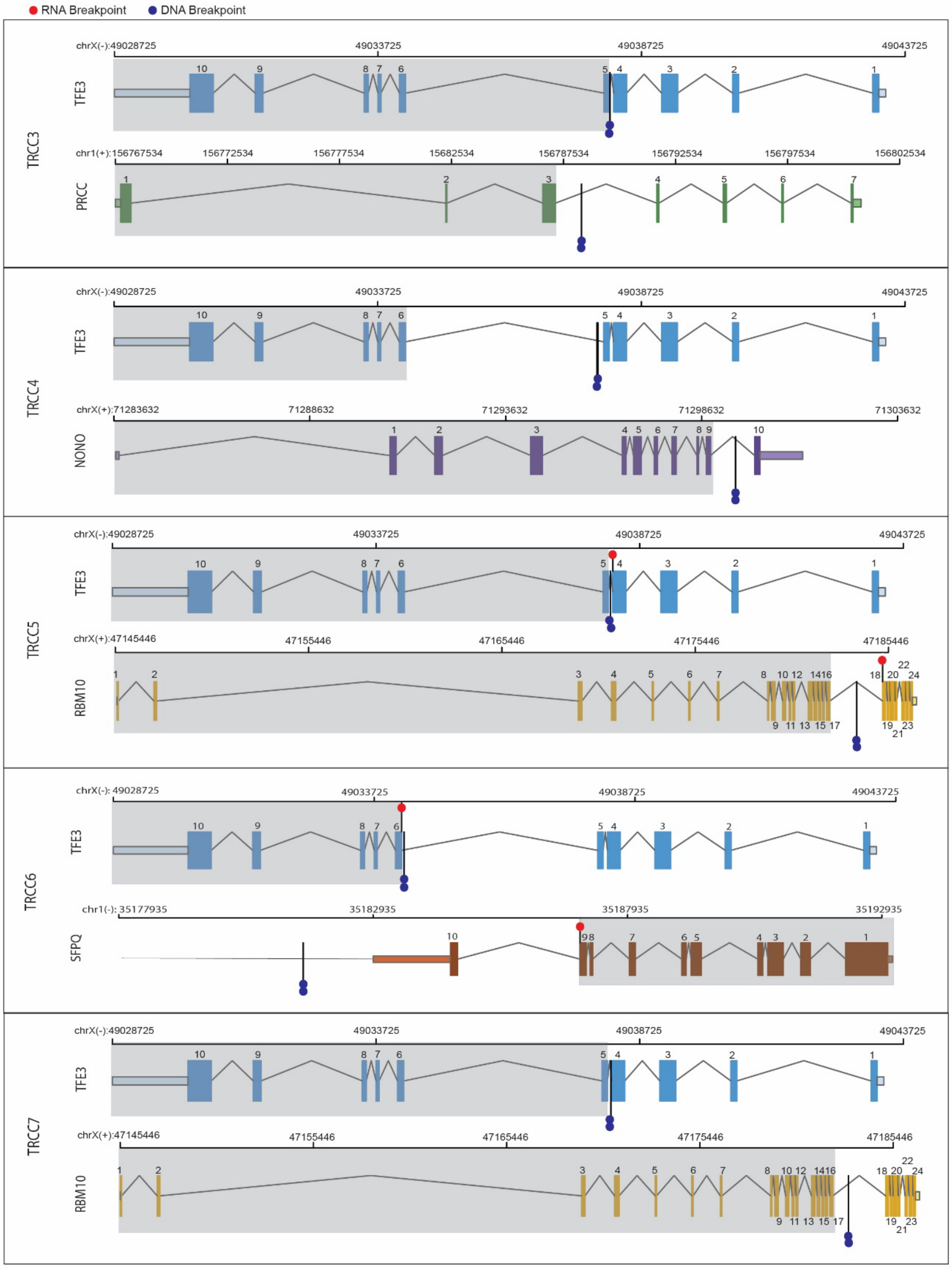
RNA Breakpoint
DNA Breakpoint
TRCC3
TFE3
PRCC
TRCC4
TFE3
NONO
TRCC5
TFE3
RBM10
TRCC6
TFE3
SFPQ
TRCC7
TFE3
RBM10

fig S6-1

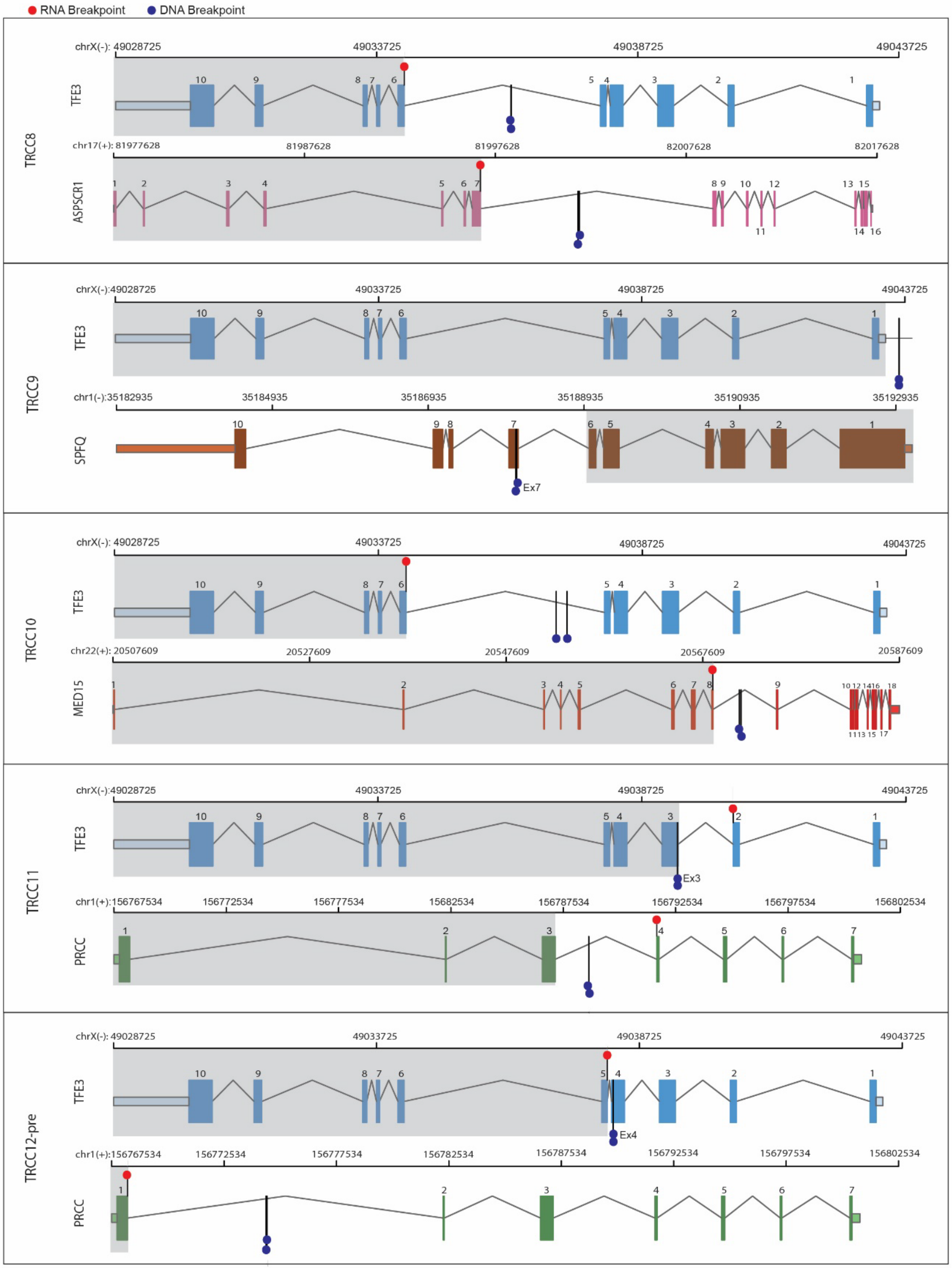
RNA Breakpoint
DNA Breakpoint
TRCC8
TFE3
ASPSCR1
TRCC9
TFE3
SPFQ
Ex7
TRCC10
TFE3
MED15
TRCC11
TFE3
PRCC
Ex3
TRCC12-pre
TFE3
PRCC
Ex4
chrX(-): 49028725
49033725
49038725
49043725
chr17(+): 81977628
81987628
81997628
82007628
82017628
chr1(-):35182935
35184935
35186935
35188935
35190935
35192935
chr22(+): 20507609
20527609
20547609
20567609
20587609
chr1(+):156767534
156772534
156777534
15682534
156787534
156792534
156797534
156802534
156782534

fig S6-2

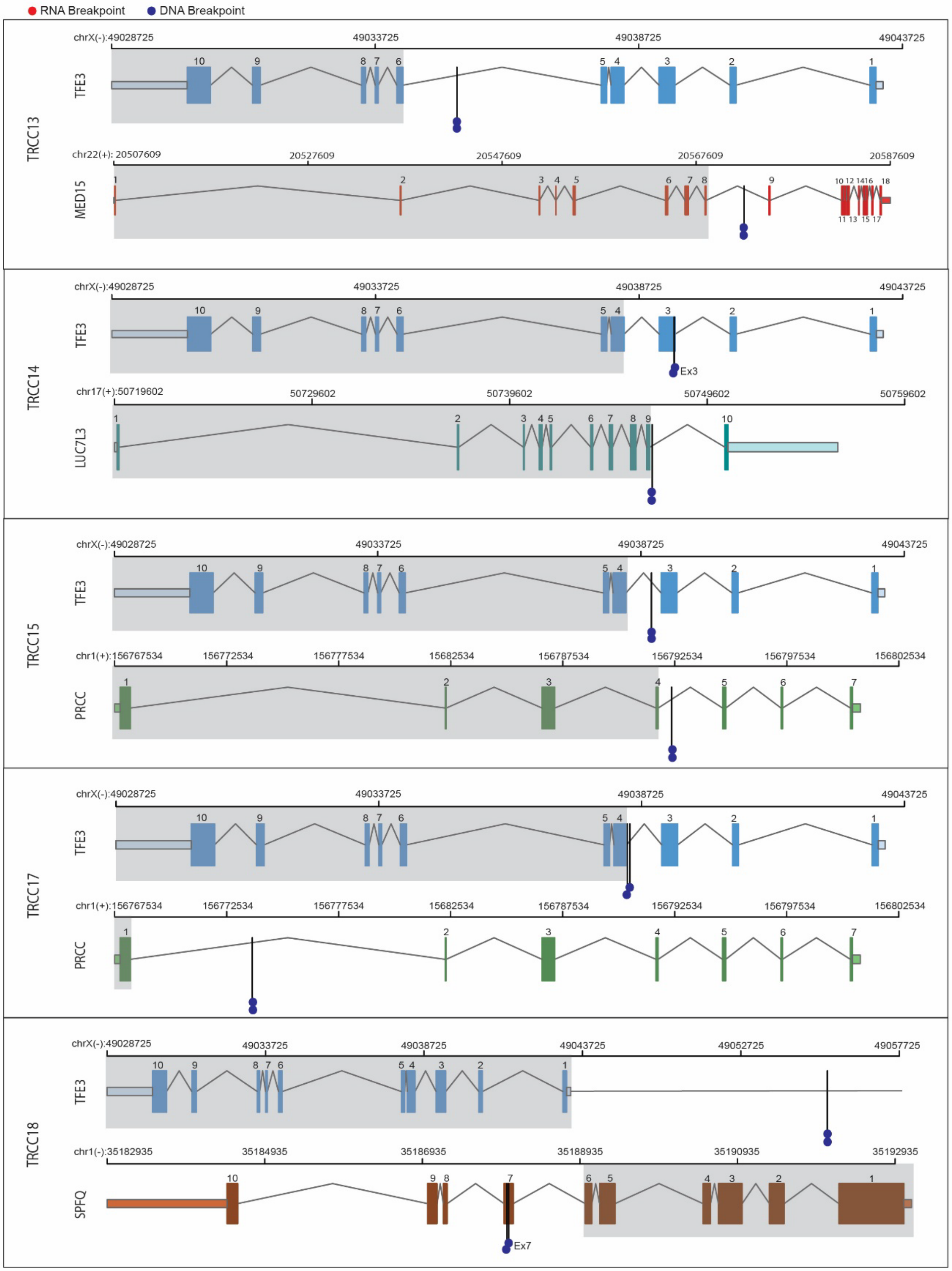
RNA Breakpoint
DNA Breakpoint
TRCC13
TFE3
chrX(-):49028725
49033725
49038725
49043725
MED15
chr22(+): 20507609
20527609
20547609
20567609
20587609
TRCC14
TFE3
Ex3
LUC7L3
chr17(+):50719602
50729602
50739602
50749602
50759602
TRCC15
TFE3
PRCC
chr1(+):156767534
156772534
156777534
15682534
156787534
156792534
156797534
156802534
TRCC17
TFE3
PRCC
TRCC18
TFE3
49052725
49057725
SPFQ
chr1(-):35182935
35184935
35186935
35188935
35190935
35192935
Ex7

fig S6-3

**Fig. S6. Diagrammatic view of *TFE3* fusions identified from DNA and RNA sequencing.** In each sample, the exons in the oncogenic fusion transcripts (Partner-*TFE3*) are highlighted in gray-shaded boxes of the *TFE3* gene (top) and its partner (bottom). DNA breakpoints are marked with vertical lines with dark blue lollipops; RNA breakpoints in the fusion or reciprocal fusion transcripts (in cases where either was determined by STAR-fusion) are marked with vertical lines with red lollipops. The breakpoints in the fusion transcripts could be verified in RNA-Seq data for only a subset of samples (**table S4**).

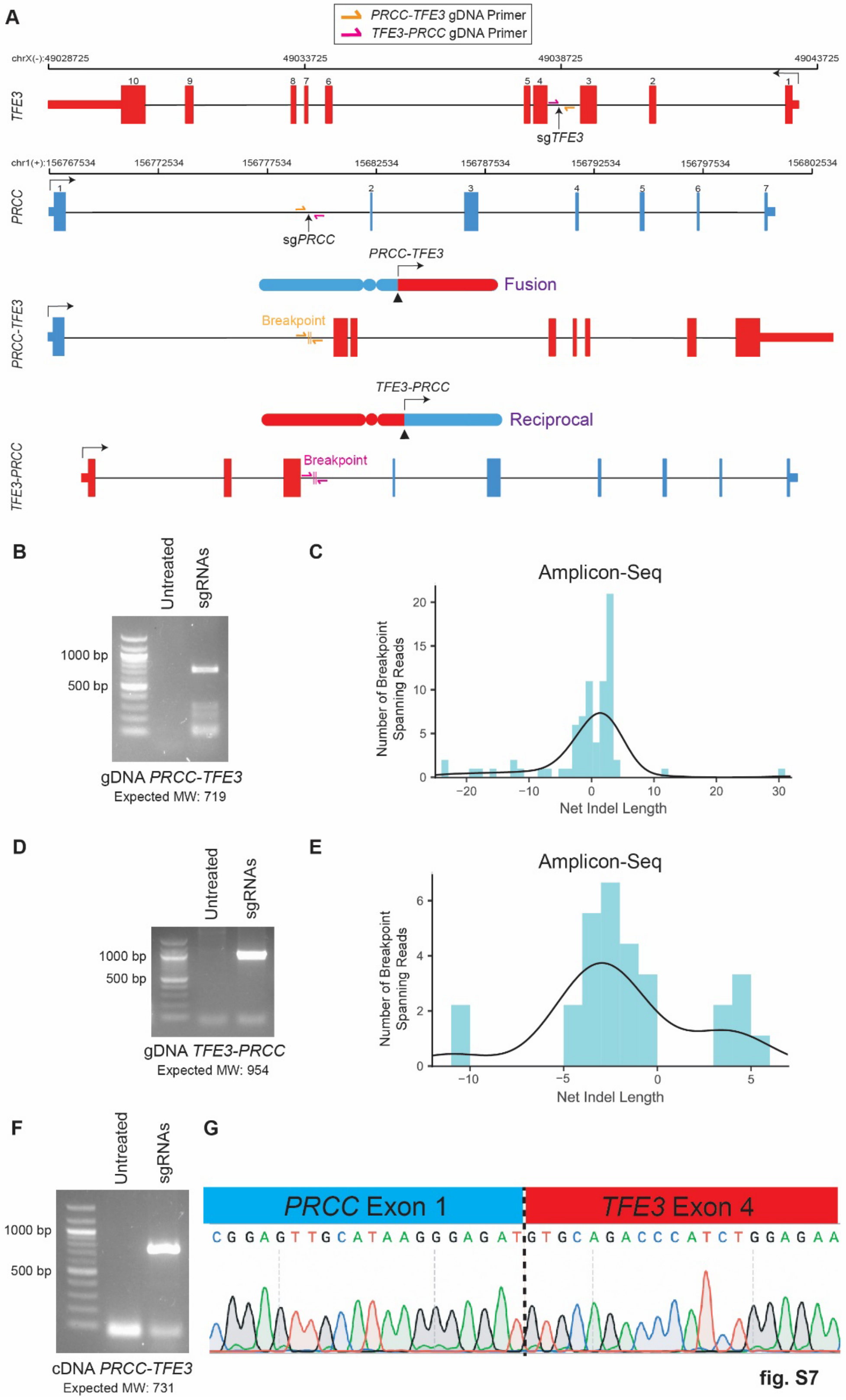
A
PRCC-TFE3 gDNA Primer
TFE3-PRCC gDNA Primer
chrX(-):49028725
49033725
49038725
49043725
TFE3
sgTFE3
chr1(+):156767534
156772534
156777534
15682534
156787534
156792534
156797534
156802534
PRCC
sgPRCC
PRCC-TFE3
Fusion
Breakpoint
TFE3-PRCC
Reciprocal
B
Untreated
sgRNAs
1000 bp
500 bp
gDNA PRCC-TFE3
Expected MW: 719
C
Amplicon-Seq
Number of Breakpoint Spanning Reads
Net Indel Length
D
Untreated
sgRNAs
1000 bp
500 bp
gDNA TFE3-PRCC
Expected MW: 954
E
Amplicon-Seq
Number of Breakpoint Spanning Reads
Net Indel Length
F
Untreated
sgRNAs
1000 bp
500 bp
cDNA PRCC-TFE3
Expected MW: 731
G
PRCC Exon 1
TFE3 Exon 4
CGGAGTTGCATAAGGGAGATGTGCAGACCCATCTGGAGAA

fig. S7

**Fig. S7. *In vitro* engineering of *TFE3* fusions via CRISPR/Cas9 recapitulates WGS findings. (A)** Genomic locations of sgRNAs used to engineer *PRCC-TFE3* fusions via CRISPR/Cas9 and PCR primers for recovering both *PRCC-TFE3* and *TFE3-PRCC* fusions. **(B)** DNA gel showing *PRCC-TFE3* fusions in the bulk edited population of HEK293 cells. **(C)** Distribution of the number of deleted or inserted nucleotides in each PCR amplicon relative to the predominant *PRCC-TFE3* breakpoint (chr1:156777745-chrX:49038604) inferred from the population. Density distribution is overlayed. **(D)** DNA gel showing *TFE3-PRCC* fusion in the edited bulk population. **(E)** Distribution of insertions/deletions relative to inferred *TFE3-PRCC* breakpoint (chrX:49038605-chr1:156777746). Density distribution is overlayed. Two amplicons displayed insertions of sequences of Cas9 DNA (~75bp) and were excluded. **(F)** Expressed *PRCC-TFE3* fusion transcripts detected via RT-PCR across the predicted RNA fusion breakpoint in the edited bulk population. **(G)** Sanger sequencing trace confirming *PRCC-TFE3* RNA breakpoint.

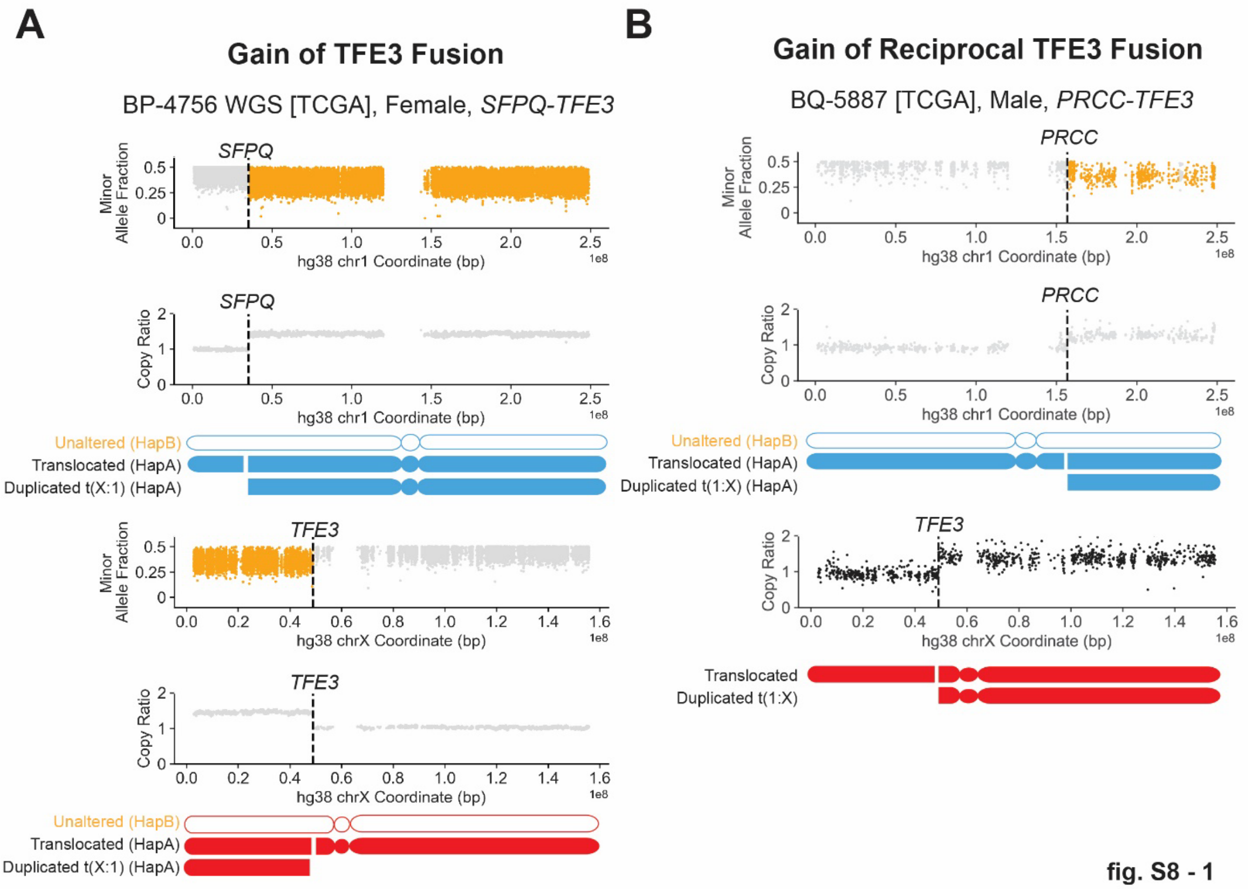
A
Gain of TFE3 Fusion
BP-4756 WGS [TCGA], Female, SFPQ-TFE3
SFPQ
Minor Allele Fraction
hg38 chr1 Coordinate (bp)
Copy Ratio
Unaltered (HapB)
Translocated (HapA)
Duplicated t(X:1) (HapA)
TFE3
hg38 chrX Coordinate (bp)
B
Gain of Reciprocal TFE3 Fusion
BQ-5887 [TCGA], Male, PRCC-TFE3
PRCC
Duplicated t(1:X) (HapA)
Translocated
Duplicated t(1:X)

**fig. S8 - 1**

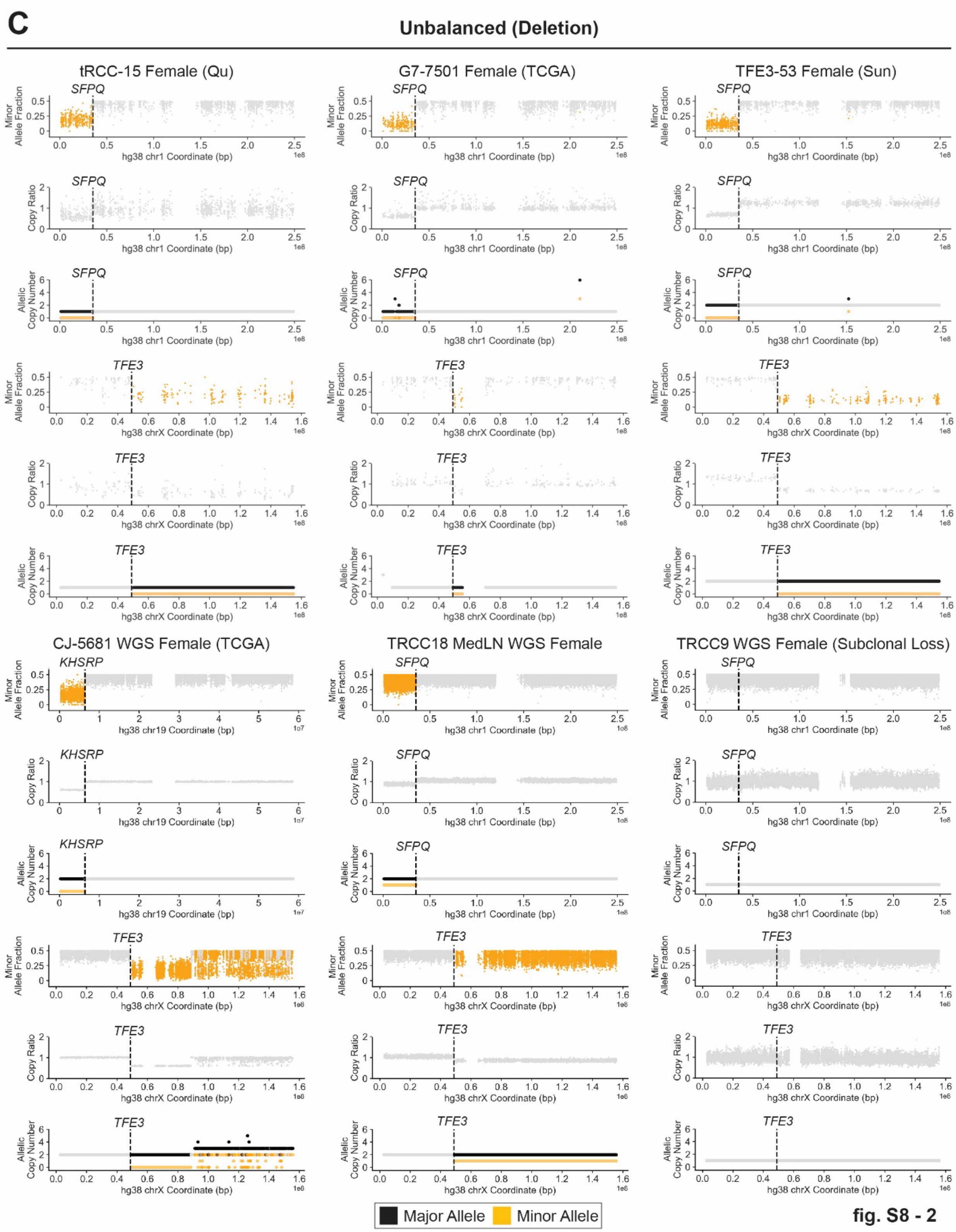
C
Unbalanced (Deletion)
tRCC-15 Female (Qu)
G7-7501 Female (TCGA)
TFE3-53 Female (Sun)
CJ-5681 WGS Female (TCGA)
TRCC18 MedLN WGS Female
TRCC9 WGS Female (Subclonal Loss)
SFPQ
TFE3
KHSRP
Minor Allele Fraction
Copy Ratio
Allelic Copy Number
hg38 chr1 Coordinate (bp)
hg38 chrX Coordinate (bp)
hg38 chr19 Coordinate (bp)
Major Allele
Minor Allele

fig. S8 - 2

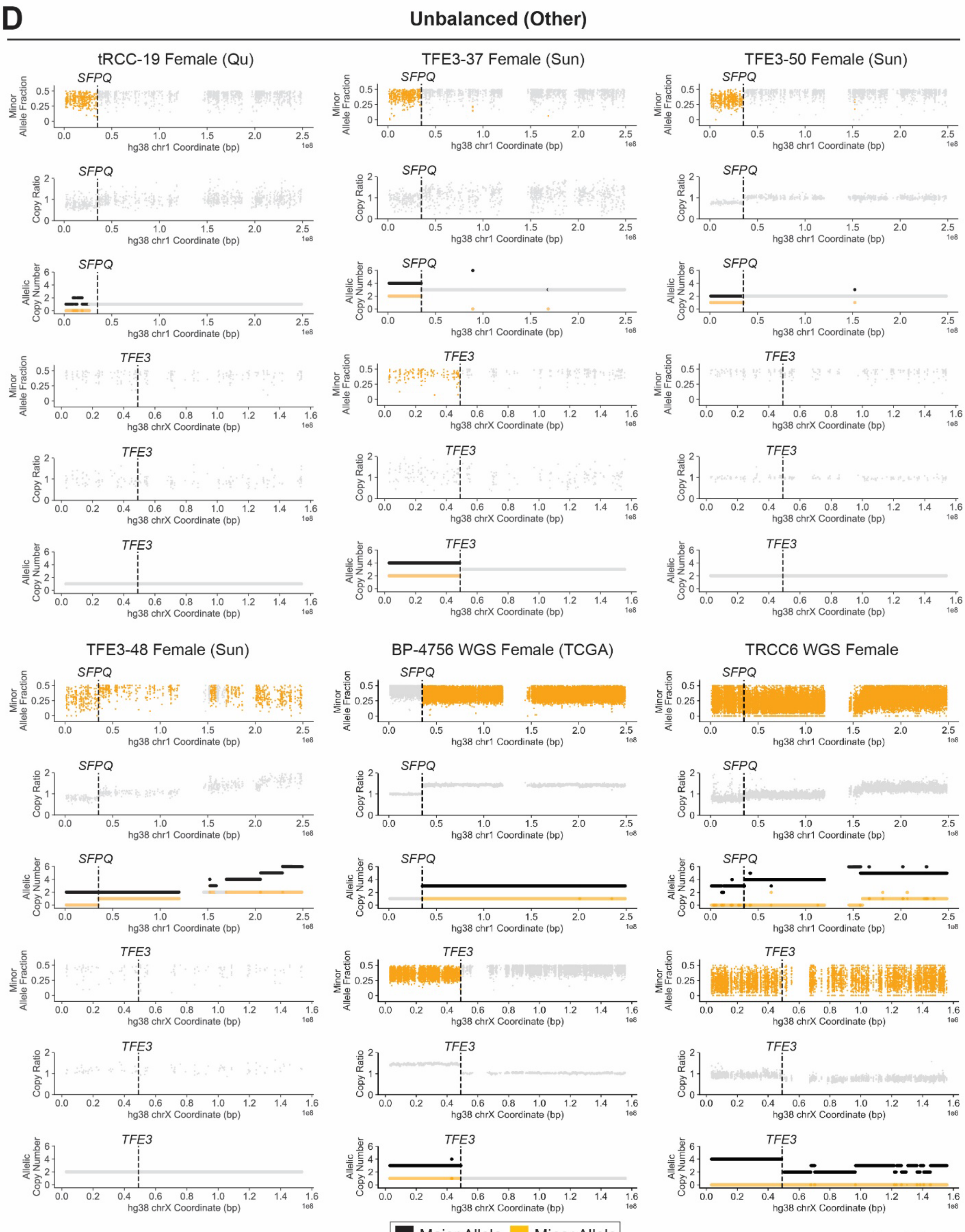
D
Unbalanced (Other)
tRCC-19 Female (Qu)
TFE3-37 Female (Sun)
TFE3-50 Female (Sun)
TFE3-48 Female (Sun)
BP-4756 WGS Female (TCGA)
TRCC6 WGS Female
SFPQ
TFE3
Minor Allele Fraction
Copy Ratio
Allelic Copy Number
hg38 chr1 Coordinate (bp)
hg38 chrX Coordinate (bp)
1e8
Major Allele
Minor Allele

fig. S8 - 3

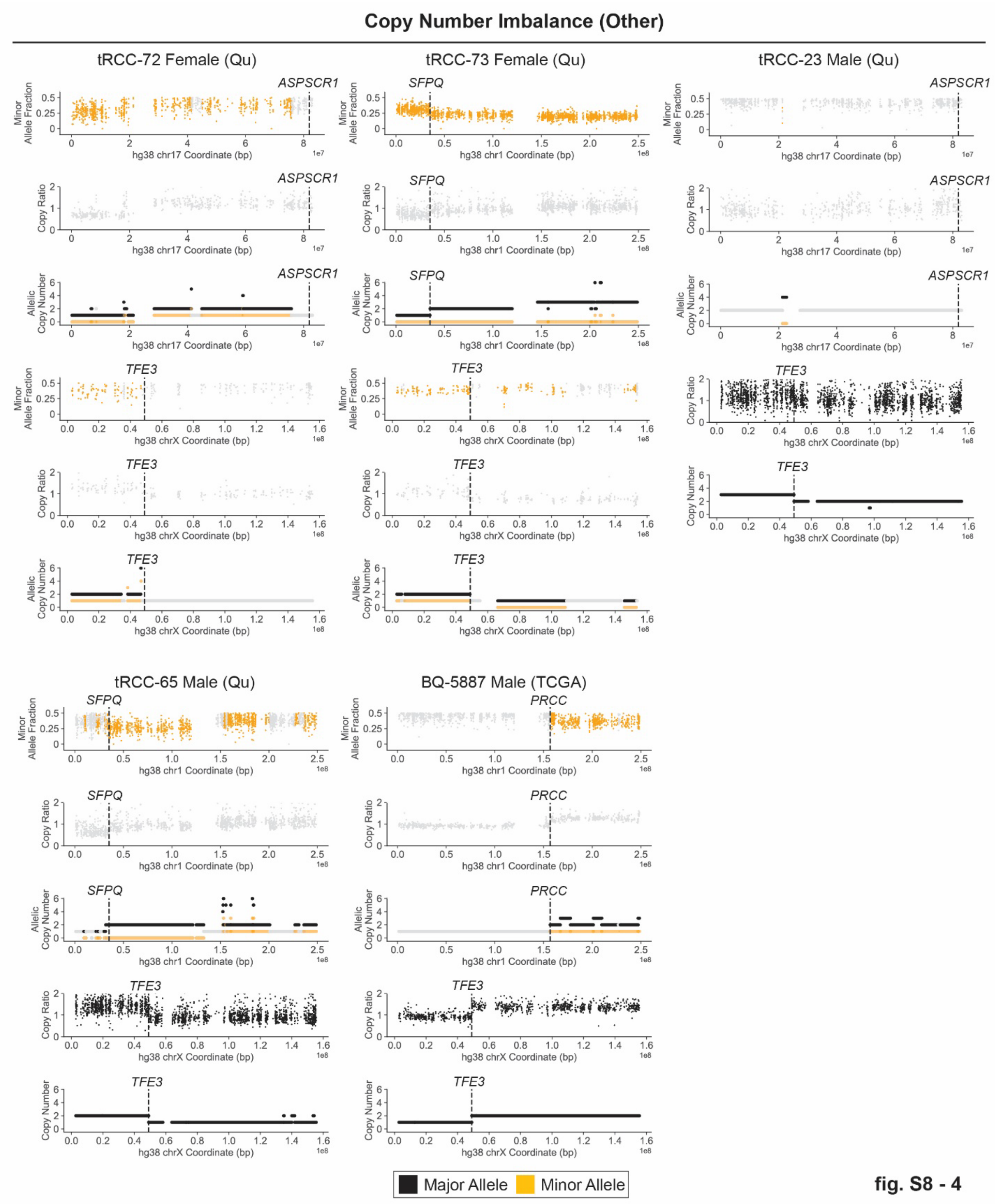

Copy Number Imbalance (Other)
tRCC-72 Female (Qu)
tRCC-73 Female (Qu)
tRCC-23 Male (Qu)
tRCC-65 Male (Qu)
BQ-5887 Male (TCGA)
ASPSCR1
SFPQ
TFE3
PRCC
Minor Allele Fraction
Copy Ratio
Allelic Copy Number
Copy Number
hg38 chr17 Coordinate (bp)
hg38 chr1 Coordinate (bp)
hg38 chrX Coordinate (bp)
1e7
1e8
Major Allele
Minor Allele


fig. S8 - 4

**Fig. S8. Copy Number Imbalance of *TFE3* translocations. (A)** Minor allele fraction (top) and read depth ratio (tumor/normal) from the whole-genome sequencing data of a tumor (TCGA BP-4756, female, *SFPQ-TFE3*) showing copy-number gain of the translocated chromosome t(X;1) containing the oncogenic *TFE3* fusion. Regions with copy number imbalance are highlighted in orange. **(B)** Minor allele fraction and read-depth ratio from whole-exome sequencing data of a tRCC tumor (TCGA BQ-5887, male, *PRCC-TFE3*) showing copy-number gain of the translocated chromosome with the reciprocal *TFE3* fusion. Regions with copy number imbalance are highlighted in orange. **(C)** tRCC samples in the aggregate cohort showing deletion (DNA copy number lower than the basal copy-number state across the genome) of the reciprocal *TFE3* fusion. Deletion/loss of the *TFE3* fusion was never observed. Shown for each sample are the minor allele fractions, read-depth ratios, and segmented allelic copy number of both chrX and the partner chromosome. The allelic DNA copy-number of these external cohorts was calculated by TITAN (*76*), which does not perform local haplotype inference. **(D)** tRCC samples in the aggregate cohort showing copy-number imbalance across translocation breakpoints at either *TFE3* or its partner locus. Copy number plots are as described in (C). For cases with copy-number imbalance on one breakpoint but not the other, a plausible explanation is that the *TFE3* fusions arose from chromoplexy events (multi-way balanced translocations) as shown in **Fig. 1E** (TRCC15), with copy-number imbalance generated by subsequent chromosomal losses. Although direct validation of this explanation is not possible without whole-genome sequencing, we identified 4 potential chromoplexy cases in WES cohorts, suggesting that chromoplexy may contribute to a small fraction of *TFE3* fusions.

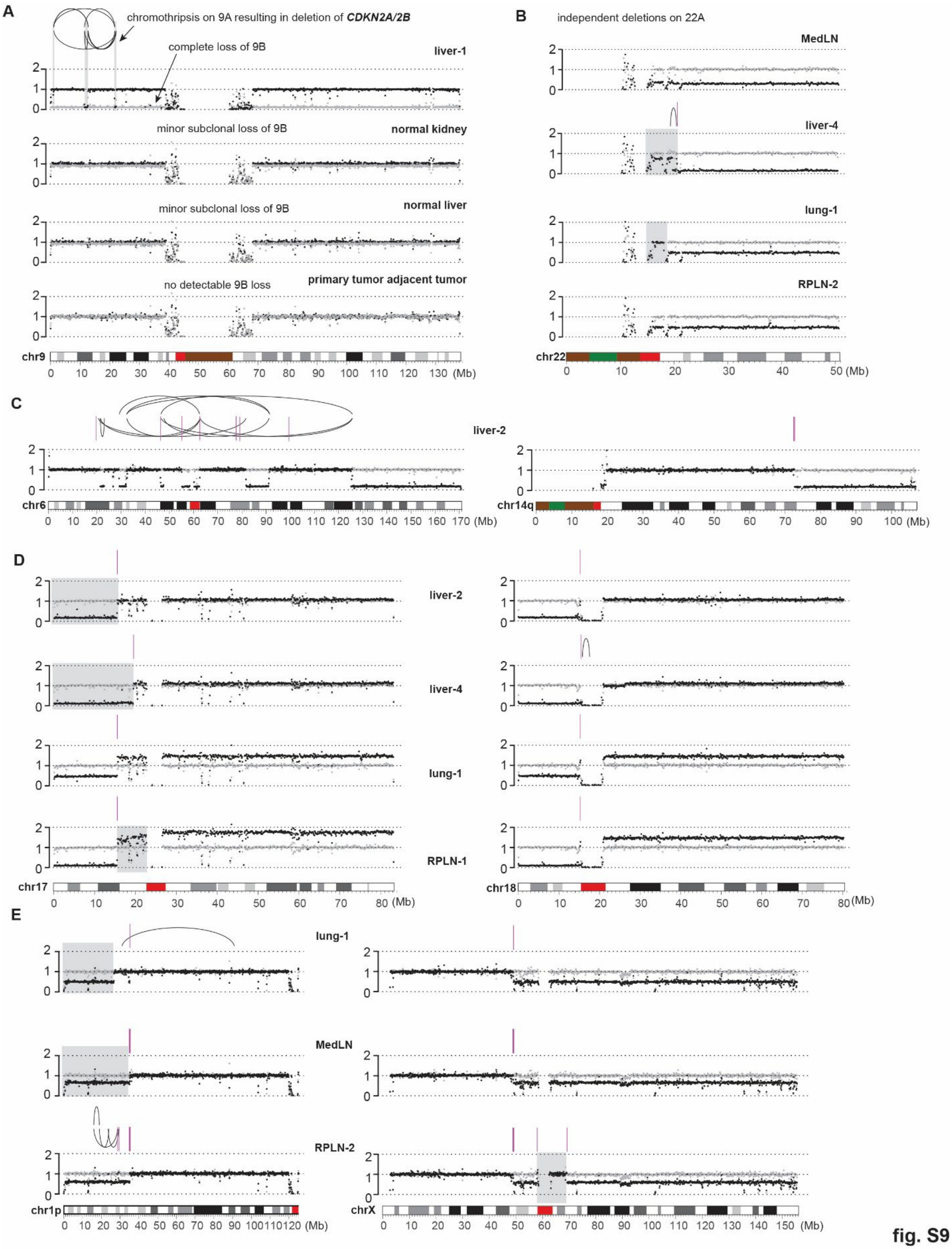
A
chromothripsis on 9A resulting in deletion of CDKN2A/2B
complete loss of 9B
liver-1
minor subclonal loss of 9B
normal kidney
minor subclonal loss of 9B
normal liver
no detectable 9B loss
primary tumor adjacent tumor
chr9
B
independent deletions on 22A
MedLN
liver-4
lung-1
RPLN-2
chr22
C
liver-2
chr6
chr14q
D
liver-2
liver-4
lung-1
RPLN-1
chr17
chr18
E
lung-1
MedLN
RPLN-2
chr1p
chrX
(Mb)

fig. S9

**Fig. S9. Examples of SCNAs and SCNA evolution in TRCC18. (A)** Haplotype-specific DNA copy number (gray and black dots, 100kb intervals) and rearrangements of chr9. Shown are the data for the liver-1 metastatic lesion (representative of ancestral alterations of the founding tumor clone), the normal kidney and normal liver samples taken at rapid autopsy showing minor chr9 losses indicating the presence of tumor cells (gray dots lower than the black), and the normal adjacent kidney tissue sampled along with the primary tumor at the time of nephrectomy showing complete allelic balance (gray dots overlaying on top of black dots). We therefore excluded the two normal autopsy samples from mutation analysis used for phylogenetic inference due to the presence of contaminating tumor cells. **(B)** Independent deletions of chr22q detected in the MedLN sample (complete deletion), Liver-4 (terminal deletion), Lung-1 (terminal deletion with a different boundary), and RPLN-2 (complete deletion). These deletions are absent in the remaining liver autopsies and have a low clonal fraction in the primary. Interestingly, all the deletions occurred to the same 22 homolog; we cannot rule out the possibility that these deletions may have arisen from progressive deletions of an unstable chr22q and therefore were related. **(C)** Chromothripsis of chr6 that is linked to a terminal deletion on 14q, both of which inferred to be present in the founding tumor clone (see **Fig. 3**). The clustering of breakpoints on chr6 is very similar to the clustering of breakpoints seen in RCC16 and in our prior studies of breakage-fusion-bridge cycles(*45*); it is highly plausible that the rearrangements of both chr6 and chr14 arose from micronuclei or chromosome bridges. This pattern contrasts with the simple outcome of reciprocal translocations associated with *TFE3* fusions between chr1 and chrX. **(D)** Large segmental deletions on 17p (left) and 18p (right) inferred to be present in the founding tumor clone. The same breakpoints on 17p and 18p are seen in Liver-2 (and other Liver metastases except Liver-4), Lung metastases, and RPLN-1, but there is copy-number gain of the translocated chromosome t(18q,17q) in different metastases that may be due to positive selection; the Liver-4/MedLN lesions show a larger deletion on 17p, indicating downstream evolution of the translocated chromosome. Notably, there is sloping copy-number variation near the 17p terminus (black) in RPLN-1 that is absent in the intact homolog (gray); this pattern suggests ongoing evolution of the translocated chromosome that is consistent with varying copy-number states and breakpoints seen in the different metastatic samples. **(E)** Independent deletions of t(1;X) in the lung metastases (using Lung-1 as representative), MedLN/Liver-4 (using MedLN as representative), and RPLN-2 samples. In the Lung samples, the deletion spanned the entire segment from chrX but left a small segment on chr1 intact, although there are focal deletions near the translocation site that deleted the reciprocal translocation. In the MedLN/Liver-4 samples, the loss spanned the entire t(1;X) chromosome but occurred after whole-genome duplication; therefore, this must be independent from the deletion in Lung metastases. Finally, the deletion in RPLN-2 resulted from chromothripsis of the t(1;X) chromosome, with partial retention of the Xq and complex rearrangements between 1p and X; this event again has to be independent from MedLN/Liver-4, since it is inferred to be clonal or have occurred before WGD; it is also independent from the deletion in Lung metastases due to the presence of segments with complementary deletion/retention. It is possible that the deletions in Lung metastases and in RPLN-2 both had descended from an ancestor with an unstable chromosome generated by chromothripsis/DNA fragmentation; this possibility cannot be directly tested given the divergence between the RPLN-2 metastasis and the Lung metastases and also does not pose a conflict to the phylogenetic inference.

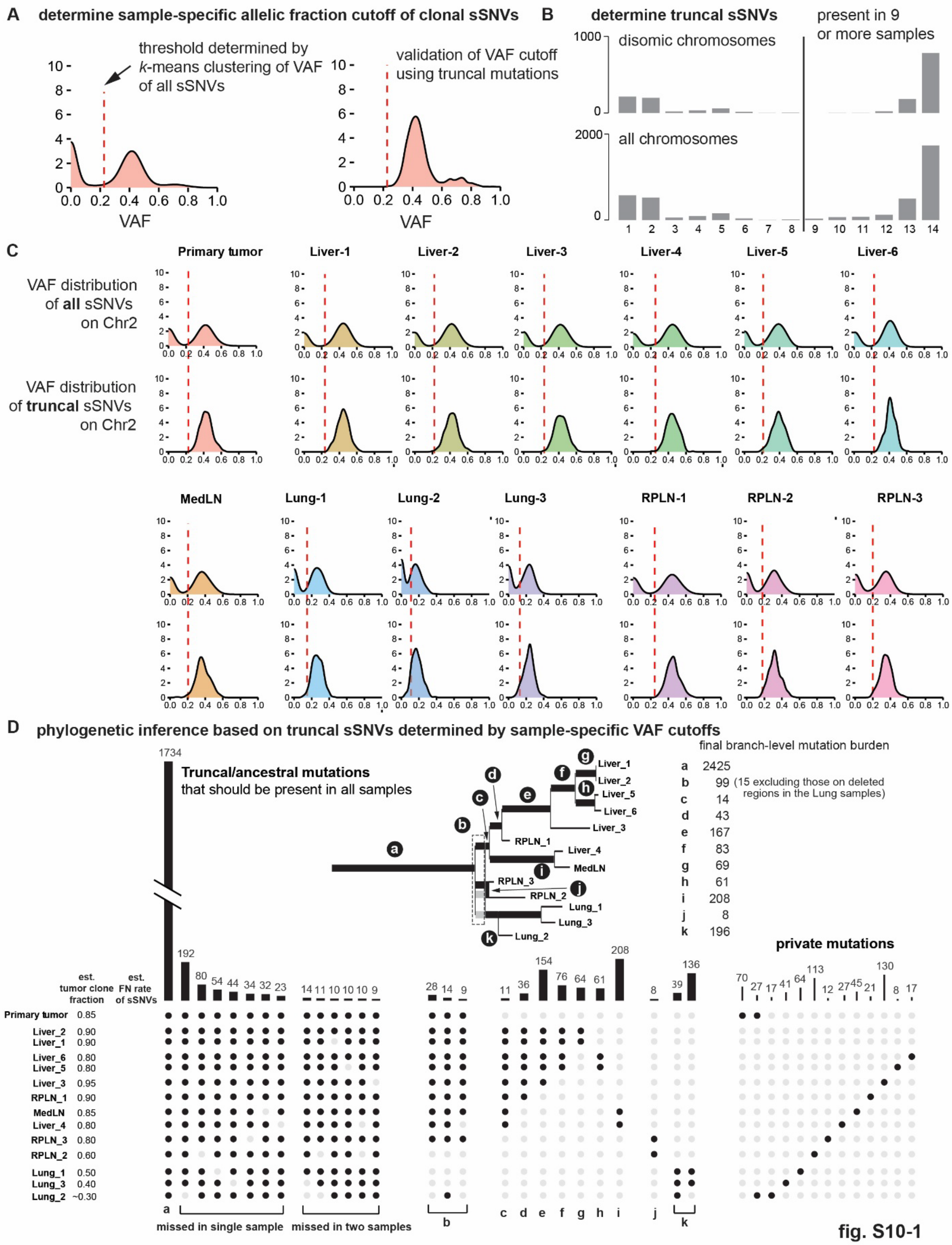
A determine sample-specific allelic fraction cutoff of clonal sSNVs
threshold determined by k-means clustering of VAF of all sSNVs
validation of VAF cutoff using truncal mutations
VAF
B determine truncal sSNVs
disomic chromosomes
all chromosomes
present in 9 or more samples
C
VAF distribution of all sSNVs on Chr2
VAF distribution of truncal sSNVs on Chr2
Primary tumor
Liver-1
Liver-2
Liver-3
Liver-4
Liver-5
Liver-6
MedLN
Lung-1
Lung-2
Lung-3
RPLN-1
RPLN-2
RPLN-3
D phylogenetic inference based on truncal sSNVs determined by sample-specific VAF cutoffs
Truncal/ancestral mutations that should be present in all samples
final branch-level mutation burden
a 2425
b 99 (15 excluding those on deleted regions in the Lung samples)
c 14
d 43
e 167
f 83
g 69
h 61
i 208
j 8
k 196
private mutations
est. tumor clone fraction
est. FN rate of sSNVs
Primary tumor 0.85
Liver_2 0.90
Liver_1 0.90
Liver_6 0.80
Liver_5 0.80
Liver_3 0.95
RPLN_1 0.90
MedLN 0.85
Liver_4 0.80
RPLN_3 0.80
RPLN_2 0.60
Lung_1 0.50
Lung_3 0.40
Lung_2 ~0.30
missed in single sample
missed in two samples

fig. S10-1

E Phasing of somatic mutations on chrX based on VAFs in different tumors

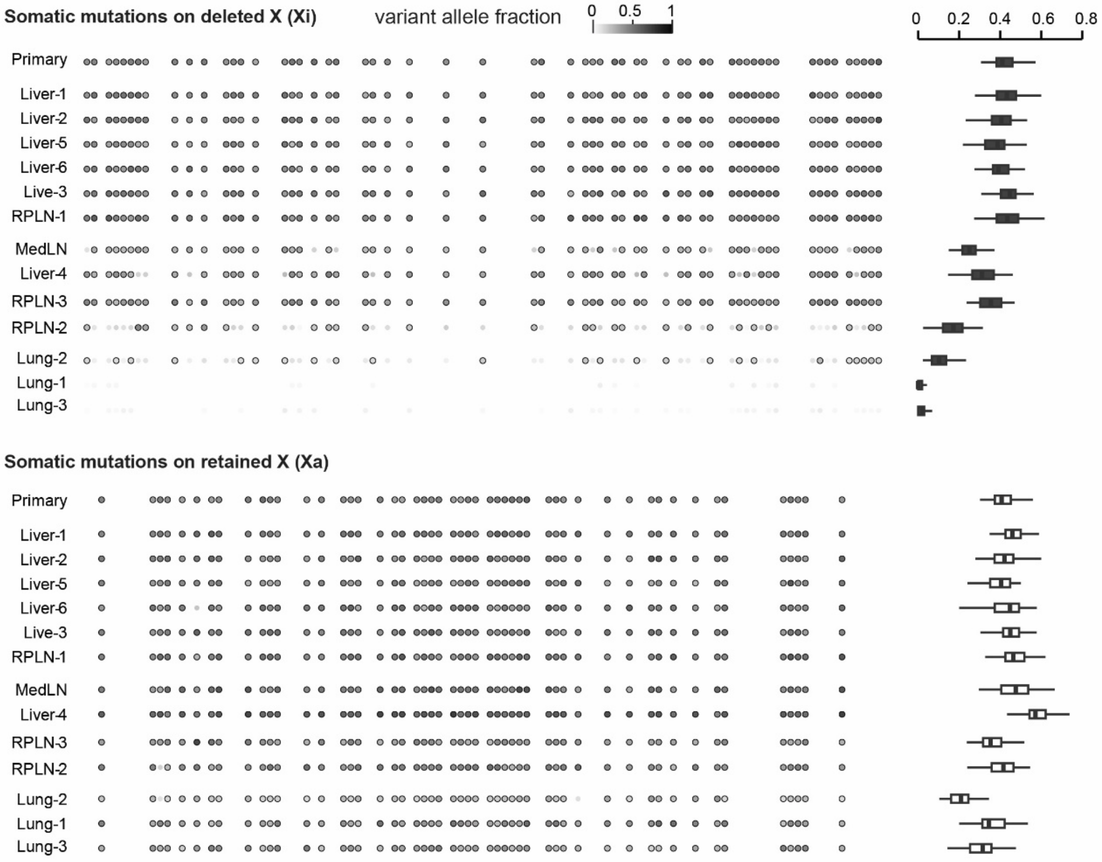


fig. S10-2

**Fig. S10. Phylogenetic trees of TRCC18 metastases constructed from somatic mutations. (A)** As the tumor samples may contain minor subclones either from within the tumor, or due to handling of multiple autopsy samples taken simultaneously, we first determined the minimum variant allelic fractions for mutations to be considered as clonal to each metastasis. The minimum VAF cutoff was determined by a simple *k*-means clustering to separate low VAF mutations (less likely to be clonal) from high VAF mutations (more likely to be clonal). The use of hard cutoff parameters inevitably causes false negative or false positive mutation detection. We then assessed the VAF distribution of truncal mutations (defined as those passing the minimum VAF cutoff in 9 or more tumor samples) to verify that most truncal mutations (clonal by definition) pass the cutoff and derive the false negative detection rate. **(B)** Histograms showing the number of somatic mutations passing the minimum VAF cutoffs in a certain number of tumor samples. We operationally chose truncal mutations as those present in 9 or more samples; this threshold does not substantially affect the results of phylogenetic inference as most truncal mutations pass the cutoff in nearly all (13 or 14) samples, either when restricted to disomic chromosomes (for which there is no DNA loss and therefore all mutations should have been preserved) or across the entire genome. **(C)** Validation of the VAF cutoff in each tumor sample based on truncal mutations identified on chr2 (no allelic imbalance in any autopsy sample). While there is a minor DNA copy-number gain of chr2 in the primary tumor, this does not impact the analysis as the primary is not used for phylogenetic inference. **(D)** Phylogenetic inference based on sSNVs detected in the metastatic lesions. The phylogeny was first constructed based on the number of mutations consistent with a binary tree and then refined by assigning each mutation to a branch based on the maximum likelihood calculated from a constant false positive rate (0.01) and sample-specific false negative rate estimated as described in A. The phylogenetic tree was determined based on mutations shared by a subset of biopsies, with mutations assigned to each evolutionary branch labelled with **a-k**. Mutations inferred to be truncal (left) and private (right) were not used for the phylogenetic inference. The final numbers of mutations assigned to each evolutionary branch based on MLE are listed to the right of the phylogenetic tree. Note that the three groups of mutations assigned to branch **b** are almost all from regions on chr1 and chrX that are deleted in the Lung metastases and in RPLN-2; these mutations are classified as ancestral and their absence in the Lung and RPLN-2 genomes represents downstream deletion events in these genomes. Similar patterns of mutations were found on chr4, chr11,and chr22 that are deleted in MedLN/Liver-4, Lung, and RPLN-2/RPLN-3 samples. We manually reviewed mutations with different co-occurrence patterns (at least 5 mutations with the same pattern) to re-assign those that are absent in a subset of tumors due to deletions as truncal. After this manual curation, we identified 15 mutations that are shared between RPLN-2/RPLN-3 and the Liver/RPLN-1/MedLN branch, which suggests they share a more recent common ancestor than with the Lung metastases. We note that the small numbers of mutations shared between RPLN-2/RPLN-3 (8) and shared between the RPLN-2/RPLN-3 and the Liver/RPLN-2/MedLN branches pose the most uncertainty in the phylogenetic tree. These uncertainties, however, do not impact the inference of SCNA evolution as shown in **fig. S9**, as the examples of SCNA evolution were all supported by segmental SCNA breakpoints with high clonality, indicating their late evolutionary timing after the divergence of these branches. **(E)** Phasing of somatic mutations to the deleted chrX and the retained chrX based on the VAFs in all tumor samples. Mutations are ordered by their locations on chrX. Shown for each mutation are the variant allele fractions (graydots) in all tumor samples; outlined dots indicate mutations whose VAF passes the sample-specific VAF cutoff for clonal mutation. Mutations on the retained X are preserved in all tumors (bottom), whereas those on the deleted X have either low or zero VAFs in RPLN-2 and in the Lung metastases, or reduced VAFs in the Liver-4/MedLN samples (where the chromosome loss occurred after whole-genome duplication). The distribution of VAFs of all mutations phased to either chrX is shown on the right for each tumor sample as a boxplot (see also **Fig. 3**). The retained X was further verified to be the active X based on RNA-Sequencing analysis of the primary tumor (**Fig. 4**).

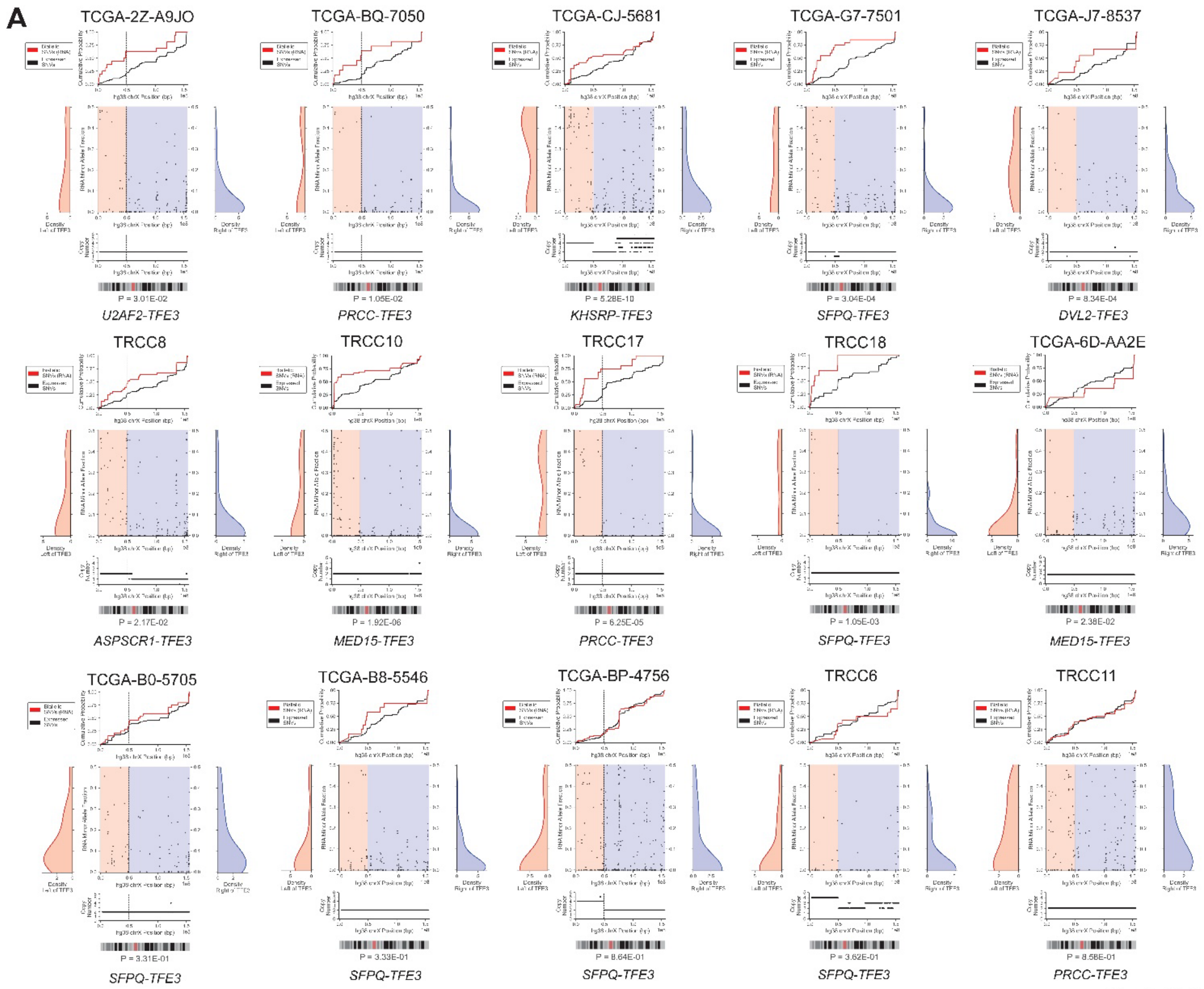
A
TCGA-2Z-A9JO
P = 3.01E-02
U2AF2-TFE3
TCGA-BQ-7050
P = 1.05E-02
PRCC-TFE3
TCGA-CJ-5681
P = 5.28E-10
KHSRP-TFE3
TCGA-G7-7501
P = 3.04E-04
SFPQ-TFE3
TCGA-J7-8537
P = 8.34E-04
DVL2-TFE3
TRCC8
P = 2.17E-02
ASPSCR1-TFE3
TRCC10
P = 1.92E-06
MED15-TFE3
TRCC17
P = 6.25E-05
PRCC-TFE3
TRCC18
P = 1.05E-03
SFPQ-TFE3
TCGA-6D-AA2E
P = 2.38E-02
MED15-TFE3
TCGA-B0-5705
P = 3.31E-01
SFPQ-TFE3
TCGA-B8-5546
P = 3.33E-01
SFPQ-TFE3
TCGA-BP-4756
P = 8.64E-01
SFPQ-TFE3
TRCC6
P = 3.62E-01
SFPQ-TFE3
TRCC11
P = 8.58E-01
PRCC-TFE3

fig. S11-1

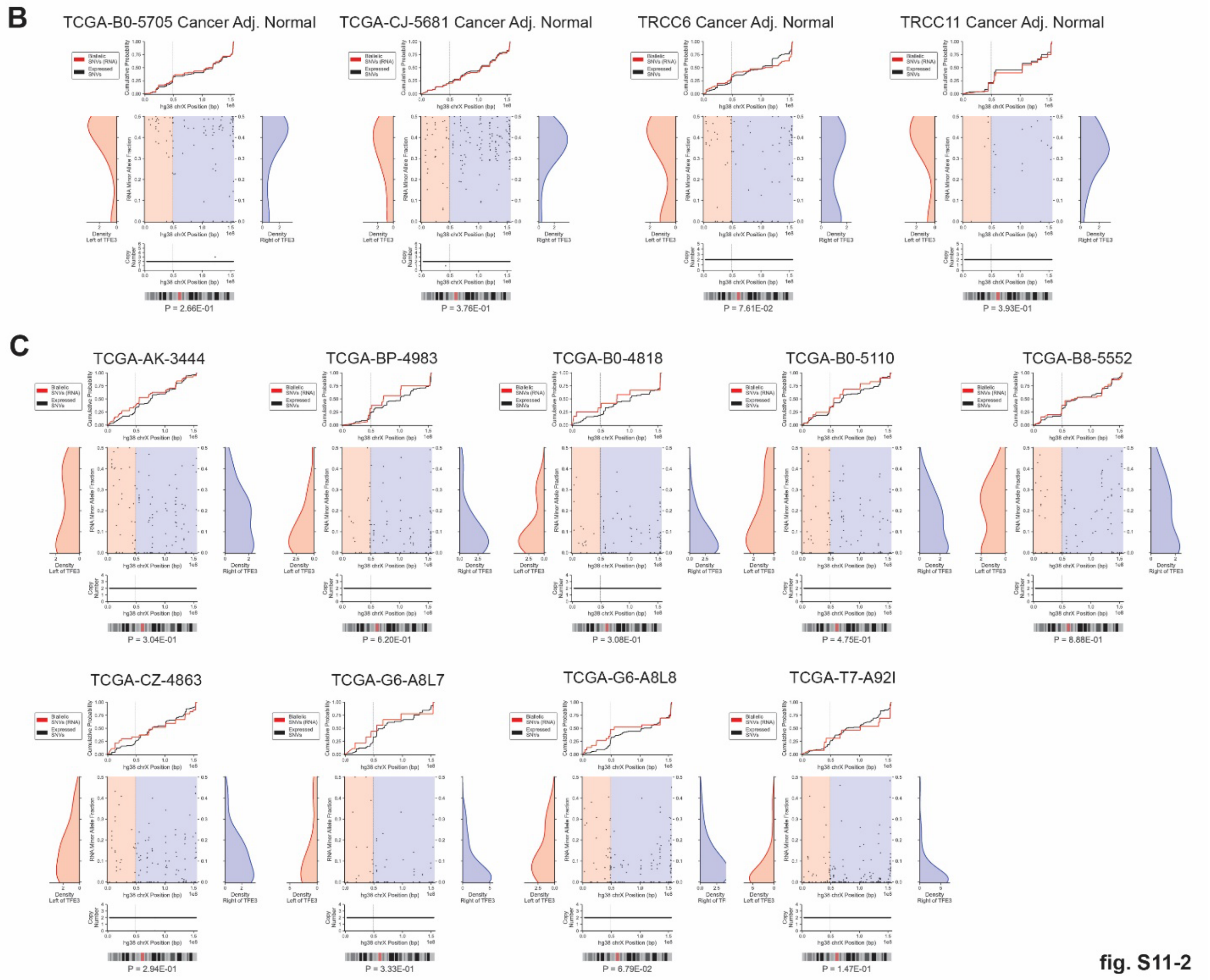


**Fig. S11. chrXi-reactivated samples across tRCC aggregate cohort.** Summary of allelic RNA fraction for chrX at expressed germline heterozygous sites (SNVs, genes escaping X-inactivation excluded) in the following categories of samples: female tRCCs **(A)**; cancer-adjacent normal tissue from female tRCC samples **(B);** female ccRCCs without large copy number alterations on chrX **(C).** In each plot, black line sums the cumulative distribution (beginning at the p-terminus) for germline SNV sites that show RNA expression in the tumor sample (threshold ≥ 10 DNA reads and ≥ 10 RNA reads; see Methods). Red line shows cumulative distribution for expressed germline SNVs that are biallelically expressed (threshold minor allele fraction ≥ 0.2, threshold ≥ 10 DNA reads and ≥ 10 RNA reads). Segmented DNA copy number is also shown across chrX. *P*-values calculated by Mann-Whitney U test between distributions of minor allele fraction for heterozygous sites to the left and right of the *TFE3* breakpoint.

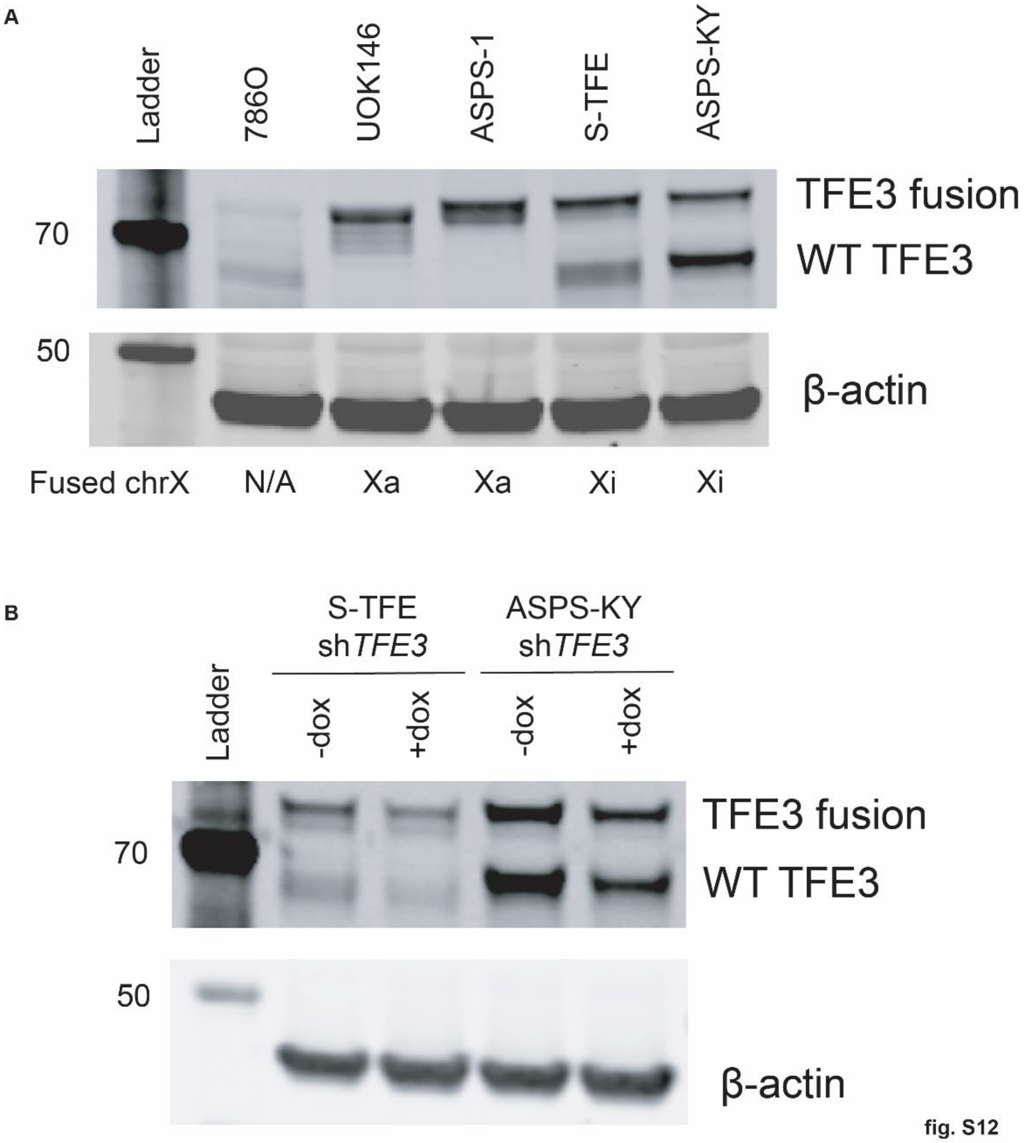


**Fig. S12. Support for chrXi reactivation in in vitro *TFE3* fusion models. (A)** TFE3 western blot across a panel of *TFE3* fusion cell lines (tRCC: UOK146, S-TFE; alveolar soft part sarcoma: ASPS-1, ASPS-KY) and *TFE3* fusion-negative ccRCC cell line 786O. The pattern of Western blot is suggestive of both chrXa and chrXi fusions across this cell line panel (double band: chrXi fusion, single band: chrXa fusion). **(B)** TFE3 western blot with doxycycline-inducible shRNA-mediated knockdown of *TFE3* in ASPS-KY and S-TFE cells with target sequence in the *TFE3* 3'UTR**,** demonstrating that both bands represent TFE3 species.

### Supplementary Table Captions

**Table S1. Sample information, clinical annotations, and sequencing metrics for tRCC WGS cohort.** For each sample in the tRCC WGS cohort, table lists: (A) Sample ID; (B) sex (male/female); (C) sample type (FFPE or frozen); (D) sequencing platform (standard or linked-read WGS); (E) tumor site (primary or metastasis) (F) tumor tissue acquisition method (resection or biopsy); (G) source of normal (tumor-adjacent normal or blood); (H) treatment status at the time of tumor sample acquisition (treatment-naïve or post-treatment); (I) prior therapy (if post-treatment); (J) coverage in tumor; (K) coverage in normal; (L) tumor purity; (M) tumor ploidy.

**Table S2. Summary of somatic variants across tRCC WGS cohort.** For each tRCC sample profiled by WGS, table lists: (A) Sample ID; (B) SNV rate (mutations/Mb); (C) indel rate (indels/Mb); (D) number of SVs of various classes per sample, after SV filters as previously described; (E) identity of *TFE3* fusion

**Table S3. Somatic mutations in cancer driver genes.** Somatic variants in Tier 1 driver genes in the cancer gene census, along with predicted functional effect, in tRCC samples profiled by WGS.

**Table S4. Annotation of *TFE3* genomic breakpoints.** Genomic coordinates for *TFE3* breakpoints across tRCC cohort obtained via manual review, annotated with nucleotides of extent of loss/gain of DNA at break-ends and microhomology (-) or insertion (+) at breakpoints. TFE3 genomic breakpoints detected in WGS by the SVaBA tool and in RNA (where available) by STAR-Fusion are also annotated. Fusions are annotated as arising via intra-chromosomal or autosomal translocation.

**Table S5. Somatic mutations in TRCC18 and their co-occurrence patterns used for phylogenetic inference.**

**Table S6. Primer and sgRNA sequences used for CRISPR/Cas9 engineering of *TFE3* fusions**

**Table S7. Summary of fusion partner, sex, and copy number imbalance status across aggregate tRCC cohort.** Sample ID, dataset, subject sex, copy number imbalance status, and fusion partner annotated for tRCC samples across 11 datasets (this study; TCGA; Sun et al.; Qu et al, Durinck et al., Malouf et al., MSK Panel, MSK WES, OncoPanel, PCAWG, and Sato et al.).
